# Population-scale immunoglobulin genetics resolves the human B-cell system

**DOI:** 10.64898/2026.03.30.715216

**Authors:** Zain Ali, Aitzkoa Lopez de Lapuente Portilla, Gudmar Thorleifsson, Antton Lamarca-Arrizzabalaga, Caterina Cafaro, Gisli H Halldorsson, Ludvig Ekdahl, Mineto Ota, Pall Melsted, Lilja Stefansdottir, Aslaug Jonsdottir, Asgeir Sigurdsson, Erna Ivarsdottir, Keishi Fujio, Stephen J. Harding, Bjorn R Ludviksson, Thorvardur Jon Love, Sigurdur Y Kristinsson, Patrick Sulem, Daniel F Gudbjartsson, Kari Stefansson, Unnur Thorsteinsdottir, Ingileif Jonsdottir, Thorunn Olafsdottir, Björn Nilsson

## Abstract

Immunoglobulins (Ig) mediate adaptive humoral immunity, yet the regulation of B-cell responses *in vivo* in humans remains inaccessible to direct experimentation. Here we use population-scale Ig genetics to resolve molecular regulation of the human B-cell system. Analysis of circulating IgA, IgG, IgM, and six composite Ig traits in 114,697 individuals identifies 504 genetic associations. Integration with regulatory genomics, plasma proteomics, and immunophenotyping maps these effects across the B-cell hierarchy, recovering known regulators and revealing previously unrecognized genes in humoral immunity. At key control nodes – including Fcγ receptors, the immunoglobulin heavy-chain locus and the TACI–APRIL signaling axis – variants form allelic series generating graded perturbations of antibody output. Ig-associated loci show extensive overlap with autoimmunity, immunodeficiency and B-cell malignancy. These findings demonstrate that Ig traits, analyzed at population scale, encode fine-grained information about the regulation of the human B-cell system and link natural variation in humoral immunity to immune-mediated disease.

## INTRODUCTION

Immunoglobulins (Ig) are the effector molecules of adaptive humoral immunity^1^. They arise from a complex differentiation program that begins with B-cell commitment and culminates in the secretion of mature antibodies by plasma cells. Although these processes have been extensively studied in model systems, the *in vivo* regulation of the human B-cell system is inaccessible to experimental perturbation.

Circulating Ig levels provide an integrated readout of B-cell activity and have long served as clinical indicators of humoral immune function. Inter-individual differences in these traits offer an opportunity to probe regulation of the B-cell system through genetic variation. However, the extent to which population variation in Ig traits encodes information about its regulatory architecture remains unclear. Previous genetic studies of Ig traits have been modest in scale (≤19,129 individuals) and have provided only a partial view of the genetic architecture governing antibody production^2-9^.

Here, we analyzed nine Ig traits in 114,697 individuals to resolve the genetic regulation of adaptive humoral immunity. By integrating genetic association signals with regulatory genomics, plasma proteomics, and high-resolution immunophenotyping, we show that Ig traits encode unexpectedly fine-grained information about human B-cell function. Together, these analyses establish circulating Ig traits as quantitatively resolved *in vivo* sensors of B-cell system state and provide a powerful framework for dissecting human humoral immune regulation and its perturbation in immune-mediated disease.

## RESULTS

### Genome-wide association study

To identify DNA sequence variants influencing B-cell physiology, we analyzed IgA, IgG, and IgM levels in 114,697 individuals from Iceland (n = 91,807), Sweden (n = 7,847), and the UK Biobank (n = 13,612; **Figure 1a**; **Table S1A**). To capture additional aspects of humoral immune regulation, we also analyzed six composite traits derived from standardized, log-transformed Ig measurements: AGM (reflecting total Ig levels), AG (class-switched output), M/AG (pre- *versus* post-switch bias), and A/M, G/M, and A/G (isotype-specific class-switching biases; **Table S1B**). These composite traits increase sensitivity to detect genetic effects not apparent in single-isotype analyses^2^.

**Figure 1.**
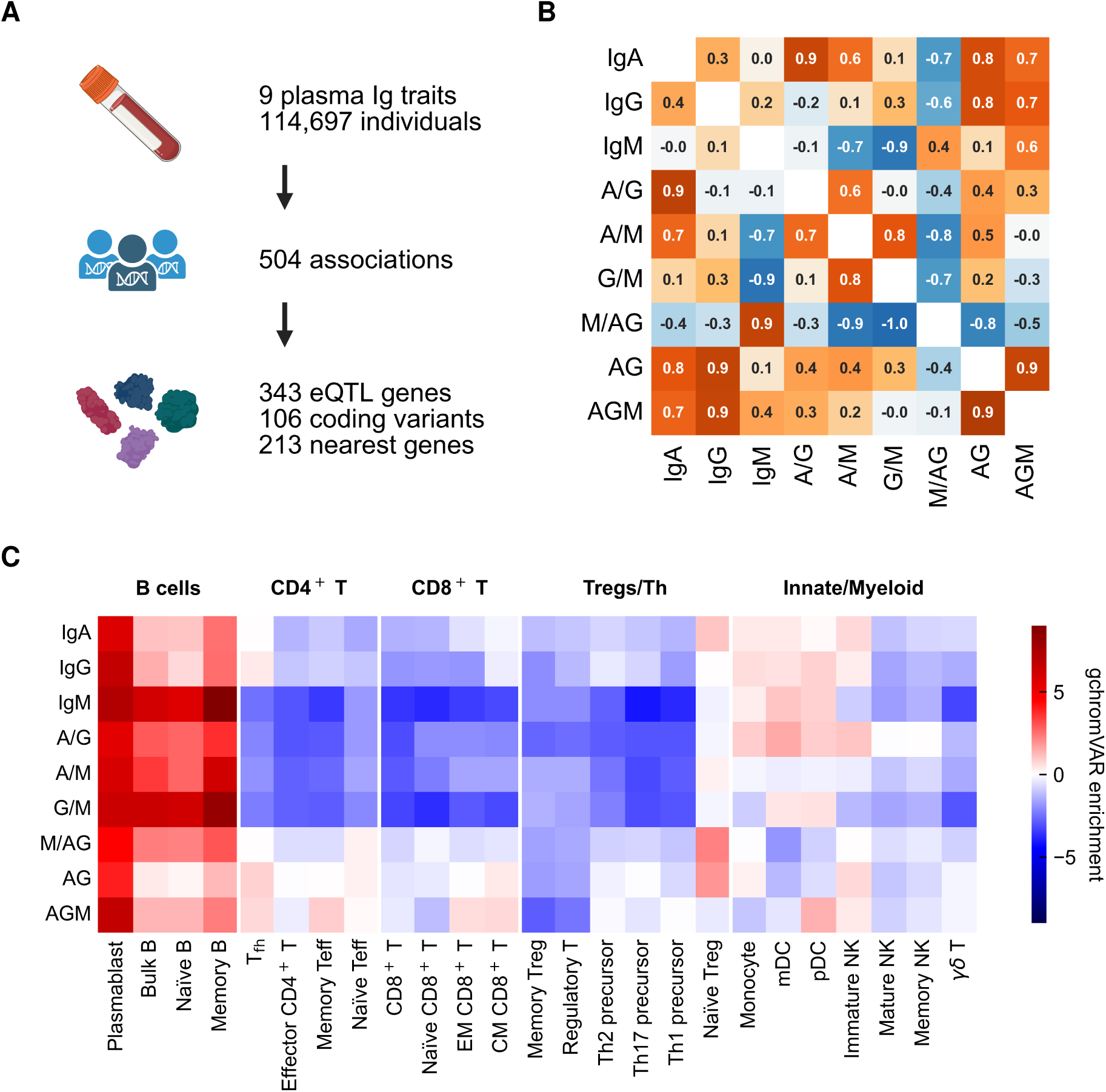
Genetic architecture of circulating immunoglobulin traits across the human B-cell system. **(A)** Study design. Genome-wide association analyses were performed for IgA, IgG, IgM, and six derived composite traits in 114,697 individuals from Iceland, Sweden, and the UK Biobank. Conditional analysis identified 1,144 genome-wide signals, which were resolved to 504 unique variants through 95% credible sets and inter-trait colocalization. **(B)** Enrichment of fine-mapped Ig-associated variants in accessible chromatin regions across immune cell types. Heatmap shows gchromVar enrichment scores for each Ig trait across 26 immune cell populations^69^. **(C)** Pairwise correlations between Ig traits (lower triangle: genetic correlations from linkage disequilibrium score regression^13^; upper triangle: phenotypic correlations). Numbers and color scale represent Pearson correlation coefficients.

We performed association testing against 62.2 million variants (**Methods**). Following multiple-testing correction **(Table S1C**)^10^, conditional analyses identified 1,144 independent signals across the nine Ig traits (**Table S2A**), which were collapsed to 504 unique variants using 95% credible sets and inter-trait colocalization analysis (**Table S2B-D**; **Figure 1A**; **Methods**)^11,12^. Among these, 472 were not reported previously, including 33 rare variants (effect allele frequency, EAF, ≤1%; **Table S2B**, **E**). Lead variants explained 11-21% of trait variance, consistent with heritability estimates of 13-18% by linkage disequilibrium score regression (**Table S2F**)^13^. Fine-mapping resolved 70 signals to single putative causal variants (**Figure S1A**). Most variants were polymorphic across gnomAD ancestries (**Table S2G**). Notably, 186 signals detected in composite traits were not captured by any single-isotype analysis **(Figure S1B**). We next investigated the biological mechanisms through which these variants influence humoral immunity.

### Functional annotation

#### Prioritization of candidate genes at Ig-associated loci

To prioritize candidate effector genes at Ig-associated loci, we integrated: (i) non-synonymous coding variants in credible sets (posterior inclusion probability > 10%; **Table S3A**); and (ii) *cis*-eQTLs in immune cells and blood (colocalization posterior probability > 90%; **Table S3B**)^14,15^. This conservative approach implicated 397 genes across 290 signals. For loci lacking molecularly supported candidates, we assigned the nearest protein-coding gene to the lead variant (**Table S2B**). Candidate genes were enriched for Ig-related mouse phenotypes and Reactome terms (**Tables S3C-D**)^16-18^. In RNA-sequencing data for 81 cell types^19^, these genes were most highly expressed in B cells and professional antigen-presenting cells (**Table S3F**). Implicated genes span all key stages of Ig biology (**Figure 2**), including B-cell transcription factors (*EBF1*, *NFKB1*, *IKZF1/3, IRF4, PAX5*, *RUNX3*)^20-23^; cytokines and their receptors (*TNFSF13*/APRIL, *TNFRSF13B*/TACI, *TNFRSF17*/BCMA, *TNFSF13B*/BAFF)^24-26^; plasma cell effectors (*ATG5*, *ELL2, SLAMF7*)^27-29^; and Ig effector and clearance components (*FCGR2A/B*, *FCGR3A/B*, *FCGRT*)^30,31^. Consistently, Ig-associated variants were enriched in B-cell open chromatin (permutation test, *P =* 7.9×10^-5^), indicating that Ig variation is predominantly mediated through B-cell-autonomous regulatory elements (**Figure 1C**). Together, these results indicate that Ig-associated loci resolve into a coherent regulatory network spanning the B-cell life cycle.

**Figure 2.**
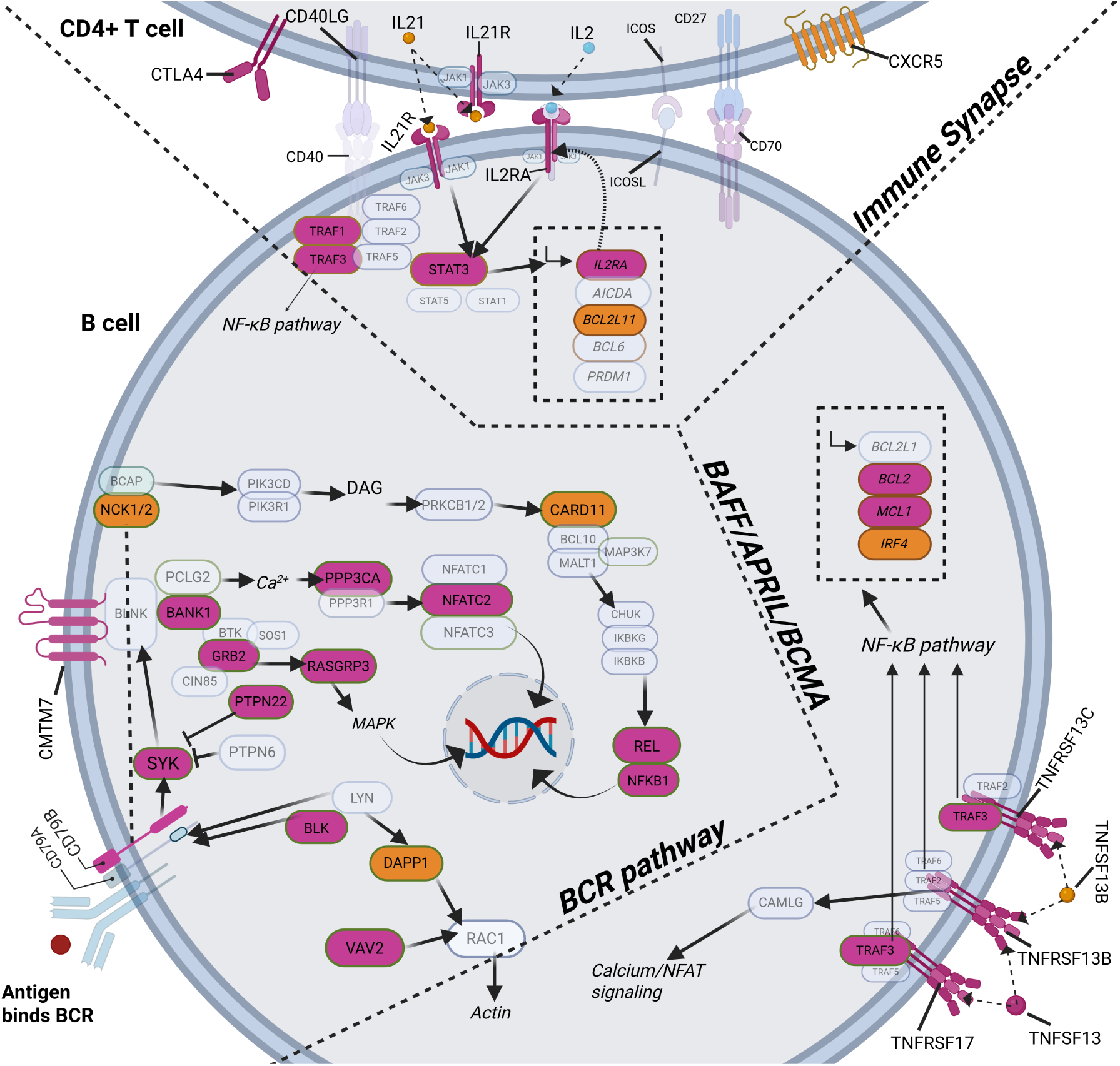
Ig-associated variants converge on core regulatory pathways of the human B-cell system. Candidate genes identified in the Ig genome-wide association study span all major stages of B-cell biology, including transcriptional regulation, cytokine signaling, plasma-cell function, and immunoglobulin effector and clearance pathways. The pathway diagram highlights three central B-cell programs recovered in our analyses: B-cell receptor (BCR) signaling, the BAFF–APRIL–BCMA cytokine axis, and the immune synapse between B cells and helper T cells. Genes shown in purple were supported by coding variants or colocalizing cis-expression quantitative trait loci (eQTLs), whereas genes in orange were assigned by proximity. Transparent nodes denote pathway components not implicated in this study. Representative components include BCR signaling molecules (SYK, BLK, BLNK, and BANK1), downstream transcriptional effectors (NFAT and NF-κB pathways), T-cell costimulatory molecules (CD40 and IL21R), and BAFF/APRIL cytokines and their receptors (TNFRSF13B/TACI, TNFRSF17/BCMA, and TNFRSF13C/BAFFR). Pathway topology was reconstructed from Reactoeactome^17^.

#### Ig genetic architecture recapitulates B-cell differentiation hierarchies

To determine how genetic variation partitions across antibody classes, we compared genetic signals across Ig traits to delineate shared and isotype-specific mechanisms. IgM showed minimal genetic correlation with IgA and IgG (*r*_g_ =-0. 0.012 and 0.13, respectively), consistent with largely distinct networks governing pre-switched *versus* class-switched antibody production. In contrast, IgA and IgG were more correlated (*r*_g_ = 0.36, *P =* 4.9×10^-11^), reflecting shared pathways in class-switched B-cell populations (**Figure 1B**; **Table S4**).

Twenty-three candidate genes (5.8%) were shared across all three isotypes (**Table S2B**; **Figure S2**), including core B-cell transcription factors (*IKZF1, IKZF3*, and *IRF4*), signaling regulators (*TNFRSF13B/*TACI and *TNFSF13/*APRIL), and survival genes (*BCL2*)^32^. By contrast, isotype-specific loci reflect specialized functions: IgA-exclusive loci support mucosal immunity (*e.g.*, *LTBR* and *INAVA*)^33-36^; IgG loci mirror effector function (*e.g.*, Fcγ receptors and Fcγ-glycosylating enzymes such as *ST6GAL1*)^30,31,37^; IgM loci regulate early B-cell development (*e.g.*, *PAX5*)^38^. Together, these observations indicate that Ig-associated variants are organized along the B-cell differentiation hierarchy, with distinct genetic architectures underpinning each isotype.

#### Disease convergence of Ig-associated loci

Ig-associated loci show extensive convergence with the genetic architecture of immunological diseases. Cross-referencing with the GWAS Catalog and ClinVar revealed that 148 Ig-associated variants overlapped risk variants for 141 disease outcomes (*r*^2^ > 0.8 between Ig and catalog lead variants; **Table S11A**; **Figure S3**)^39,40^. Overlap was most pronounced for autoimmune and allergic conditions (n = 99 variants), B-cell malignancies (n = 34), and immunodeficiencies or infection susceptibility (n = 17). Consistent with this convergence, OMIM^41^ implicated 76 candidate genes in heritable immunodeficiencies or developmental syndromes with immunological features (**Table S11B**). The IntOGen^42^ and Mitelman databases^43^ identified 44 candidate genes as somatic drivers enriched in B-cell lymphomas and multiple myeloma (**Table S11C**). In line with these enrichments, 111 candidate genes were essential in B-cell cancer cell lines in DepMap (**Table S11D**)^44^.

Among rare disease variants, three independent alleles at *IRF4-EXOC2* and *GLRX5* that markedly increase IgM have been linked to Waldenström’s macroglobulinemia (**Figure S1C**)^45-47^, consistent with the monoclonal IgM production that defines this disorder. Two loss-of-function variants in *TNFRSF13B* (TACI) that reduce total Ig are known risk factors for common variable immunodeficiency (**Figure S1D**)^48,49^. We also identified a missense variant in *DOCK2*, an established severe immunodeficiency gene (rs761943047-C, p.Val406Leu; AlphaMissense = 0.61, CADD = 26.7; **Figure S1D**)^50,51^.

Collectively, Ig-associated loci show broad and biologically coherent convergence with immune disorders, particularly within the B-cell lineage.

#### Proteomic signatures reveal mediators of Ig variation

To define molecular mediators of Ig-associated variants, we next integrated our association signals with plasma proteomics data from 54,265 UK Biobank participants^52-54^. A total of 237 Ig-associated variants correlated (*r*^2^ > 0.8) with 1,354 *trans*- and 59 *cis*-pQTLs, influencing 730 distinct proteins (**Table S5A**). Twelve proteins show Bonferroni-significant enrichment of one or more Ig traits (**Table S5B**), revealing recurrent mechanistic pathways.

The strongest signal emerged for IgM-associated loci, which overlapped strongly with J-chain (30 variants; Binomial test *P =* 1.1×10^-8^) and CD5L (23 variants; *P =* 6.7×10^-5^; **Table S5B**), with a near-perfect concordance between IgM and pQTL effects across all mappable IgM-associated variants (J-chain: inverse variance-weighted, IVW, regression β = 0.62, *P =* 3.6×10^-193^; CD5L: β = 0.75, *P =* 2.7×10^-211^; **Figure S4A, Table S5C**). J-chain polymerizes IgM monomers into pentamers, while CD5L associates with IgM via J-chain^55,56^. These stoichiometric relationships provide validation of IgM associations through an independent readout.

A second pattern was observed for IgA-associated loci, which were enriched for LY96 *trans*-pQTLs (20 variants; Binomial test *P =* 9.6×10^-9^; **Table S5B**), a secreted TLR4 co-receptor that is essential for bacterial lipopolysaccharide recognition^57^, with similarly strong concordance across IgA-associated variants (IVW regression β = 0.68, *P =* 2.2×10^-212^, **Figure S4B, Table S5C**). Although canonically myeloid-expressed, LY96 is also expressed in plasma cells and is upregulated in *ex vivo* stimulated B cells (**Figure S4D-E**). LY96 has no known physical association with IgA, but the strength of this association, comparable to that of stoichiometric IgM subunits, indicates a previously unrecognized molecular relationship.

Finally, loci associated with class-switched Ig levels (IgA, IgG, and AG) were enriched for *trans*-pQTL effects on proteins expressed on mature B and plasma cells, including FCRL5, IL5RA, MZB1, and TNFRSF17/BCMA (**Table S5B-C**; **Figure S4C-D**). The effect size concordance was highly significant (IVW regression β = 0.44 to 0.68, *P =* 10^-13^ to 10^-59^) but weaker than the IgM–Jchain/CD5L and IgA–LY96 correlations, consistent with variation in activity of mature B and plasma cells rather than direct molecular coupling. Notably, FCRL5 and BCMA are targets of multiple myeloma immunotherapy, and plasma IL5RA and BCMA levels are elevated in carriers of multiple myeloma risk alleles^58^.

Taken together, these results demonstrate that *trans*-pQTL analysis can recover known stoichiometric subunits of pentameric IgM with high fidelity, lending confidence to the unexpected finding that circulating IgA and LY96 are coupled at a comparable scale. Additionally, the class-switched Ig *trans*-pQTL signatures implicate genetic variation in mature B- and plasma-cell activity as a central axis of humoral immune regulation.

#### Cellular mechanisms of genetic regulation of circulating immunoglobulins

To determine whether Ig-associated variants act through specific immune cell populations, we leveraged the BloodVariome compendium, a high-resolution atlas of inherited effects on human immune cell states (1,005 flow cytometry traits across 127 immune cell subsets in 11,983 individuals; Lopez de Lapuente Portilla *et al.*, companion manuscript). Twenty-two Ig-associated variants colocalized with at least one immune cell trait (posterior probability > 90%; **Table S6**; **Figure 3, S5**). Of these, 12 affected B-cell traits, indicating B-cell-intrinsic effects (**Figure S5A-L**), whereas the remaining 10 affected non-B-cell traits, consistent with indirect mechanisms of Ig regulation, primarily via T helper cells (**Figure S5M-O**), regulatory T cells (**Figure S5P-Q**), and dendritic cells (**Figure S5R-V**).

**Figure 3.**
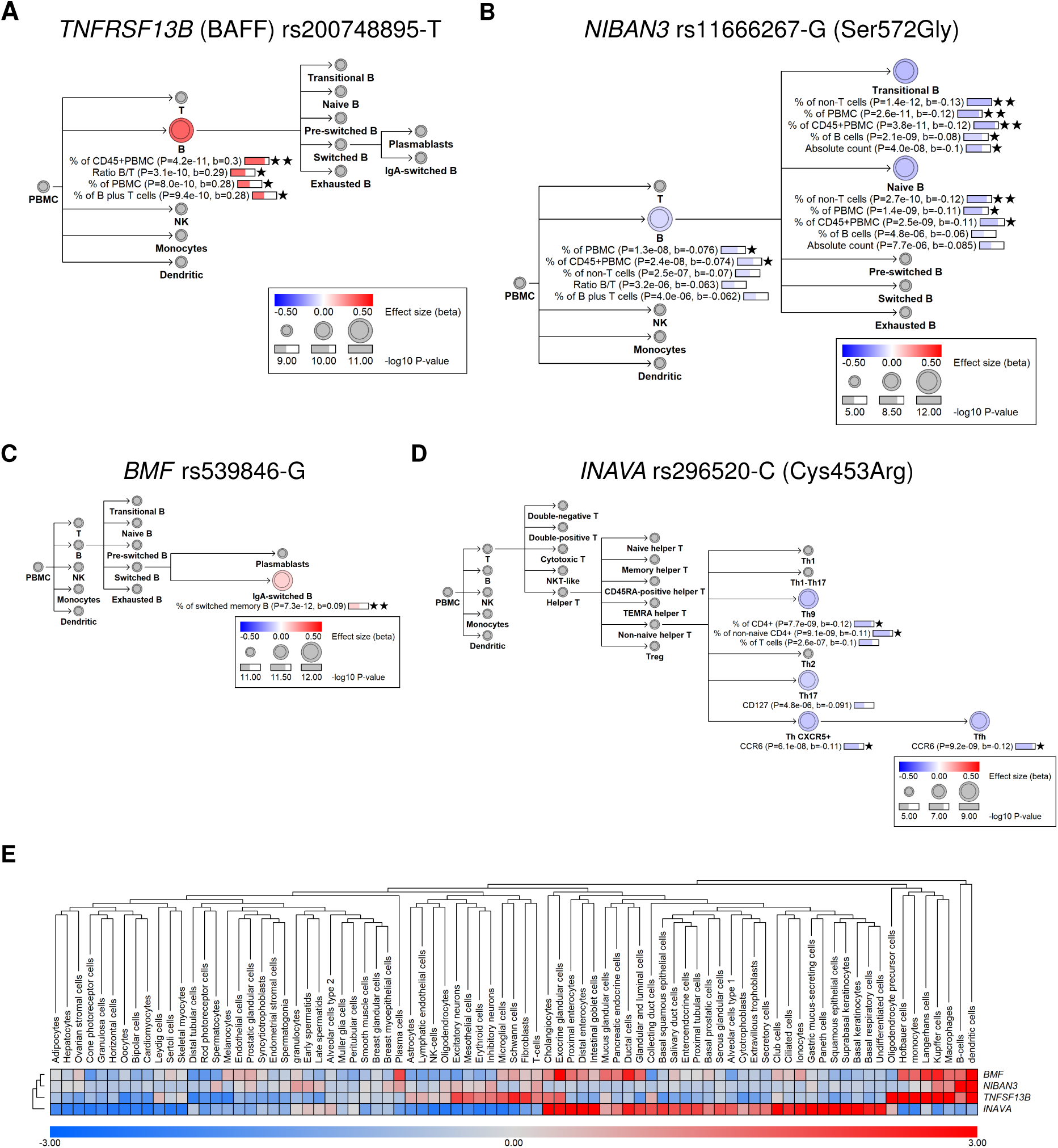
Genetic colocalization with BloodVariome resolves cellular mechanisms of Ig regulation. To define the cellular basis of Ig-associated variants, we integrated our results with the BloodVariome GWAS dataset (1,005 flow cytometry traits across 127 immune cell subsets in 11,983 individuals; companion manuscript). We identified 22 colocalized associations, including 12 with direct effects on B-cell traits and 10 consistent with indirect regulation via helper T cells, regulatory T cells, or dendritic cells. Visualizations of the effects of all colocalized variants are provided in **Figure S4**. **(A)** Immune cell effects of the 3′-UTR variant rs374039502-A in *TNFSF13B* (BAFF). The variant increases total Ig levels and B-cell frequencies, consistent with BAFF-driven B-cell survival. **(B)** Missense variant in *NIBAN3* (rs11666267-G; p.Ser572Gly) reduces IgM levels and frequencies of naïve and transitional B cells, genetically implicating *NIBAN3* in early human B-cell development. **(C)** Variant rs539846-G at *BMF* elevates IgA levels and IgA-switched B cells while downregulating BMF expression, consistent with its pro-apoptotic role. **(D)** Example of indirect regulation: missense variant rs296520-C in *INAVA* (p.Cys453Arg) reduces IgA and the IgA/IgG ratio while decreasing CCR6 expression on follicular helper T (Tfh) cells, as well as Th9 and Th17 frequencies, linking mucosal barrier pathways to T cell–dependent control of IgA production. **(E)** Expression profiles across 81 human cell types^103^ showing predominant immune expression of *TNFRSF13B*, *NIBAN3*, and *BMF*, and epithelial enrichment of *INAVA*.

Ig-associated variants with B-cell–intrinsic effects resolve discrete regulatory control points along the B-cell differentiation axis. The low-frequency variant rs374039502-A in the 3’-UTR of *TNFSF13B* (BAFF) increases total Ig (β_AGM_ = 0.22, *P*_AGM_ = 3.2×10^-17^), B-cell counts (β = 0.302, *P =* 4.2×10^-11^), and plasma BAFF levels (β = 0.79, *P =* 3.1×10^-246^), consistent with BAFF-driven B-cell expansion (**Figure 3A, S5A**)^59^. The variant also increases the risk of tonsillectomy (OR = 1.26; **Table S11A**), linking elevated BAFF levels to tonsillar hyperplasia. Similarly, the *NIBAN3* missense variant rs11666267-G (p.Ser572Gly; CADD = 22.0) reduces IgM (β_IgM_ = -0.03, *P*_IgM_ *=* 6.3×10^-9^) and lowers total, transitional, and naïve B cell frequencies (β = -0.13 to -0.074, *P =* 1.4×10^-12^ to 2.4×10^-8^; **Figure 3B**, **S5B**).

*NIBAN3* encodes a membrane protein expressed in B cells^60^. Knockout mice show increased B-cell proliferation and IgM/IgG3 production^61,62^, whereas overexpression enhances apoptosis, together supporting rs11666267-G as a gain-of-function allele. Further, at *BMF*, rs539846-G increases IgA (β_IgA_ = 0.04, P_IgA_ = 1.1×10^-15^) and IgA-switched memory B cells (β = 0.09, *P =* 1.2×10^-11^) while downregulating *BMF* in switched memory B cells (**Figure 3C**, **S5C, S6**), consistent with BMF encoding a pro-apoptotic factor whose reduced expression may promote survival of class-switched B cells^63,64^.

Among extrinsic signals, the missense variant rs296520-C at *INAVA* (p.Cys453Arg) reduces A/G (β = -0.08, *P =* 3.5×10^-47^**; Table S2B**) and decreases CCR6 surface expression on follicular helper T (Tfh) cells (**Figure 3D**, **S5M**). *INAVA* is highly expressed in intestinal epithelial cells and T cell subsets (**Figure 3E**), where it functions as a scaffold protein that amplifies NOD2- and Toll-like receptor-mediated signaling and promotes mucosal immunity. Genetic variation at *INAVA* has been linked to inflammatory bowel disease^35,36^, and *Ccr6*^-/-^mice exhibit reduced IgA production^65^. These observations support a model in which rs296520-C influences IgA by modulating Tfh function (**Figure S5M**)^66-68^.

Collectively, these findings show that Ig-associated variants act across both the B-cell lineage and upstream immune circuits.

### Mechanistic interrogation of key Ig loci

Having systematically annotated Ig-associated variants across regulatory, proteomic, and cellular layers, we next performed focused mechanistic analyses of selected loci. These include both novel B cell regulators and well-characterized hubs of humoral immunity – specifically the immunoglobulin heavy-chain (*IGH*) locus, Fcγ receptors, *TNFRSF13B* (TACI), and *TNFSF13* (APRIL) – where we uncover previously unrecognized functional variation and regulatory mechanisms.

#### Novel B-cell regulators

Beyond loci with established roles in B-cell biology, we identified multiple genes not previously implicated in humoral immunity (**Table S2B**). We highlight four loci where convergent molecular evidence uncovers novel regulators of B-cell function.

Two independent variants upstream of *ZNF608* reduce IgM (rs12517864-A, β = -0.10, *P =* 5.5×10^-68^; rs4279337-A, β = -0.065, *P =* 2.4×10^-26^; **Figure 4A-B**). rs12517864-A colocalizes with increased *ZNF608* expression in B cell subsets (**Figure S7**) and exhibits allelic imbalance in chromatin accessibility in heterozygous primary B cells (A allele fraction = 0.92, binomial test *P =* 2.14×10^-14^; **Figure 4C**)^69^, supporting causality. The variant overlaps a SPI1 binding site, and motif analysis predicts increased SPI1 affinity concordant with *ZNF608* upregulation (**Figure 4D**). *ZNF608* is recurrently mutated in B-cell lymphomas^70,71^ and is preferentially expressed in germinal center (GC) B cells, parallelling the GC master regulator *BCL6* (**Figure 4E**)^72^.

**Figure 4.**
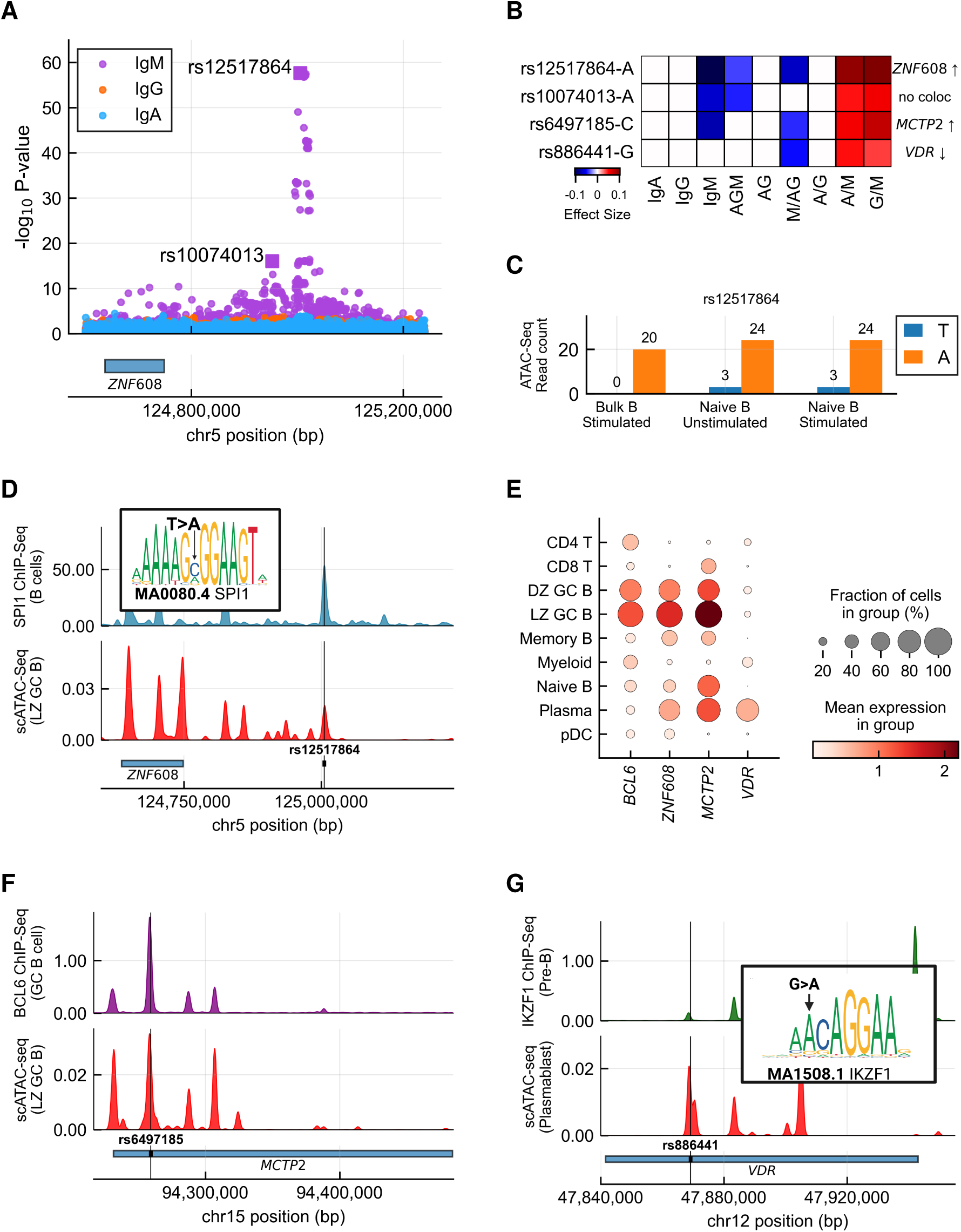
Human genetics identifies novel regulatory nodes in the B-cell differentiation program. Beyond established regulators, multiple Ig-associated loci nominate genes not previously linked to B-cell biology. **(A)** Regional association plot at the *ZNF608* locus showing summary statistics for IgA (cyan), IgG (orange), and IgM (magenta). Two independent signals are observed (rs12517864-A and rs4279337-A). **(B)** Effect-size heatmap for four association signals implicating three novel genes: *ZNF608*, *MCTP2*, and *VDR*. All four variants influence IgM or IgM-derived ratio traits. Colocalizing eQTL genes and directional effects are annotated. **(C)** Allelic imbalance at rs12517864 across three B-cell subsets from a heterozygous donor^69^. Strong allelic skew toward the A allele, associated with reduced IgM, is observed in stimulated bulk B cells, unstimulated naïve B cells, and stimulated naïve B cells (data from Gene Expression Omnibus, accession no. GSM3686919^104^). **(D)** Regulatory landscape of the *ZNF608* locus showing SPI1 ChIP–seq signal in primary B cells and single-cell ATAC–seq accessibility in light-zone germinal center B cells. Variant rs12517864 overlaps a SPI1 binding site within accessible chromatin. Inset: JASPAR SPI1 motif predicts increased binding affinity of the A allele, consistent with increased *ZNF608* expression. **(E)** Single-cell RNA-sequencing expression patterns in human tonsils^105^ for *BCL6*, *ZNF608*, *MCTP2*, and *VDR*. *BCL6*, *ZNF608*, and *MCTP2* show concordant enrichment in germinal center B cells (dark and light zones), whereas *VDR* is preferentially upregulated in plasmablasts, indicating activity at later differentiation stages. **(F)** Regulatory landscape of the MCTP2 locus showing BCL6 ChIP–seq signal in germinal center B cells (Gene Expression Omnibus, accession no. GSM1668937)^75^ and single-cell ATAC–seq accessibility in light-zone germinal center B cells^105^. rs6497185-C overlaps a BCL6 binding site and is associated with increased *MCTP2* expression in unswitched memory B cells. **(G)** Regulatory landscape of the VDR locus showing IKZF1 ChIP–seq signal in a pre-B-cell line (Gene Expression Omnibus, accession no. GSM2882824)^106^ and single-cell ATAC–seq accessibility in plasmablasts^105^. Variant rs886441-G overlaps an IKZF1 binding site and is associated with reduced VDR expression. Inset: JASPAR IKZF1 motif (accession no. MA1508.1) predicts reduced binding affinity of the G allele.

A germinal center signature is also evident at a second locus. rs6497185-C reduces IgM (β = -0.07, *P =* 6.1×10^-38^) and upregulates the calcium-dependent signaling regulator *MCTP2* in unswitched memory B cells (**Figure 4B, S7**)^73,74^. Like *ZNF608*, *MCTP2* is enriched in GC B cells, and the variant overlaps an accessible BCL6 binding site (**Figure 4F**)^75^. Together, these data implicate *ZNF608* and *MCTP2* as candidate components within the GC transcriptional network governed by *BCL6* and *SPI1*^72,76^, At a third locus, rs886441-G decreases M/AG (β = -0.049, *P =* 1.7×10^-12^) and downregulates the vitamin D receptor gene *VDR* in plasmablasts (**Figure 4B, S7C**). Motif analysis predicts reduced IKZF1 binding at this element, consistent with lower *VDR* expression (**Figure 4G**). *VDR* is strongly upregulated in tonsillar plasmablasts **(Figure 4E**), and rs886441 intersects a regulatory element co-bound by IRF4 and PRDM1 during plasmablast differentiation (**Figure S7D**)^77^, positioning *VDR* within transcriptional circuitry governing plasma cell commitment^78^. These data support an inhibitory role for vitamin D signaling in class-switched Ig production^79^. Although vitamin D deficiency has been linked to autoimmunity, effects have largely been attributed to T cells^80-83^. Our findings provide human genetic evidence that VDR signaling in plasmablasts modulates Ig production^84,85^.

Finally, at *ARHGAP15*, both a common and a rare variant reduce IgA (rs79716587-A and rs140397066-G; EAF = 13.2% and 0.7%; β = -0.043 and -0.17, *P =* 6.8×10^-10^ and 4.0×10^-11^; **Figure S8**; **Table S2B**). The rare variant colocalizes with reduced *ARHGAP15* expression in B cells (**Figure S8A-B**), implicating this Rac GTPase–activating protein as a novel regulator of B cell signaling.

To date, most genes regulating B-cell physiology have been identified in model organisms. Our results demonstrate that population-scale Ig genetics can directly nominate genes, cell types, and regulatory mechanisms involved in humoral immunity in humans.

#### Composite ratio traits resolve class-switch regulation at the IGH locus

We next turned to major canonical hubs of humoral regulation. The *IGH* locus on chromosome 14q32.33 encodes immunoglobulin heavy-chain genes that undergo somatic recombination to generate antibodies of distinct isotypes^86^. Due to its complexity, the regulatory architecture of *IGH* has been difficult to resolve through human genetic studies.

We identified 17 independent signals at *IGH*, including 12 not previously reported (**Figure 5A**; **Table S7**)^2,87^. These variants showed 2.5-fold enrichment in the constant region (permutation test *P =* 2.7×10^-3^), with five of 17 lead variants mapping to the 3’RR1 super-enhancer (OR = 19.9; Fisher’s exact *P =* 2.8×10^-5^), a central regulatory hub for class-switch recombination^88^. Consistent with this architecture, most *IGH* signals (15/17; 88%) showed their strongest association with composite ratio traits capturing class-switching dynamics^2^.

**Figure 5.**
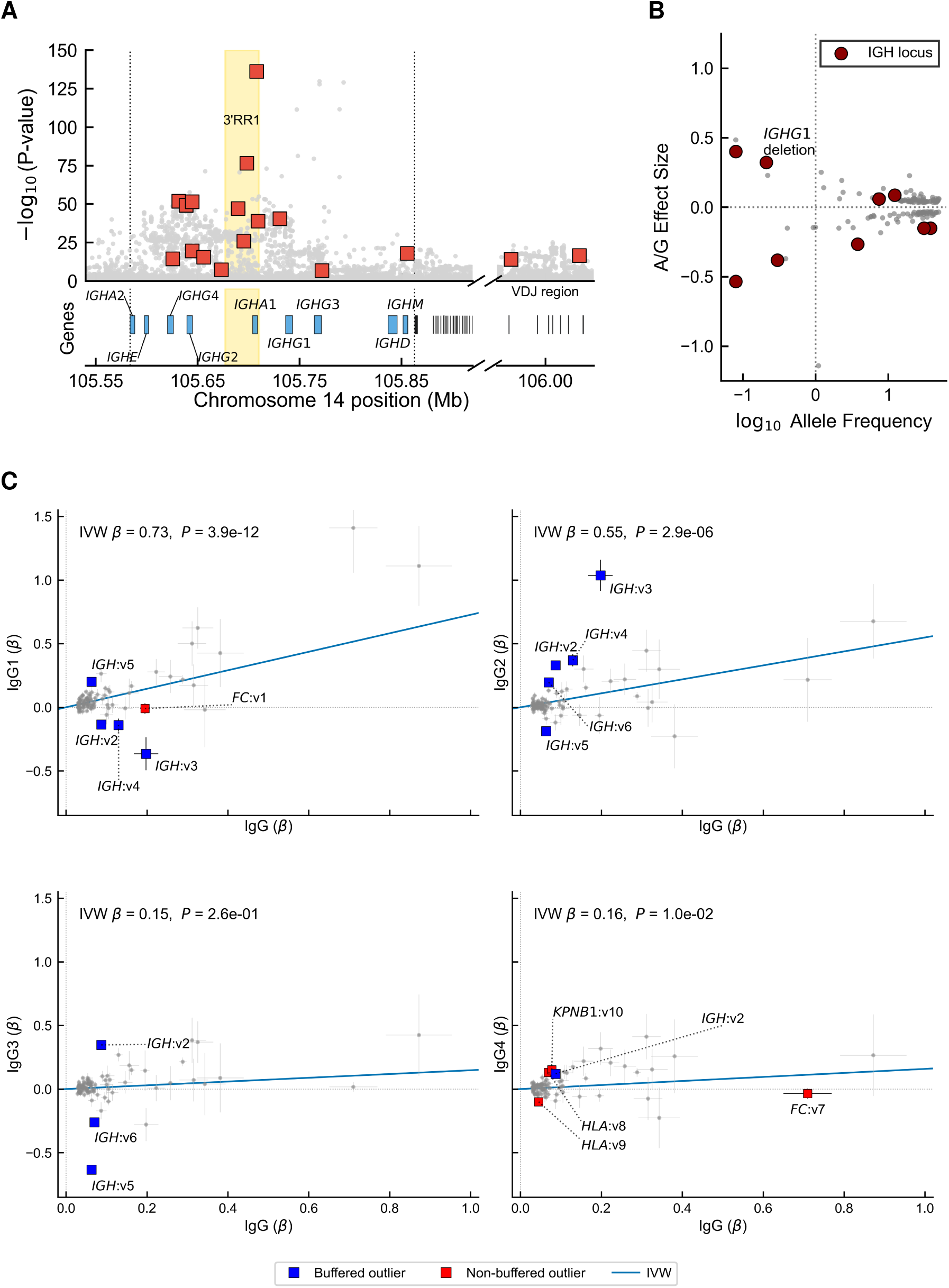
The *IGH* locus encodes intrinsic buffering of IgG subclass output. **(A)** Association landscape across the *IGH* locus on chromosome 14q32.33 showing the minimum P value across nine Ig traits for each variant. Seventeen independent signals were identified (v1–v17; **Table S7**). Associations are enriched within the constant region that mediates class-switch recombination and further concentrated at the 3′ regulatory region 1 (3′RR1), a super-enhancer critical for isotype selection. **(B)** Effect size *versus* effect allele frequency for A/G-associated variants. Variants at the *IGH* locus are highlighted, including the *IGHG1* deletion tagged by rs587597004. **(C)** Comparison of total IgG effects with IgG1–4 subclass effects across 89 IgG-associated variants that could be mapped to our prior IgG subclass GWAS (**Table S8A**)^87^. The solid blue lines represent weighted regression through the origin. Outlier variants (large squares) were classified as buffered (blue) or non-buffered (red). Five buffered outliers were identified, all mapping to the IGH locus (labelled v2–v6 for brevity in the panels; rsIDs are provided in **Table S8B**). For example, rs191766497-T (v3), a splice variant causing IgG2 deficiency, is accompanied by compensatory increases in IgG1, IgG3, and IgG4. Additional outliers were observed at loci outside the *IGH* region (v1, v7, v8, v9, v10). However, these did not show buffering behavior. Together, these results support IgG subclass compensation as an intrinsic property of the *IGH* locus.

Four rare A/G-associated variants exhibited particularly large effects (**Figure 5B**; **Table S7**), including rs587597004-T, which tags a 28-kb deletion spanning *IGHG1* that causes IgG1 deficiency^87^, highlighting the sensitivity of composite ratio traits for resolving high-impact perturbations of class switching. Notably, only two of the 17 signals contain coding variants in their credible sets (**Table S2C**, **S7**), indicating that the majority of the *IGH* variants are mediated through non-coding *cis*-regulation mechanisms. Together, these findings demonstrate that composite ratio traits substantially enhance resolution of human class-switch control and reveal a predominantly *cis*-regulatory architecture at *IGH*.

#### IgG subclass buffering is an intrinsic property of the IGH locus

The four IgG subclasses (IgG1-4) are encoded by distinct constant region genes within *IGH*. In a recent genome-wide association study on IgG1-4 subclass levels in 4,334 individuals^87^, we observed that an *IGHG1* deletion tagged by rs587597004-T reduced IgG1 while increasing IgG3, thereby buffering total IgG levels. This finding suggested mechanisms that preserve overall IgG output despite perturbations in IgG subclasses.

Leveraging the larger sample size of the present study, we evaluated 89 IgG-associated variants for evidence of subclass buffering using the IgG1-4 dataset (**Table S8A**)^87^. Using an outlier-based regression framework, we distinguished variants with proportional subclass effects from those showing reciprocal deviations consistent with compensatory regulation (**Methods**). Five IgG-associated variants showed evidence of buffering, all mapping to *IGH* (**Figure 5B**, **Table S8B**). For example, rs191766497-T, an *IGHG2* splice variant causing familial IgG2 deficiency, markedly reduced IgG2 (β = -1.04, *P =* 1.5×10^-17^) while increasing IgG1 (β = 0.37, *P =* 4.7×10^-3^)^87^. The remaining *IGH* outliers showed similar reciprocal shifts. Notably, no buffering signals were detected outside *IGH*, indicating that subclass compensation is a locus-intrinsic property rather than a general feature of IgG regulation.

Together, these results identify subclass buffering as an intrinsic systems property of the *IGH* locus that stabilizes total IgG output in the face of subclass-specific genetic perturbation. Defining the molecular basis of this buffering architecture will require larger genetic studies of IgG subclasses and functional dissection of *IGH* regulatory elements.

#### IgG traits resolve functional variation across the Fcγ receptor locus

The Fcγ receptor (FcγR) locus on chromosome 1q23 encodes activating and inhibitory receptors that bind IgG to mediate immune-complex clearance, cytotoxic effector function, and inhibitory feedback. We identified 24 independent signals at this locus (20 novel), of which 23 influence IgG or IgG-derived composite traits, including the four rare variants with the largest effects on IgG in our study (**Figure 6A–C**; **Table S9**). These signals affect multiple Fcγ receptor genes (**Tables S3B** and **S9**). We highlight four examples illustrating distinct mechanisms:

**Figure 6.**
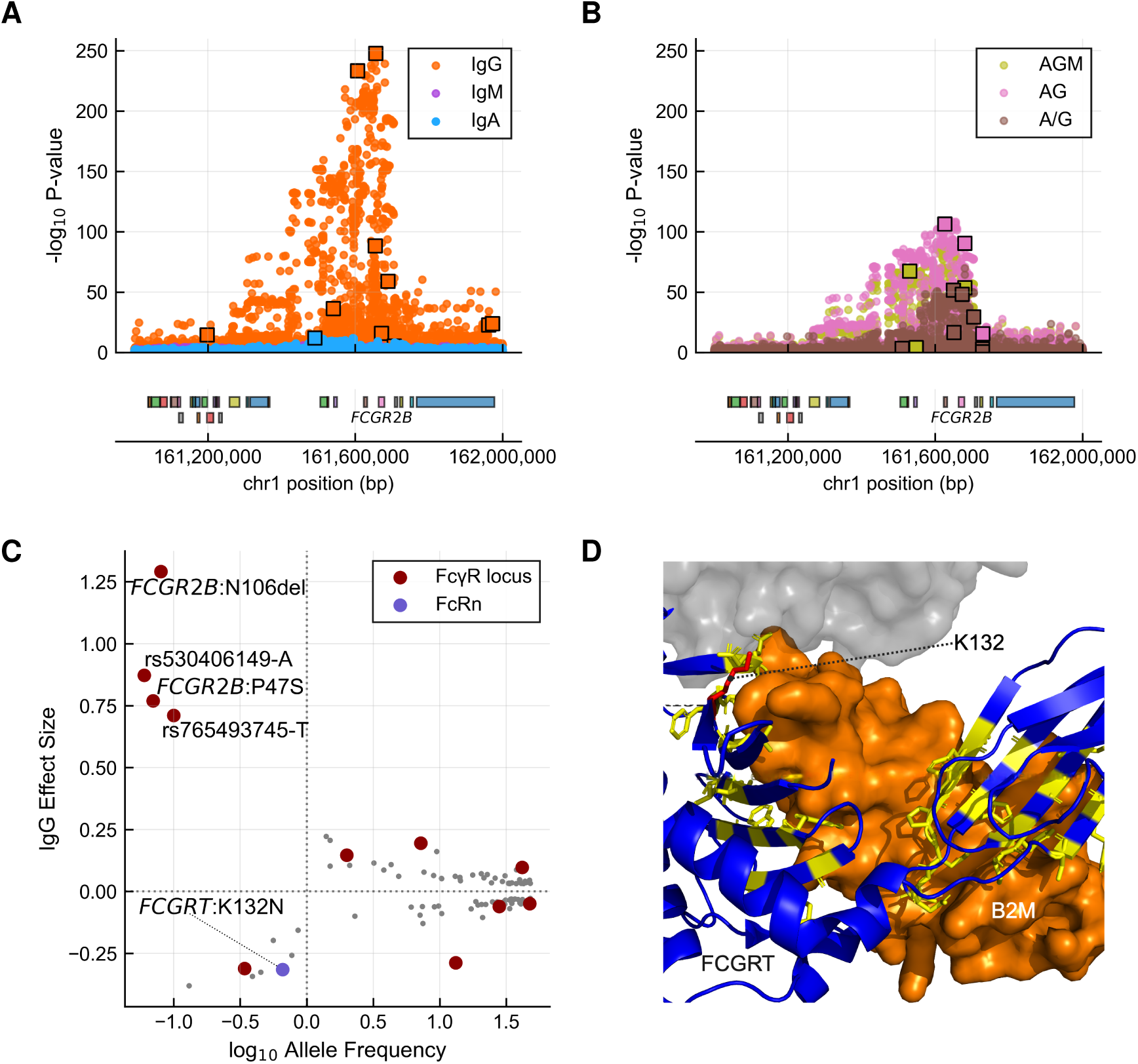
Fcγ receptor variation is a dominant genetic determinant of circulating IgG. **(A)** Regional association plot of the Fcγ receptor (FcγR) locus on chromosome 1q23 for IgG (orange), IgM (magenta), and IgA (cyan). **(B)** Regional association plot of the Fcγ receptor (FcγR) locus on chromosome 1q23 for IgG-derived composite traits: AGM (green), AG (pink), and A/G (grey). **(C)** Effect size *versus* effect allele frequency for IgG-associated variants. Variants mapping to the FcγR locus are highlighted. The three variants with the largest effects on IgG all localize to the FcγR locus, including a novel coding variant in *FCGR2B* (rs149249317-T; p.Pro47Ser). **(D)** Outside the canonical FcγR locus, the neonatal Fc receptor (FCGRT; FcRn) protects IgG from lysosomal degradation and mediates recycling into circulation. A rare coding variant in *FCGRT* (rs150420714-C; p.Lys132Asn; EAF = 0.7%) markedly reduces IgG and is predicted to impair protein function. Shown is the crystal structure of FcRn (blue ribbon) in complex with β2-microglobulin (B2M; orange surface; accession no. PDB 4N0U). FcRn residues within 4.0 Å of B2M are highlighted in yellow. Lys132 (red) lies at the FcRn–B2M interface, suggesting that the p.Lys132Asn substitution may disrupt this interaction required for FcRn-mediated IgG recycling.

First, the well-characterized *FCGR2A* variant rs1801274-A (p.His167Arg) reduces IgG2 binding affinity and lowers the A/G ratio (β_A/G_ = −0.038, P_A/G_ = 4.6×10^-11^; **Table S9**), consistent with impaired clearance of IgG2 immune complexes. Second, rs143596860-A downregulates the inhibitory Fcγ receptor *FCGR2B* (FcγRIIb) in dendritic cells (β = −0.60, *P =* 2.0×10^-44^; **Table S3B**; **Figure S9A**) and increases class-switched Ig (β_AG_ = 0.08, P_AG_ = 9.4×10^-52^; **Table S9**), consistent with reduced inhibitory feedback. Third, rs12139524-C downregulates *FCRLB* in memory B-cell subsets (**Figure S9B**; **Table S3B**) and reduces class-switched Ig (β_AG_ = −0.05, P_AG_ = 7×10^-19^; **Table S9**). *FCRLB* is an intracellular protein of largely unknown function expressed in B cells; the only prior loss-of-function study in mice was confounded by linkage with *Fcgr2b*^89^. These data provide human genetic evidence supporting a role for *FCRLB* in promoting class-switched Ig production. Finally, a rare coding variant in *FCGR2B* (rs149249317-T, p.Pro47Ser; EAF = 0.07%) markedly increases IgG (β_IgG_ = 0.77, P_IgG_ = 1.1×10^-15^; **Figure S9C–D**; **Table S9**). Pro47 lies in the disordered interdomain linker adjacent to the first IgC2-type domain of the FcγRIIb ectodomain (**Figure S9C**). Although predicted benign (AlphaMissense = 0.085; CADD = 0.13), variant classifiers are known to have reduced sensitivity in such regions^90^ and proline is enriched in non-helical interdomain linkers, where it confers structural rigidity^91^, raising the possibility that substitution to flexible serine at this position perturbs FcγRIIb function.

These results show that IgG traits resolve functional variation at the complex FcγR locus and highlight the central role of FcγRIIb autoinhibition in regulating IgG levels.

#### A hypomorphic FCGRT variant modulates IgG half-life

The neonatal Fc receptor (FcRn), encoded by *FCGRT* at chromosome 19q13.33, protects IgG from lysosomal degradation and recycles it into circulation, a process central to IgG’s long half-life (∼3 weeks)^92^. IgG Fc variants that enhance FcRn binding are bioengineered into IgG-based therapeutic antibodies to extend their half-life, whereas pharmacological inhibition of FcRn is under investigation to decrease the half-life of endogenous IgG in the treatment of IgG-mediated autoimmune diseases^93^.

We identify a rare coding variant, rs150420714-C (p.Lys132Asn, EAF = 0.7%), that markedly reduces circulating IgG (β_IgG_ = -0.32, P_IgG_ = 8.7×10^-19^; **Figure 6C**) and is predicted to impair protein function (AlphaMissense = 0.5; CADD = 23.9). A crystal structure of the FcRn–β2 microglobulin complex (Protein Data Bank no. 4N0U)^94^ places Lys132 at the β2 microglobulin-binding interface, an interaction known to be essential for FCGRT function (**Figure 6D**). Carriers of rs150420714-C exhibit reduced IgG levels, supporting the interpretation that this variant reduces IgG half-life through increased lysosomal degradation.

This rare coding variant links human *FCGRT* variation to steady-state IgG levels and may contribute to inter-individual variation in responses to IgG-based therapies.

#### Allelic series at the TACI–APRIL axis quantitatively tune humoral immunity

Quantitative control of humoral immunity is mediated by the TACI–APRIL signaling axis. APRIL (*TNFSF13*), produced primarily by myeloid cells, binds TACI (*TNFRSF13B*) on mature B cells and plasma cells to promote class switching and antibody production. Disruption of this pathway causes IgA deficiency and CVID^48,49^. Across *TNFRSF13B* and *TNFSF13*, we identify allelic series that quantitatively tune TACI–APRIL signaling.

At *TNFRSF13B* (TACI), four signals associate with Ig traits. Two rare extracellular missense variants, Ala181Glu (rs72553883-T; EAF = 0.88%) and Cys104Arg (rs34557412-G; EAF = 0.34%), abrogate TACI signaling^95^ and markedly reduce total Ig (β_AGM_ = -0.57 and -0.25, P_AGM_ = 4.1×10^-28^ and 5.7×10^-10^; **Figure 7A**), while increasing CVID risk^48,49^. Both variants reduce plasma TACI in *cis* (**Figure 7A**; **Table S5A**), expand pre-switched memory B cells in BloodVariome (**Figure S5D,E**), and increase plasma FCRL2 (pre-switched memory B-cell marker; **Table S5A**), while reducing mature B-cell and plasma-cell markers (TNFRSF17/BCMA and IL5RA; **Figure 7A; Table S5A**), collectively supporting a developmental bottleneck at the class-switch transition^96,97^.

**Figure 7.**
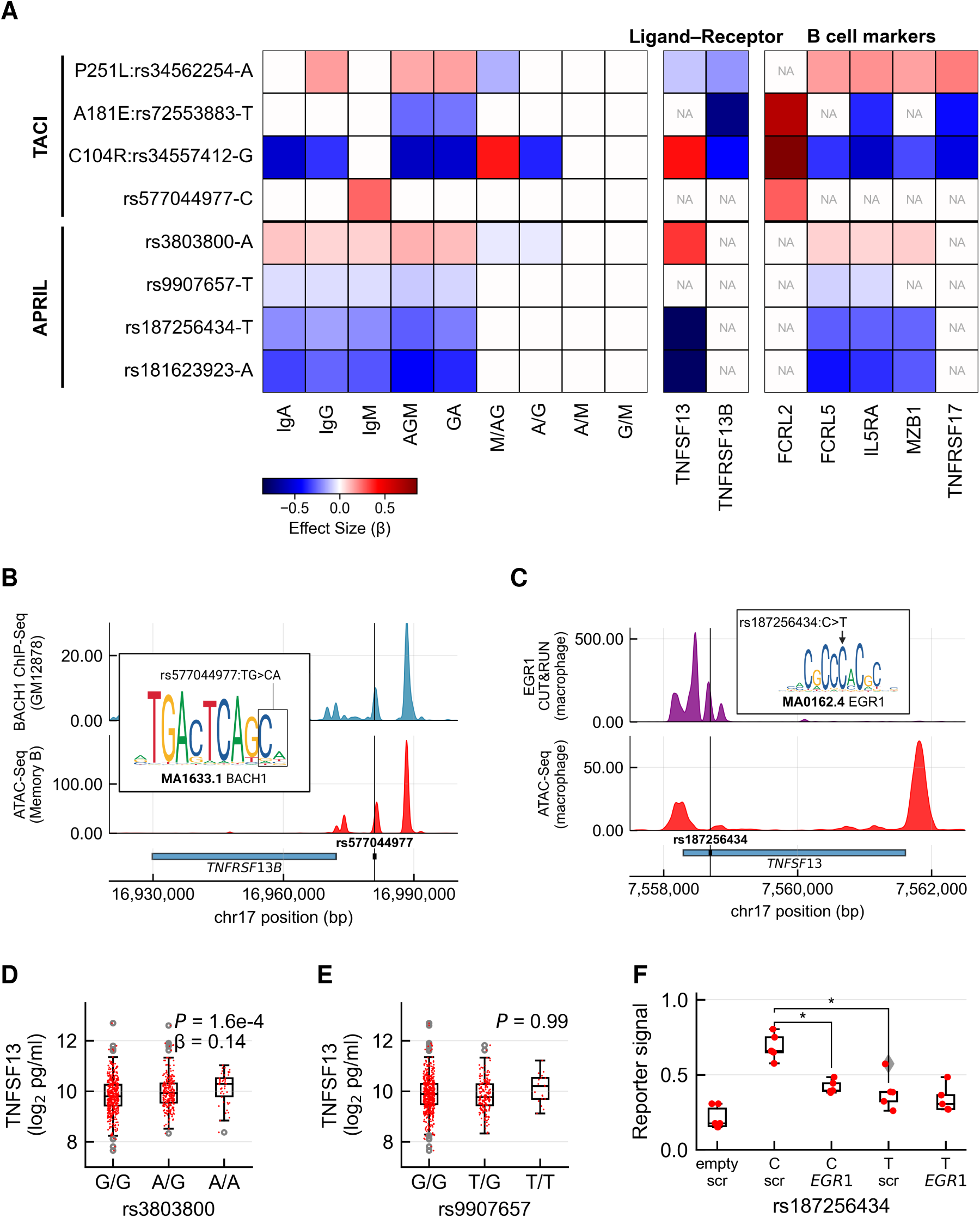
Allelic series at the APRIL–TACI axis quantitatively tune humoral immunity. **(A)** Effect-size heatmaps for allelic series at *TNFRSF13B* (TACI) and *TNFSF13* (APRIL). *Left*: effects across Ig traits. *Right*: plasma pQTLs related to TACI–APRIL signaling and B-cell subpopulation markers. FCRL2 marks unswitched memory B cells, with elevated levels concordant with accumulation of unswitched memory B cells observed by flow cytometry (**Figure S5D,E**). TNFRSF17, IL5RA, MZB1, and FCRL5 reflect mature B-cell and plasma-cell pool size^58^. **(B)** *TNFRSF13B* (TACI) locus showing the location of rs577044977 overlaid with ATAC-sequencing data for memory B cells^69^ and ChIP-seq data for BACH1 in GM12878 cells^107^. rs577044977-C is predicted to increase binding of the BACH1 transcriptional repressor, consistent with reduced *TNFRSF13B* expression (**Figure S10A**). **(C)** *TNFSF13* (APRIL) locus showing the location of rs187256434 overlaid with ATAC-sequencing^108^ and EGR1 CUT&RUN data^109^ for macrophages. The rare variant rs187256434-T overlaps an EGR1 binding site and is predicted to disrupt the EGR1 motif, consistent with reduced TNFSF13 protein levels in plasma pQTL data (**Table S5A**). **(D)** ELISA quantification of plasma APRIL in 793 blood donors stratified by rs3803800 genotype. The A allele is associated with increased APRIL, consistent with *cis*-pQTL effects in proteomics data (**Table S5A**). **(E)** ELISA quantification of plasma APRIL stratified by rs9907657 genotype showing no significant association (regression *P =* 0.99). **(F)** Luciferase reporter assays for rs187256434 alleles in the monocytic MOLM-13 cell line. The T allele shows reduced activity relative to the C allele. siRNA-mediated knockdown of EGR1 abolishes the allelic difference by selectively reducing C allele activity, supporting disruption of an EGR1 binding site by rs187256434-T.

In contrast, the intracellular variant Pro251Leu (rs34562254-A; EAF = 9.5%) shows reciprocal effects – elevating IgA and IgG, reducing FCRL2, and increasing BCMA and IL5RA – consistent with enhanced TACI signaling and increased multiple myeloma risk (**Figure 7A; Table S5A**)^58,98^. Finally, a rare upstream regulatory variant, rs577044977-C (EAF = 0.53%), downregulates *TNFRSF13B* mRNA in blood and elevates IgM and plasma FCRL2, consistent with selective expansion of IgM-producing memory B cells (**Figure 7A, S10A**). Collectively, these four variants delineate a graded continuum of TACI perturbations affecting class switching, plasma-cell output, and disease risk.

At *TNFSF13* (APRIL), four variants converge on coordinated regulation of all three Ig isotypes via distinct cellular mechanisms (**Figure 7A**). Two common variants act through separate myeloid compartments: rs3803800-A increases Ig by upregulating *TNFSF13* in monocytes, whereas rs9907657-T lowers Ig by downregulating *TNFSF13* in neutrophils (**Figure S10B-C**). Notably, only rs3803800-A elevates plasma APRIL, confirmed by both proteomics and ELISA in 793 blood donors (**Figure 7D-E**; **Tables S5A** and **S10B-C**). These data genetically implicate monocytes rather than neutrophils as a primary source of circulating plasma APRIL in humans^99^. In addition, two rare *TNFSF13* variants, rs181623923-A (EAF = 0.77%) and rs187256434-T (EAF = 0.86%), markedly reduce total Ig (**Table S2B**) and plasma APRIL (**Table S5A**). rs181623923-A downregulates *TNFSF13* mRNA in blood, whereas rs187256434-T disrupts an *EGR1* binding site in macrophages (**Figure 7C**) and reduces transcriptional activity, as confirmed by luciferase assays and EGR1 siRNA knockdown in monocytic MOLM-13 cells (**Figure 7F, S11, Table S11B**). Given that rare *TNFSF13* loss-of-function mutations have been reported in CVID^100^, these regulatory variants may represent hypomorphic alleles influencing immunodeficiency risk.

Together, the *TNFRSF13B* and *TNFSF13* allelic series demonstrate how graded perturbations of a central ligand–receptor axis quantitatively tune circulating Ig levels, B-cell state composition, and risk of immunodeficiency and plasma cell malignancy.

## DISCUSSION

Population-scale genetics of immunoglobulin traits provides a window into the human B-cell system. Across 114,697 individuals we identify 504 variants influencing IgA, IgG, IgM, and composite Ig traits. Integration of these signals with regulatory genomics, plasma proteomics, and immune cell phenotyping shows that variation in circulating immunoglobulins provides genetically interpretable readouts of B-cell system state.

Circulating Ig levels have traditionally been viewed as relatively coarse clinical measures influenced by demographic and environmental factors. Our results show that, when analyzed at population scale, these traits encode detailed information about the B-cell system. Genetic perturbations affecting antibody levels recover many established regulators of B-cell biology while also implicating genes not previously linked to humoral immunity. In this way, naturally occurring genetic variation functions as a series of *in vivo* perturbations that expose the regulatory organization of the human humoral immune response.

Several loci identified here highlight key control points in antibody production. At the TACI–APRIL signaling axis, multiple independent variants influence immunoglobulin levels, plasma protein signatures, and B-cell subset composition. Together these variants form allelic series producing graded perturbations of antibody output, illustrating how germline variation can quantitatively tune humoral immune set points. Similar patterns are observed at Fcγ receptor loci and within the immunoglobulin heavy-chain region, where genetic variants influence class-switch dynamics and subclass distribution. These observations indicate that naturally occurring genetic variation modulates humoral immunity across multiple regulatory layers spanning B-cell activation, differentiation, and antibody secretion.

The genetic architecture of Ig traits intersects extensively with immune-mediated disease. Many loci overlap risk variants for autoimmunity, immunodeficiency, and B-cell malignancy, placing quantitative variation in antibody production at the interface between physiological immune regulation and pathology. Several regulatory nodes highlighted by this study – including Fcγ receptors, the TACI–APRIL signaling pathway, and the neonatal Fc receptor – are established or emerging therapeutic targets.

Several limitations should be noted. Most mechanistic inferences derive from genetic and molecular association analyses and will require experimental validation. The cohorts analyzed are predominantly of Northern European ancestry, and extending studies of immunoglobulin genetics to more diverse populations will be important for refining causal inference and understanding the global architecture of humoral immune regulation. Finally, circulating immunoglobulin levels represent the aggregate output of the B-cell system and do not capture antigen specificity or context-dependent immune responses.

Together these findings show that population-scale genetics of immunoglobulin traits can resolve key features of the regulatory architecture of the human B-cell system. More broadly, they illustrate how large-scale human genetic variation can be leveraged to dissect complex immune systems directly in their physiological context.

## DATA AVAILABILITY

Summary statistics for the association studies of the nine Ig traits have been deposited in the GWAS Catalog database (https://www.ebi.ac.uk/gwas/; accession number pending). The BloodVariome data are available at (https://bloodvariome.medfak.lu.se/). Gene expression datasets are available from the NCBI Gene Expression Omnibus repository (https://www.ncbi.nlm.nih.gov/geo/; accession numbers GSE107011, GSE139369)^101,102^ and the ProteinAtlas (https://www.proteinatlas.org/). Disease annotations were retrieved from the GWAS Catalog (https://www.ebi.ac.uk/gwas/), ClinVar (https://www.ncbi.nlm.nih.gov/clinvar/), OMIM (https://www.omim.org/), IntOGen (https://www.intogen.org/), and the Mitelman Database (https://mitelmandatabase.isb-cgc.org/). The pooled CRISPR-Cas9 screening data for cancer cell lines is available through the DepMap portal (https://depmap.org).

## ACKNOWLEDGEMENTS

This work was supported by grants from the Swedish Research Council (2017-02023 and 2018-00424), the Swedish Cancer Society (200696), European Research Council (consolidator grant no. 770992 BloodVariome and EU-MSCA-COFUND 754299 CanFaster), and Region Skåne. We thank the personnel at Clinical Chemistry and Clinical Immunology and Transfusion Medicine for their assistance with sample collection. We are indebted to the blood donors and patients who participated in the study.

## AUTHOR CONTRIBUTIONS

Z.A., A.L.d.L.P., B.N., I.J., G.T., T.O., K.S. designed the study. C.C., B.R.L, T.L.J., S.Y.K., S.J.H., K.F. collected samples and data, Z.A., G.T., L.E., B.N., A.L.d.L.P., G.H.H., P.M., L.J., A.J., A.S., A.L., M.O. analyzed data. Z.A., A.L.d.L.P., G.T., I.J., T.O., B.N. drafted the manuscript. All authors contributed to the final manuscript.

## Abbreviations

ATAC-seq: assay for transposase-accessible chromatin sequencing
BAFF: B-cell activating factor
BCR: B-cell receptor
ChIP-seq: chromatin immunoprecipitation sequencing
CVID: common variable immunodeficiency
DZ: dark zone
eQTL: expression quantitative trait locus
FcγR: Fcγ receptor
GC: germinal center
GWAS: genome-wide association study
IGH: immunoglobulin heavy chain
IVW: inverse-variance weighted
LZ: light zone
pQTL: protein quantitative trait locus
scATAC-seq: single-cell ATAC-seq
scRNA-seq: single-cell RNA sequencing
TACI: transmembrane activator and CAML interactor
3′RR1: 3′ regulatory region 1.

## MATERIALS AND METHODS

### Study design and population

#### Iceland

Preexisting IgA, IgG and IgM measurements were obtained from Landspitali, the National University Hospital of Iceland (LSH), Akureyri Hospital (FSA) and the Icelandic Medical Center (Laeknasetrid) Laboratory in Mjodd (RAM), Reykjavik, Iceland and The Iceland Screens, Treats, or Prevents Multiple Myeloma (iStopMM) study. All measurements made in LSH, FSA and RAM between 1991 and 2020 were included in this study. iStopMM is a nationwide, population-based screening study and randomized controlled trial conducted among Icelandic residents born in 1975 or earlier. All eligible residents (n = 148,711) were invited to participate between September 2016 and February 2018. A total of 80,759 individuals (54.3%) provided informed consent. The study protocol, information materials, biobank, and questionnaires were approved by the Icelandic National Bioethics Committee (No. 16-022, 2016-04-26) and the Icelandic Data Protection Authority, with registration at ClinicalTrials.gov (NCT03327597)^110^.

Following enrollment of iStopMM, serum samples for screening were collected in conjunction with routine clinical blood draws within the universal Icelandic healthcare system. The electronic sampling infrastructure links major hospital and primary care laboratories, covering ∼92% of the Icelandic population. In total, 75,422 serum samples were retrieved for analysis. Samples were transported to the clinical laboratory at Landspítali – The National University Hospital of Iceland (Reykjavík), where serum was aliquoted using automated Thermo Scientific® TC automation and aliquoter instruments. Each sample was assigned a pseudonymous study identification code. All serum samples were stored and managed according to standardized laboratory procedures before shipment for analysis.

Quantification of IgG, IgA, and IgM, in the iStopMM study was performed at The Binding Site laboratory (Birmingham, UK) using Hevylite® immunoassays on the Optilite® turbidimeter (The Binding Site Group Ltd., Birmingham, UK) in accordance with the manufacturer’s instructions.

#### Sweden

IgA, IgG, and IgM levels were measured in randomly ascertained blood donors and primary care patients from Skåne country in southern Sweden. EDTA tubes were collected subject to ethical approval (Lund University Ethical Review Board, dnr 2022-01414-02). IgA, IgG, and IgM were measured using clinically accredited methods at the Clinical Chemistry unit at Skåne University Hospital. The measured values were log-transformed and standardized by subtracting the mean from each value and dividing by the standard deviation. After deduplication and geographic filtering, 7,841 individuals with IgA values, 7,842 with IgG, 7,841 with IgM were available for association analysis.

#### United Kingdom

The UK Biobank cohort includes information on phenotype and genotype data from approximately half a million citizens from across the UK who have all provided informed consent for their data to be used for scientific research^111^. The Immunoglobulin measurements were obtained from the Primary Care data using the following Read codes: IgA(43r1.,XaJoT,43J5.,XE25B); IgG (Xa7kR,43J3.,XE259); and IgM (43me.,XaJpg,43J4.,XE25A). This yielded 52,869 Ig measurements for 15,282 individuals. This research was conducted under the UKB application number 56270.

### Genotyping and genetic ancestry analysis

#### Iceland

Genotypes for 173,025 Icelanders that have been genotyped using Illumina SNP chips were long-range phased^112^ and variants identified in the whole-genome sequences of 63,460 Icelanders were imputation into chip-typed individuals and their close non-chip-typed family members^113^. The sequencing was done using Illumina standard TruSeq methodology. Only samples with a genome-wide average coverage of 20X were considered. Autosomal SNPs and INDEL’s were called using Graphtyper version 1.4.5^114^. Variants that did not pass quality control were excluded from the analysis. Information about haplotype sharing was used to improve variant genotyping, taking advantage of the fact that all sequenced individuals had also been chip-typed and long-range phased.

#### UK

Sample for the participants from UK Biobank were genotype using Affymetrix UK BiLEVE Axiom or UK Biobank Axiom chips, long range phased and imputed based in a reference set of whole-genome sequencing data for 150,119 UK Biobank samples^115^. A subset of 431,079 individuals of British–Irish ancestry was defined^115^ and for that subset 20 genetic principal components were generated and used to adjust the trait values prior to association analysis. This subset included 13,612 individuals with Ig measurements.

#### Sweden

The Swedish samples were genotyped Illumina OmniExpress-24 and Global Screening single-nucleotide polymorphism microarrays and phased together with 570,100 samples of North-Western Europe origin using Eagle^116^. Samples and variants with < 98% yield were excluded. For imputation, we created a haplotype reference panel by phasing the whole-genome sequencing genotypes for 17,408 individuals of European origin, including 3,704 Swedish samples, sequenced using HiSeqX and NovaSeq PCR-free Illumina technology to a mean depth of at least 30X^113^. Single-nucleotide polymorphisms and indels were called using joint calling with Graphtyper^114^. The 17,408 individuals were also genotyped using the Illumina OmniExpress and Global Screening microarrays and those genotypes were long-range phased using Eagle. The sequence genotypes were phased with chip genotypes and 170 million variants identified in the whole-genome sequencing data were imputed into the phased chip data. The population structure was analysed using ADMIXTURE v1.23^117^ in supervised mode with 1,000 Genomes populations CEU, CHB, and YRI as training samples^118^ and Swedish individuals as test samples. Samples with < 0.9 CEU ancestry were excluded. To identify and exclude individuals of Finnish/Saami origin, we included individuals with less than 0.3 CHB and less than 0.05 YRI ancestry. The remaining samples were projected onto 20 principal components calculated with a European reference panel. UMAP^119^ was used to reduce test sample coordinates to two dimensions. Additional European samples not in the original reference set were also projected onto the principal components and UMAP components to identify ancestries, and samples with Swedish ancestry were identified. The inclusion of Finnish/Saami individuals allowed us to confirm that we could identify a distinct UMAP cluster of individuals with the following properties: **(a)** elevated CHB ancestry according to ADMIXTURE; **(b)** enriched for individuals who we knew to have been born in Finland; **(c)** on principal component 1 and 2, were positioned in the region occupied by Finnish individuals. Those individuals were excluded and the association analysis included 7,847 individuals with Ig measurements.

### Association testing

Prior to association testing, the log transformed trait values were adjusted for covariates. For Iceland the measured values were adjusted for sex, age and laboratory, for UK the adjustment was done for sex, year of birth, age, age squared and 20 PC’s, and for Sweden for sex and 20 PC’s. After adjustment an average was calculated for individuals with multiple measurements and the adjusted trait values were standardized using an inverse normal transform. The trait values were tested for association with 62 million genetic variants using a linear mixed model implemented in BOLT-LMM^116^, using the expected allele count from the imputation as the explanatory variable and normalized trait values as response. Linkage disequilibrium score regression was used to account for distributional inflation in the test statistics due to cryptic relatedness and population stratification^13^. To account for multiple testing, we applied a class-based Bonferroni correction, grouping variants by genomic annotation and adjusting the significance threshold for each category based on the number of variants^10^.

Stepwise conditional analysis was used to identify independent signals in each associated region for each of the nine Ig traits. The genome was divided into regions of genome-wide association signals, separated by at least 1 Mb, then adjusted for the strongest association signal in the region. If, after conditioning on the lead signal, if there was still a genome-wide significant association in the region, the process was repeated, adjusting both signals. This was repeated until there was no significant association in the region. Finally, all identified signals were adjusted for all the other signals. To define the 99% credible set of plausible causal variants for each association signal, we applied a Bayesian refinement approach Maller, 2012 #1356}. For regions with multiple signals this was done separately for each signal using conditionally adjusted P values for all variants.

### Inter-trait colocalization

To identify shared association signals between multiple traits, we performed inter-trait colocalization using the coloc package^12,120^. We identified all 95% credible sets that overlap with at least one variant as candidates for colocalization and calculated colocalization probabilities for such overlapping pairs. Pairs that show a posterior probability of colocalization (PP.H4) greater than 0.9 were considered as representing shared signals. Groups of colocalizing association signals were then combined into shared association signals. This can create situations where a single association signal is represented by multiple, distinct lead variants corresponding to distinct credible sets. In these cases, we take the lead variant corresponding to the most significant trait.

### Identification of candidate genes

To identify candidate genes, we searched for coding variants and colocalizing eQTLs within the 95% credible sets. We prioritized a gene as a candidate gene if: (i) the credible set contains a non-synonymous coding variant (missense, frameshift, stop-gain, splice donor/acceptor) with posterior inclusion probability> 0.1; or (ii) the association colocalizes with a *cis*-eQTL in eQTL data for sorted immune cells (posterior probability PP.H4 > 0.9). For eQTL analysis, we used the following datasets: macrophages (n= 84)^121^; monocytes, neutrophils, CD4^+^ T cells (n = 197)^122^; regulatory T cells (n = 119)^123^; CD4^+^ and CD8^+^ T cells, monocytes, neutrophils, platelets, B cells, ileum, rectum, transverse colon (n = 322)^124^; B cells (n = 282)^125^; monocytes (n = 424)^126^; fibroblasts, T cells, lymphoblastoid cell lines (n = 195)^127^; Natural killer cells (n = 247)^128^; CD4^+^ and CD8^+^ T cells (n = 297)^129^; whole blood (n = 491)^130^; neutrophils (n = 93)^131^; macrophages (n =1 68)^132^; monocytes (n = 200)^133^; fifteen immune cell types (n = 91)^134^; and microglia (n = 104)^135^. If neither criterion was met, we prioritized the protein-coding gene closest to the lead variant.

### ATAC-sequencing enrichment analysis

To assess the enrichment of Ig-associated variants in accessible chromatin of different immune cell types, we used ATAC-sequencing data for 25 bulk-sorted immune cell types (NCBI GEO accession no. GSE118165) To analyze the data for bulk-sorted cells, we used gchromVAR (https://github.com/caleblareau/gchromVAR/). B cell enrichment was assessed by comparing the mean gchromVAR z-score difference between B cell (n = 4) and non-B cell (n = 21) types, averaged across all nine traits, against the exact permutation distribution obtained by reassigning the B cell label to all possible groupings of four cell types.

### Analysis of candidate gene expression across tissues

To test for enrichment of candidate genes in tissues, we used the ProteinAtlas single-cell type dataset^103^, which encompasses mRNA sequencing data for 81 cell types across a broad range of human tissues, including the immune cell lineages analyzed in our study. Using a one-sided Student’s t-test, we compared normalized expression values for candidate vs other genes in the genome. To annotate candidate genes with expression across blood and immune cells at high cell type resolution, we used publicly available RNA-sequencing data from Monaco *et al.* and Granja *et al.* (Gene Expression Omnibus accession no. GSE107011 and GSE139369)^101,102^.

### Gene set enrichment analysis

We performed gene set enrichment analysis only using high-confidence candidate genes supported by molecular evidence. We used the EnrichR^18^ tool to determine enrichment of high-confidence candidate genes in the Reactome pathway database^17^ for biological processes, the MGI database of mutant mice phenotypes, the OMIM database^41^ for disease-associated genes and the IntOGen^42^ and Mitelman^43^ databases for somatic driver genes in cancers.

### Transcription factor binding and motif prediction

To identify transcription factors that overlap credible set variants, we used the ReMap2022^136^ database and the FABIAN-variant tool^137^ to predict binding motifs affected by those variants. These analyses were done in a focused way for specific association signals, using prior knowledge to prioritize ChIP-sequencing datasets from primary immune cells and transcription factors with well-characterized role in B cell physiology.

### IgG subclass outlier analysis

In our prior GWAS on IgG1-4 subclass levels, we observed reciprocal effects on IgG1 and IgG3 of the *IGHG1* deletion-tagging variant rs587597004-T^87^. To search for buffering effects across IgG-associated variants, we therefore applied an outlier detection approach based on the expected proportional relationship between subclass and total IgG levels. We reasoned that variants influencing total IgG should produce proportional changes across all subclasses, whereas variants with compensatory effects on IgG subclass levels should produce reciprocal effects that deviate from this pattern.

To identify genetic variants with disproportionate effects on individual IgG subclasses relative to total IgG, we performed outlier detection using inverse variance-weighted (IVW) regression. For each subclass, we computed residuals from the IVW fit as the difference between the observed subclass effect size and the effect predicted by the IVW slope (β_IVW_ × β_IgG_). Residuals were standardized by dividing by the standard error of the subclass effect estimate to obtain unitless deviation scores comparable across variants with differing measurement precision. Variants were classified as outliers if the absolute value of their standardized residual exceeded two standard deviations of the standardized residual distribution across all IgG-associated variants. We defined buffered outliers as those variants that exhibited reciprocal effects (*i.e.*, a positive effect on at least one subclass, and negative on at least one other subclass). We performed this analysis in an unbiased manner for 89 IgG-associated variants that could be successfully mapped to our IgG1-4 subclass dataset. We detected ten outliers, including five with reciprocal effects consistent with compensatory buffering, all which mapped to the *IGH* locus.

### Three-dimensional protein modeling

The three-dimensional model of the FCGRT-B2M complex (Protein Data Bank identifier 4NOU)^94^ was created using PyMOL (https://pymol.org/). Additionally, we used PyMOL to label all amino acid residues in FCGRT within 4 Å of B2M.

### Plasma ELISA

We used a commercial ELISA kit (R&D Systems, catalog no. DY884B) to detect TNFSF13. ELISA was performed on blood samples for the BloodVariome Compendium taken from Lund University Hospital Clinical Chemistry, which were left overnight at room temperature to allow plasma separation. To ensure that this did not result in TNFSF13 protein loss due to degradation, we obtained fresh blood samples and found that overnight storage at room temperature did not significantly reduce ELISA readings. In total, we analyzed 793 samples, which were also genotyped as part of the BloodVariome compendium.

### Luciferase assay and siRNA knockdown

We synthesized 70-bp sequences centered on rs187256434 (Integrated DNA Technologies) and cloned them into the pGL3-Basic reporter vector using KpnI and BglII restriction sites. Each sequence was centered on the variant, and the two constructs differed only for rs187256434. 240 ng of Renilla luciferase construct were co-transfected with 10 μg of firefly construct in 5 million MOLM-13 cells, using the Neon electroporation system (Thermo Scientific). At 24 h post-electroporation, luciferase and Renilla activity were measured using DualGlo Luciferase (Promega; #E1960) on a GLOMAX 20/20 Luminometer. For co-electroporation, luciferase plasmids were co-transfected with 50 nM *EGR1* siRNA (Qiagen; #1027416) or 50 nM control siRNA (Qiagen; #1022076). Optimal siRNA concentration was determined by serial dilution, and knockdown efficiency was evaluated using qPCR quantification of *EGR1* mRNA (Thermo Scientific #4331182, assay id: Hs00152928_m1).

### Disease relevance

To identify disease risk variants overlapping with Ig-associated variants, we searched the GWAS Catalog (https://www.ebi.ac.uk/gwas/; *r*^2^ > 0.8 between Ig and catalog lead variants) and ClinVar (https://; non-benign coding variants overlapping our credible sets). To find links to Mendelian disorders characterized by abnormal immune cell function, we used OMIM (https://www.omim.org/)^41^. To investigate the overlap with genes that are somatically mutated in different tumor types, we considered genes listed as cancer driver genes in IntOGen (https://www.intogen.org/)^42^ and fusion gene partners that were recurrently reported (five or more times) in the Mitelman database^43^ (https://mitelmandatabase.isb-cgc.org). To identify effects of gene knockdown on the growth of cell lines derived from immune cell malignancies, we used the Dependency Map version 24Q4 (https://depmap.org)^44^.

## REFERENCES

1 Abbas, A. K., Lichtman, A. H. & Pillai, S. Cellular and molecular immunology. Eighth edition. edn, (Elsevier Saunders, 2015).

2 Jonsson, S. et al. Identification of sequence variants influencing immunoglobulin levels. Nat Genet 49, 1182–1191 (2017). 10.1038/ng.3897

3 Viktorin, A. et al. IgA measurements in over 12 000 Swedish twins reveal sex differential heritability and regulatory locus near CD30L. Hum Mol Genet 23, 4177–4184 (2014). 10.1093/hmg/ddu135

4 Weidinger, S. et al. Genome-wide scan on total serum IgE levels identifies FCER1A as novel susceptibility locus. PLoS Genet 4, e1000166 (2008). 10.1371/journal.pgen.1000166

5 Granada, M. et al. A genome-wide association study of plasma total IgE concentrations in the Framingham Heart Study. J Allergy Clin Immunol 129, 840–845 e821 (2012). 10.1016/j.jaci.2011.09.029

6 Liao, M. et al. Genome-wide association study identifies common variants at TNFRSF13B associated with IgG level in a healthy Chinese male population. Genes Immun 13, 509–513 (2012). 10.1038/gene.2012.26

7 Yang, C. et al. Genome-wide association study identifies TNFSF13 as a susceptibility gene for IgA in a South Chinese population in smokers. Immunogenetics 64, 747–753 (2012). 10.1007/S00251-012-0636-Y/METRICS

8 Swaminathan, B. et al. Variants in ELL2 influencing immunoglobulin levels associate with multiple myeloma. Nat Commun 6, 7213 (2015). 10.1038/ncomms8213

9 Liu, L. et al. Genetic regulation of serum IgA levels and susceptibility to common immune, infectious, kidney, and cardio-metabolic traits. Nature Communications 2022 13:1 13, 1-17 (2022). 10.1038/s41467-022-34456-6

10 Sveinbjornsson, G. et al. Weighting sequence variants based on their annotation increases power of whole-genome association studies. Nature Genetics 48, 314–317 (2016). 10.1038/ng.3507

11 Maller, J. B. et al. Bayesian refinement of association signals for 14 loci in 3 common diseases. Nature Genetics 44, 1294–1301 (2012). 10.1038/ng.2435

12 Wallace, C. A more accurate method for colocalisation analysis allowing for multiple causal variants. PLOS Genetics 17, e1009440–e1009440 (2021). 10.1371/JOURNAL.PGEN.1009440

13 Bulik-Sullivan, B. et al. LD score regression distinguishes confounding from polygenicity in genome-wide association studies. Nature Genetics 47, 291–295 (2015). 10.1038/NG.3211;TECHMETA=43,45;SUBJMETA=205,208,2138,457,631;KWRD=GENOME-WIDE+ASSOCIATION+STUDIES,POPULATION+GENETICS

14 Ota, M. et al. Dynamic landscape of immune cell-specific gene regulation in immune-mediated diseases. Cell 184, 3006–3021.e3017 (2021). 10.1016/j.cell.2021.03.056

15 Kerimov, N. et al. A compendium of uniformly processed human gene expression and splicing quantitative trait loci. Nature Genetics 2021 53:9 53, 1290-1299 (2021). 10.1038/s41588-021-00924-w

16 Bogue, M. A. et al. Mouse Phenome Database: A data repository and analysis suite for curated primary mouse phenotype data. Nucleic Acids Research 48, D716–D723 (2020). 10.1093/nar/gkz1032

17 Gillespie, M. et al. The reactome pathway knowledgebase 2022. Nucleic acids research 50, D687–D692 (2022). 10.1093/NAR/GKAB1028

18 Chen, E. Y. et al. Enrichr: Interactive and collaborative HTML5 gene list enrichment analysis tool. BMC Bioinformatics 14, 1–14 (2013). 10.1186/1471-2105-14-128/FIGURES/3

19 Karlsson, M. et al. A single-cell type transcriptomics map of human tissues. Sci Adv 7 (2021). 10.1126/sciadv.abh2169

20 Xu, H. et al. Regulation of bifurcating B cell trajectories by mutual antagonism between transcription factors IRF4 and IRF8. Nature Immunology 2015 16:12 16, 1274-1281 (2015). 10.1038/ni.3287

21 Sasaki, Y. & Iwai, K. Roles of the NF-κB pathway in B-lymphocyte biology. Current Topics in Microbiology and Immunology 393, 177–209 (2015). 10.1007/82_2015_479

22 Fedl, A. S. et al. Transcriptional function of E2A, Ebf1, Pax5, Ikaros and Aiolos analyzed by in vivo acute protein degradation in early B cell development. Nature Immunology 25, 1663-1677 (2024). 10.1038/S41590-024-01933-7;SUBJMETA=100,102,250,2502,337,631;KWRD=CHROMATIN+REMODELLING,GENE+REGULATION+IN+IMMUNE+CELLS

23 West, M. J. & Farrell, P. J. Roles of RUNX in B cell immortalisation. Advances in Experimental Medicine and Biology 962, 283–298 (2017). 10.1007/978-981-10-3233-2_18

24 Xie, P., Kraus, Z. J., Stunz, L. L. & Bishop, G. A. Roles of TRAF molecules in B lymphocyte Function. Cytokine & growth factor reviews 19, 199–199 (2008). 10.1016/J.CYTOGFR.2008.04.002

25 Mackay, F. & Schneider, P. TACI, an enigmatic BAFF/APRIL receptor, with new unappreciated biochemical and biological properties. Cytokine & growth factor reviews 19, 263–276 (2008). 10.1016/j.cytogfr.2008.04.006

26 Yu, G. et al. APRIL and TALL-1 and receptors BCMA and TACI: system for regulating humoral immunity. Nature Immunology 2000 1:3 1, 252-256 (2000). 10.1038/79802

27 Conway, K. L. et al. ATG5 regulates plasma cell differentiation. Autophagy 9, 528–537 (2013). 10.4161/auto.23484

28 Martincic, K., Alkan, S. A., Cheatle, A., Borghesi, L. & Milcarek, C. Transcription elongation factor ELL2 directs immunoglobulin secretion in plasma cells by stimulating altered RNA processing. Nat Immunol 10, 1102–1109 (2009). 10.1038/ni.1786

29 O’Connell, P., Blake, M. K., Godbehere, S., Amalfitano, A. & Aldhamen, Y. A. SLAMF7 modulates B cells and adaptive immunity to regulate susceptibility to CNS autoimmunity. Journal of Neuroinflammation 19, 1–18 (2022). 10.1186/S12974-022-02594-9/FIGURES/7

30 Takai, T. Roles of Fc receptors in autoimmunity. Nature Reviews Immunology 2002 2:8 2, 580-592 (2002). 10.1038/nri856

31 Baldwin, W. M., Valujskikh, A. & Fairchild, R. L. The neonatal Fc receptor: Key to homeostasic control of IgG and IgG-related biopharmaceuticals. American Journal of Transplantation 19, 1881–1887 (2019). 10.1111/AJT.15366

32 Qian, S. et al. The role of BCL-2 family proteins in regulating apoptosis and cancer therapy. Front Oncol 12, 985363 (2022). 10.3389/fonc.2022.985363

33 Ware, C. F. Network communications: lymphotoxins, LIGHT, and TNF. Annu Rev Immunol 23, 787–819 (2005). 10.1146/annurev.immunol.23.021704.115719

34 Wang, Y. et al. Lymphotoxin Beta Receptor Signaling in Intestinal Epithelial Cells Orchestrates Innate Immune Responses against Mucosal Bacterial Infection. Immunity 32, 403–413 (2010). 10.1016/J.IMMUNI.2010.02.011

35 Luong, P. et al. INAVA-ARNO complexes bridge mucosal barrier function with inflammatory signaling. Elife 7, e38539–e38539 (2018). 10.7554/ELIFE.38539

36 Yan, J., Hedl, M. & Abraham, C. An inflammatory bowel disease-risk variant in INAVA decreases pattern recognition receptor-induced outcomes. J Clin Invest 127, 2192–2205 (2017). 10.1172/JCI86282

37 Lauc, G. et al. Loci associated with N-glycosylation of human immunoglobulin G show pleiotropy with autoimmune diseases and haematological cancers. PLoS Genet 9, e1003225 (2013). 10.1371/journal.pgen.1003225

38 Medvedovic, J., Ebert, A., Tagoh, H. & Busslinger, M. Pax5: a master regulator of B cell development and leukemogenesis. Advances in immunology 111, 179–206 (2011). 10.1016/B978-0-12-385991-4.00005-2

39 Welter, D. et al. The NHGRI GWAS Catalog, a curated resource of SNP-trait associations. Nucleic Acids Res 42, D1001–1006 (2014). 10.1093/nar/gkt1229

40 Landrum, M. J. et al. ClinVar: Improving access to variant interpretations and supporting evidence. Nucleic Acids Research 46, D1062–D1067 (2018). 10.1093/nar/gkx1153

41 McKusick, V. A. Mendelian Inheritance in Man and its online version, OMIM. Am J Hum Genet 80, 588–604 (2007). 10.1086/514346

42 Martinez-Jimenez, F. et al. A compendium of mutational cancer driver genes. Nat Rev Cancer 20, 555–572 (2020). 10.1038/s41568-020-0290-x

43 Mitelman, F., Johansson, B. & Mertens, F.

44 Arafeh, R., Shibue, T., Dempster, J. M., Hahn, W. C. & Vazquez, F. The present and future of the Cancer Dependency Map. Nat Rev Cancer 25, 59–73 (2025). 10.1038/s41568-024-00763-x

45 McMaster, M. L. et al. Two high-risk susceptibility loci at 6p25.3 and 14q32.13 for Waldenström macroglobulinemia. Nature Communications 9, 4182–4182 (2018). 10.1038/S41467-018-06541-2

46 Sakaue, S. et al. A cross-population atlas of genetic associations for 220 human phenotypes. Nat Genet 53, 1415–1424 (2021). 10.1038/s41588-021-00931-x

47 Guler, M. & Canzian, F. Clustering of lymphoid neoplasms by cell of origin, somatic mutation and drug usage profiles: a multi-trait genome-wide association study. Blood Cancer J 15, 147 (2025). 10.1038/s41408-025-01351-4

48 Salzer, U. et al. Mutations in TNFRSF13B encoding TACI are associated with common variable immunodeficiency in humans. Nature genetics 37, 820–828 (2005). 10.1038/NG1600

49 Salzer, U. & Grimbacher, B. TACI deficiency — a complex system out of balance. Current Opinion in Immunology 71, 81–88 (2021). 10.1016/J.COI.2021.06.004

50 Dobbs, K. et al. Inherited DOCK2 Deficiency in Patients with Early-Onset Invasive Infections. New England Journal of Medicine 372, 2409–2422 (2015). 10.1056/NEJMOA1413462/SUPPL_FILE/NEJMOA1413462_DISCLOSURES.PDF

51 Moens, L. et al. Human DOCK2 Deficiency: Report of a Novel Mutation and Evidence for Neutrophil Dysfunction. Journal of Clinical Immunology 39, 298–298 (2019). 10.1007/S10875-019-00603-W

52 Eldjarn, G. H. et al. Large-scale plasma proteomics comparisons through genetics and disease associations. Nature 2023 622:7982 622, 348-358 (2023). 10.1038/s41586-023-06563-x

53 Sun, B. B. et al. Plasma proteomic associations with genetics and health in the UK Biobank. Nature 2023 622:7982 622, 329-338 (2023). 10.1038/s41586-023-06592-6

54 Ferkingstad, E. et al. Large-scale integration of the plasma proteome with genetics and disease. Nature Genetics 53, 1712–1721 (2021). 10.1038/s41588-021-00978-w

55 Oskam, N., et al. CD5L is a canonical component of circulatory IgM. Proceedings of the National Academy of Sciences of the United States of America 120, e2311265120-e2311265120 (2023). 10.1073/PNAS.2311265120;PAGE:STRING:ARTICLE/CHAPTER

56 Wang, Y., Su, C., Ji, C. & Xiao, J. CD5L associates with IgM via the J chain. Nat Commun 15, 8397 (2024). 10.1038/s41467-024-52175-y

57 Jerala, R. Structural biology of the LPS recognition. Int J Med Microbiol 297, 353–363 (2007). 10.1016/j.ijmm.2007.04.001

58 Went, M. et al. Deciphering the genetics and mechanisms of predisposition to multiple myeloma. Nat Commun 15, 6644 (2024). 10.1038/s41467-024-50932-7

59 Lied, G. A. & Berstad, A. Functional and clinical aspects of the B-cell-activating factor (BAFF): a narrative review. Scand J Immunol 73, 1–7 (2011). 10.1111/j.1365-3083.2010.02470.x

60 Boyd, R. S. et al. Proteomic analysis of the cell-surface membrane in chronic lymphocytic leukemia: Identification of two novel proteins, BCNP1 and MIG2B. Leukemia 17, 1605-1612 (2003). 10.1038/SJ.LEU.2402993;KWRD=MEDICINE

61 Hong, R. et al. The B cell novel protein 1 (BCNP1) regulates BCR signaling and B cell apoptosis. European Journal of Immunology 49, 911–917 (2019). 10.1002/EJI.201847985;WGROUP:STRING:PUBLICATION

62 Hong, R. et al. Distinct roles of BCNP1 in B-cell development and activation. International Immunology 32, 17–26 (2020). 10.1093/INTIMM/DXZ055

63 Pervushin, N. V., Nilov, D. K., Zhivotovsky, B. & Kopeina, G. S. Bcl-2 modifying factor (Bmf): “a mysterious stranger” in the Bcl-2 family proteins. Cell Death & Differentiation 2025, 1–12 (2025). 10.1038/s41418-025-01562-z

64 Labi, V. et al. Loss of the BH3-only protein Bmf impairs B cell homeostasis and accelerates γ irradiation–induced thymic lymphoma development. Journal of Experimental Medicine 205, 641–655 (2008). 10.1084/JEM.20071658

65 Lin, Y. L., Ip, P. P. & Liao, F. CCR6 Deficiency Impairs IgA Production and Dysregulates Antimicrobial Peptide Production, Altering the Intestinal Flora. Front Immunol 8, 805 (2017). 10.3389/fimmu.2017.00805

66 Saevarsdottir, S. et al. FLT3 stop mutation increases FLT3 ligand level and risk of autoimmune thyroid disease. Nature 2020 584:7822 584, 619-623 (2020). 10.1038/s41586-020-2436-0

67 Baglaenko, Y. et al. Precisely defining disease variant effects in CRISPR-edited single cells. Nature 646, 117–125 (2025). 10.1038/s41586-025-09313-3

68 Lim, J. X. et al. PD-1 receptor deficiency enhances CD30(+) T(reg) cell function in melanoma. Nat Immunol 26, 1074–1086 (2025). 10.1038/s41590-025-02172-0

69 Calderon, D. et al. Landscape of stimulation-responsive chromatin across diverse human immune cells. Nature Genetics 51, 1494–1505 (2019). 10.1038/s41588-019-0505-9

70 Krysiak, K. et al. Recurrent somatic mutations affecting B-cell receptor signaling pathway genes in follicular lymphoma. Blood 129, 473–473 (2017). 10.1182/BLOOD-2016-07-729954

71 Reddy, A. et al. Genetic and Functional Drivers of Diffuse Large B Cell Lymphoma. Cell 171, 481–489.e415 (2017). 10.1016/j.cell.2017.09.027

72 Basso, K. & Dalla-Favera, R. BCL6. master regulator of the germinal center reaction and key oncogene in B Cell lymphomagenesis. Advances in Immunology 105, 193–210 (2010). 10.1016/S0065-2776(10)05007-8

73 Lalani, S. R. et al. MCTP2 is a dosage-sensitive gene required for cardiac outflow tract development. Human Molecular Genetics 22, 4339–4348 (2013). 10.1093/HMG/DDT283

74 Shin, O. H., Han, W., Wang, Y. & Sudhof, T. C. Evolutionarily conserved multiple C2 domain proteins with two transmembrane regions (MCTPs) and unusual Ca2+ binding properties. J Biol Chem 280, 1641–1651 (2005). 10.1074/jbc.M407305200

75 Dominguez-Sola, D. et al. The FOXO1 Transcription Factor Instructs the Germinal Center Dark Zone Program. Immunity 43, 1064–1074 (2015). 10.1016/j.immuni.2015.10.015

76 Wang, H. et al. Transcription factors IRF8 and PU.1 are required for follicular B cell development and BCL6-driven germinal center responses. Proceedings of the National Academy of Sciences of the United States of America 116, 9511-9520 (2019). 10.1073/PNAS.1901258116/SUPPL_FILE/PNAS.1901258116.SD02.XLS

77 Cocco, M. et al. A dichotomy of gene regulatory associations during the activated B-cell to plasmablast transition. Life Sci Alliance 3 (2020). 10.26508/lsa.202000654

78 Nutt, S. L., Hodgkin, P. D., Tarlinton, D. M. & Corcoran, L. M. The generation of antibody-secreting plasma cells. Nature reviews. Immunology 15, 160–171 (2015). 10.1038/nri3795

79 Iho, S., Takahashi, T., Kura, F., Sugiyama, H. & Hoshino, T. The effect of 1,25-dihydroxyvitamin D3 on in vitro immunoglobulin production in human B cells. J Immunol 136, 4427–4431 (1986).

80 Vincenzi, F., Smirne, C., Tonello, S. & Sainaghi, P. P. The Role of Vitamin D in Autoimmune Diseases. Int J Mol Sci 27 (2026). 10.3390/ijms27010555

81 Yang, C. Y., Leung, P. S., Adamopoulos, I. E. & Gershwin, M. E. The implication of vitamin D and autoimmunity: a comprehensive review. Clin Rev Allergy Immunol 45, 217–226 (2013). 10.1007/s12016-013-8361-3

82 Adorini, L. & Penna, G. Control of autoimmune diseases by the vitamin D endocrine system. Nat Clin Pract Rheumatol 4, 404–412 (2008). 10.1038/ncprheum0855

83 Cantorna, M. T. Why do T cells express the vitamin D receptor? Ann N Y Acad Sci 1217, 77–82 (2011). 10.1111/j.1749-6632.2010.05823.x

84 Knippenberg, S. et al. Effect of vitamin D(3) supplementation on peripheral B cell differentiation and isotype switching in patients with multiple sclerosis. Mult Scler 17, 1418–1423 (2011). 10.1177/1352458511412655

85 Plum, L. A. et al. Antibody production in mice requires neither vitamin D, nor the vitamin D receptor. Front Immunol 13, 960405 (2022). 10.3389/fimmu.2022.960405

86 Honjo, T. IMMUNOGLOBULIN GENES. Ann. Rev. ImmunoL 1, 499–528 (1983).

87 Olafsdottir, T. A. et al. Sequence variants influencing the regulation of serum IgG subclass levels. Nature Communications 2024 15:1 15, 1-13 (2024). 10.1038/s41467-024-52470-8

88 Jodice, C. et al. Variation of the 3’RR1 HS1.2 Enhancer and Its Genomic Context. Genes (Basel) 15 (2024). 10.3390/genes15070856

89 Masuda, K. et al. Defining the immunological phenotype of Fc receptor-like B (FCRLB) deficient mice: Confounding role of the inhibitory FcgammaRIIb. Cell Immunol 266, 24–31 (2010). 10.1016/j.cellimm.2010.08.007

90 Luppino, F., Lenz, S., Chow, C. F. W. & Toth-Petroczy, A. Deep learning tools predict variants in disordered regions with lower sensitivity. BMC Genomics 26, 367 (2025). 10.1186/s12864-025-11534-9

91 George, R. A. & Heringa, J. An analysis of protein domain linkers: their classification and role in protein folding. Protein Eng 15, 871–879 (2002). 10.1093/protein/15.11.871

92 Foss, S. et al. Human IgG Fc-engineering for enhanced plasma half-life, mucosal distribution and killing of cancer cells and bacteria. Nat Commun 15, 2007 (2024). 10.1038/s41467-024-46321-9

93 Sesarman, A., Vidarsson, G. & Sitaru, C. The neonatal Fc receptor as therapeutic target in IgG-mediated autoimmune diseases. Cell Mol Life Sci 67, 2533–2550 (2010). 10.1007/s00018-010-0318-6

94 Oganesyan, V. et al. Structural insights into neonatal Fc receptor-based recycling mechanisms. J Biol Chem 289, 7812–7824 (2014). 10.1074/jbc.M113.537563

95 Fried, A. J., Rauter, I., Dillon, S. R., Jabara, H. H. & Geha, R. S. Functional analysis of transmembrane activator and calcium-modulating cyclophilin ligand interactor (TACI) mutations associated with common variable immunodeficiency. Journal of Allergy and Clinical Immunology 128, 226–228.e221 (2011). 10.1016/J.JACI.2011.01.048

96 Martinez-Gallo, M. et al. TACI mutations and impaired B-cell function in subjects with CVID and healthy heterozygotes. Journal of Allergy and Clinical Immunology 131, 468–476 (2013). 10.1016/j.jaci.2012.10.029

97 Pulvirenti, F. et al. Clinical Associations of Biallelic and Monoallelic *TNFRSF13B* Variants in Italian Primary Antibody Deficiency Syndromes. Journal of Immunology Research 2016, 1-14 (2016). 10.1155/2016/8390356

98 Ajore, R. et al. Functional dissection of inherited non-coding variation influencing multiple myeloma risk. Nat Commun 13, 151 (2022). 10.1038/s41467-021-27666-x

99 Weldon, A. J. et al. Surface APRIL Is Elevated on Myeloid Cells and Is Associated with Disease Activity in Patients with Rheumatoid Arthritis. J Rheumatol 42, 749–759 (2015). 10.3899/jrheum.140630

100 Yeh, T. W. et al. APRIL-dependent lifelong plasmacyte maintenance and immunoglobulin production in humans. J Allergy Clin Immunol 146, 1109–1120 e1104 (2020). 10.1016/j.jaci.2020.03.025

101 Granja, J. M. et al. Single-cell multiomic analysis identifies regulatory programs in mixed-phenotype acute leukemia. Nat Biotechnol 37, 1458–1465 (2019). 10.1038/s41587-019-0332-7

102 Monaco, G. et al. RNA-Seq Signatures Normalized by mRNA Abundance Allow Absolute Deconvolution of Human Immune Cell Types. Cell reports 26, 1627–1640.e1627 (2019). 10.1016/j.celrep.2019.01.041

103 Uhlen, M. et al. Proteomics. Tissue-based map of the human proteome. Science 347, 1260419 (2015). 10.1126/science.1260419

104 Minderjahn, J. et al. Mechanisms governing the pioneering and redistribution capabilities of the non-classical pioneer PU.1. Nature Communications 2020 11:1 11, 1-16 (2020). 10.1038/s41467-019-13960-2

105 King, H. W. et al. Integrated single-cell transcriptomics and epigenomics reveals strong germinal center-associated etiology of autoimmune risk loci. Science immunology 6 (2021). 10.1126/SCIIMMUNOL.ABH3768

106 Lu, X. et al. MTA2/NuRD Regulates B Cell Development and Cooperates with OCA-B in Controlling the Pre-B to Immature B Cell Transition. Cell reports 28, 472–485 e475 (2019). 10.1016/j.celrep.2019.06.029

107 Consortium, E. P. An integrated encyclopedia of DNA elements in the human genome. Nature 489, 57–74 (2012). 10.1038/nature11247

108 Novakovic, B. et al. beta-Glucan Reverses the Epigenetic State of LPS-Induced Immunological Tolerance. Cell 167, 1354–1368 e1314 (2016). 10.1016/j.cell.2016.09.034

109 Trizzino, M. et al. EGR1 is a gatekeeper of inflammatory enhancers in human macrophages. Sci Adv 7 (2021). 10.1126/sciadv.aaz8836

110 Rognvaldsson, S. et al. Monoclonal gammopathy of undetermined significance and COVID-19: a population-based cohort study. Blood Cancer J 11, 191 (2021). 10.1038/s41408-021-00580-7

111 Bycroft, C. et al. The UK Biobank resource with deep phenotyping and genomic data. Nature 562, 203–209 (2018). 10.1038/s41586-018-0579-z

112 Kong, A. et al. Detection of sharing by descent, long-range phasing and haplotype imputation. Nat Genet 40, 1068–1075 (2008). 10.1038/ng.216

113 Gudbjartsson, D. F. et al. Large-scale whole-genome sequencing of the Icelandic population. Nature Genetics 47, 435–444 (2015). 10.1038/ng.3247

114 Eggertsson, H. P. et al. Graphtyper enables population-scale genotyping using pangenome graphs. Nature Genetics 49, 1654–1660 (2017). 10.1038/ng.3964

115 Halldorsson, B. V. et al. The sequences of 150,119 genomes in the UK Biobank. Nature 607, 732–740 (2022). 10.1038/s41586-022-04965-x

116 Loh, P. R. et al. Efficient Bayesian mixed-model analysis increases association power in large cohorts. Nat Genet 47, 284–290 (2015). 10.1038/ng.3190

117 Alexander, D. H., Novembre, J. & Lange, K. Fast model-based estimation of ancestry in unrelated individuals. Genome Research 19, 1655–1664 (2009). 10.1101/gr.094052.109

118 Auton, A. et al. in Nature Vol. 526 68-74 (Nature Publishing Group, 2015).

119 Diaz-Papkovich, A., Anderson-Trocmé, L. & Gravel, S. in Journal of Human Genetics Vol. 66 85–91 (Springer Nature, 2021).

120 Giambartolomei, C. et al. Bayesian Test for Colocalisation between Pairs of Genetic Association Studies Using Summary Statistics. PLoS Genetics 10 (2014). 10.1371/journal.pgen.1004383

121 Alasoo, K. et al. Shared genetic effects on chromatin and gene expression indicate a role for enhancer priming in immune response. Nat Genet 50, 424–431 (2018). 10.1038/s41588-018-0046-7

122 Chen, L. et al. Genetic Drivers of Epigenetic and Transcriptional Variation in Human Immune Cells. Cell 167, 1398–1414 e1324 (2016). 10.1016/j.cell.2016.10.026

123 Bossini-Castillo, L. et al. Immune disease variants modulate gene expression in regulatory CD4(+) T cells. Cell Genom 2, None (2022). 10.1016/j.xgen.2022.100117

124 Momozawa, Y. et al. IBD risk loci are enriched in multigenic regulatory modules encompassing putative causative genes. Nat Commun 9, 2427 (2018). 10.1038/s41467-018-04365-8

125 Fairfax, B. P. et al. Genetics of gene expression in primary immune cells identifies cell type-specific master regulators and roles of HLA alleles. Nat Genet 44, 502–510 (2012). 10.1038/ng.2205

126 Fairfax, B. P. et al. Innate immune activity conditions the effect of regulatory variants upon monocyte gene expression. Science 343, 1246949 (2014). 10.1126/science.1246949

127 Gutierrez-Arcelus, M. et al. Passive and active DNA methylation and the interplay with genetic variation in gene regulation. Elife 2, e00523 (2013). 10.7554/eLife.00523

128 Gilchrist, J. J. et al. Natural Killer cells demonstrate distinct eQTL and transcriptome-wide disease associations, highlighting their role in autoimmunity. Nat Commun 13, 4073 (2022). 10.1038/s41467-022-31626-4

129 Kasela, S. et al. Pathogenic implications for autoimmune mechanisms derived by comparative eQTL analysis of CD4+ versus CD8+ T cells. PLoS Genet 13, e1006643 (2017). 10.1371/journal.pgen.1006643

130 Lepik, K. et al. C-reactive protein upregulates the whole blood expression of CD59 - an integrative analysis. PLoS Comput Biol 13, e1005766 (2017). 10.1371/journal.pcbi.1005766

131 Naranbhai, V. et al. Genomic modulators of gene expression in human neutrophils. Nat Commun 6, 7545 (2015). 10.1038/ncomms8545

132 Nedelec, Y. et al. Genetic Ancestry and Natural Selection Drive Population Differences in Immune Responses to Pathogens. Cell 167, 657–669 e621 (2016). 10.1016/j.cell.2016.09.025

133 Quach, H. et al. Genetic Adaptation and Neandertal Admixture Shaped the Immune System of Human Populations. Cell 167, 643–656 e617 (2016). 10.1016/j.cell.2016.09.024

134 Schmiedel, B. J. et al. Impact of Genetic Polymorphisms on Human Immune Cell Gene Expression. Cell 175, 1701–1715 e1716 (2018). 10.1016/j.cell.2018.10.022

135 Young, A. M. H. et al. A map of transcriptional heterogeneity and regulatory variation in human microglia. Nat Genet 53, 861–868 (2021). 10.1038/s41588-021-00875-2

136 Hammal, F., de Langen, P., Bergon, A., Lopez, F. & Ballester, B. ReMap 2022: a database of Human, Mouse, Drosophila and Arabidopsis regulatory regions from an integrative analysis of DNA-binding sequencing experiments. Nucleic Acids Res 50, D316–D325 (2022). 10.1093/nar/gkab996

137 Steinhaus, R., Robinson, P. N. & Seelow, D. FABIAN-variant: predicting the effects of DNA variants on transcription factor binding. Nucleic Acids Research 50, W322–W329 (2022). 10.1093/NAR/GKAC393

