## Supplementary Figures for "Population-scale immunoglobulin genetics resolves the human B-cell system"

### Figure S1

**(A)** Distribution of 95% credible set sizes across fine-mapped signals. 175 signals were resolved to fewer than 5 credible set variants, including 70 signals resolved to a single putative causal variant. Two signals at the *MAPT* locus have exceptionally large credible sets ($>$2,600 variants) and extend beyond the $x$-axis cutoff of 200. **(B)** Number of genome-wide significant associations per immunoglobulin trait. Red bars indicate associations exclusive to the indicated trait; white bars indicate associations shared with at least one other trait. **(C)** Effect allele frequency versus effect size for IgM-associated variants. Magenta points indicate variants associated with B-cell cancer risk. **(D)** Effect allele frequency versus effect size for AGM-associated variants. Magenta points indicate variants associated with immunodeficiency or chronic infection.

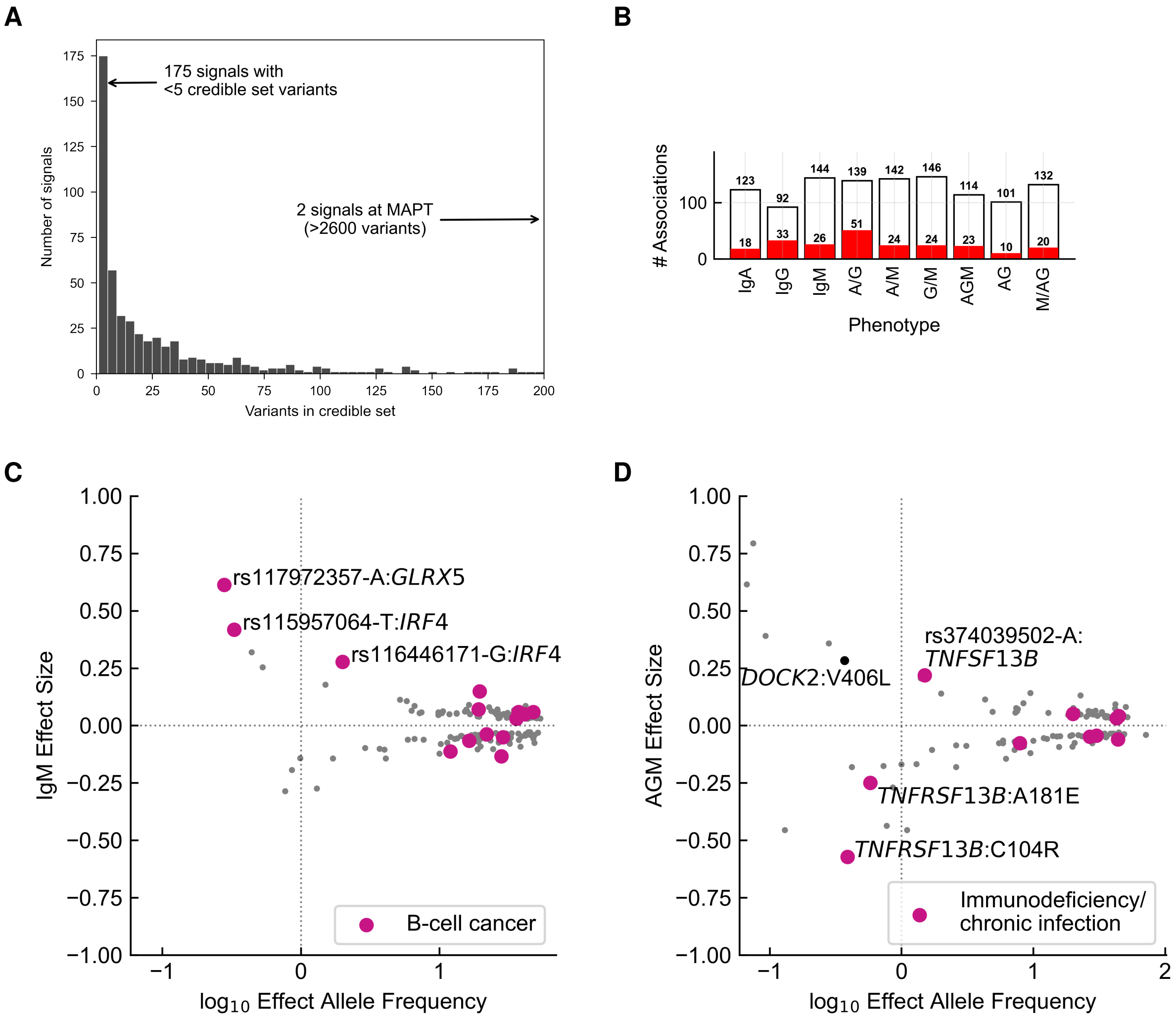

#### Figure S2

Overlap of candidate genes across the three primary immunoglobulin isotypes. 23 loci were shared across all three isotypes, including core B cell transcription factors (*IKZF3*, *IRF4*, *BACH2*), signaling regulators (*TNFRSF13B*, *TNFSF13*), and survival genes (*BCL2*). Isotype-specific loci reflect specialized biology: IgA-exclusive loci support mucosal immunity (e.g., *LTBR*, *INAVA*); IgG-exclusive loci relate to effector function (e.g., Fc$\gamma$ receptors, *ST6GAL1*); IgM-exclusive loci regulate early B cell development (e.g., *PAX5*). Representative genes for each category are annotated.

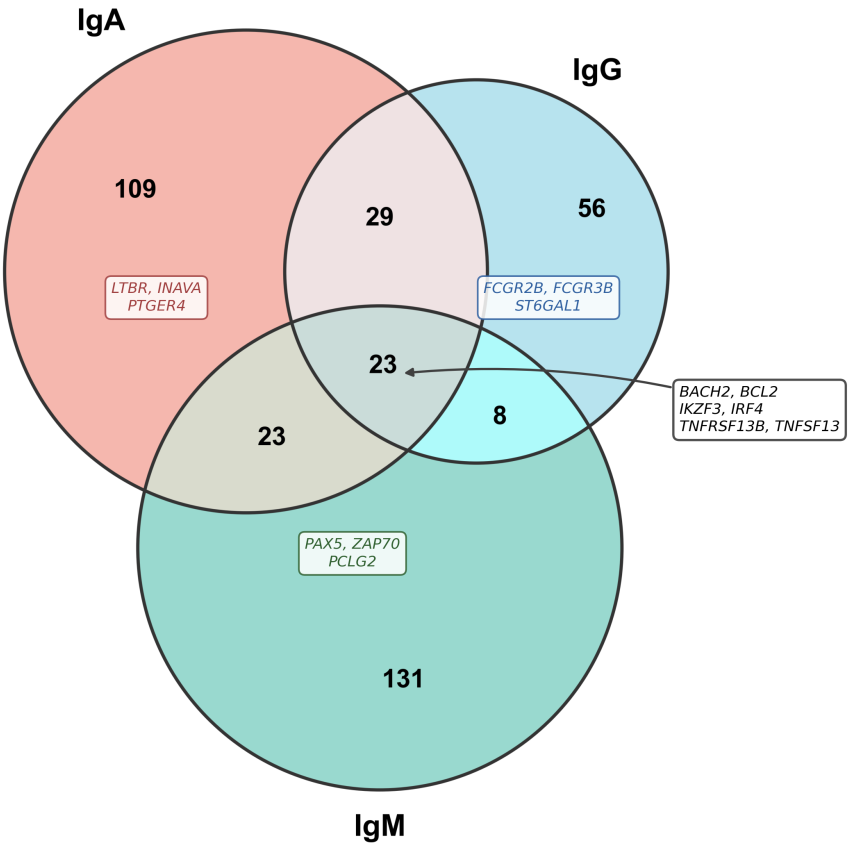

### Figure S3

Overlap between Ig-associated variants and disease risk variants in the GWAS Catalog and ClinVar. Colors indicate lead trait: IgM (deep blue); IgA (teal); IgG (green); AGM (orange); AG (red); A/G (pink); A/M (purple); M/AG (brown); G/M (grey) (**Table S2B**).

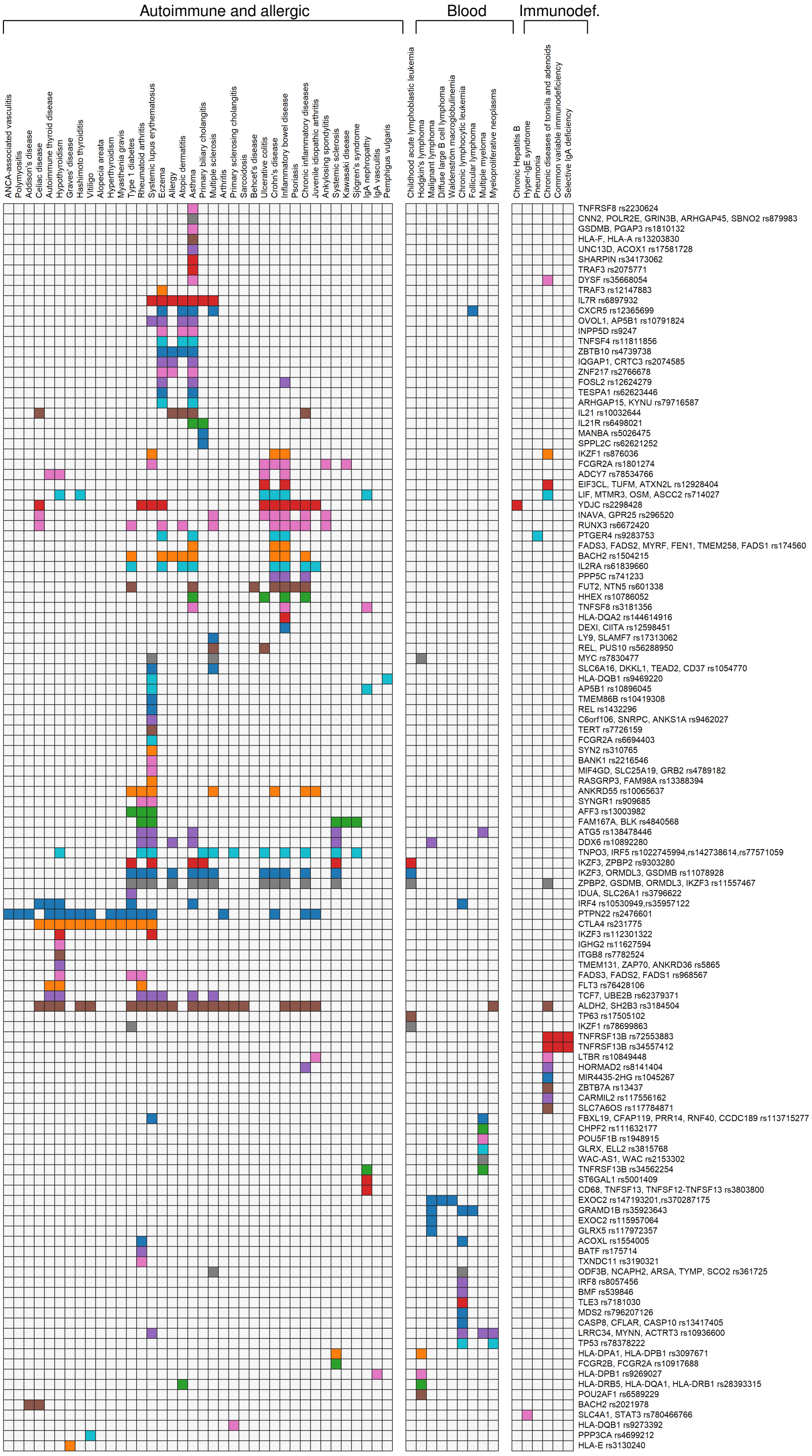

### Figure S4

IVW regression of selected pQTL and Ig effect sizes, with MR-Egger regression lines shown for comparison (dashed). **(A)** JCHAIN and CD5L *vs.* IgM; **(B)** LY96 *vs.* IgA; **(C)** FCRL5, TNFRSF17 (BCMA), MZB1, and IL5RA *vs.* AG. **(D)** Expression of *FCRL5*, *TNFRSF17*, *MZB1*, *IL5RA*, and *LY96* across hematopoiesis (data from PMID 30858613). Dot size reflects absolute expression, color scale relative expression. Cell types: HSC, hematopoietic stem cell; MPP, multipotent progenitor; LMPP, lymphoid-primed multipotent progenitor; CLP, common lymphoid progenitor; CMP, common myeloid progenitor; GMP, granulocyte-monocyte progenitor; MEP, megakaryocyte-erythrocyte progenitor; Ery, erythroid precursor; Mono, monocyte; DC, dendritic cell; NK, natural killer cell; B, B cell; PC, plasma cell. **(E)** Expression of the same genes in unstimulated and stimulated (anti-human IgG/IgM + IL-4) naïve B, memory B, and bulk B cells (data from PMID 31570894). Of the five genes, only *LY96* is upregulated upon B cell stimulation

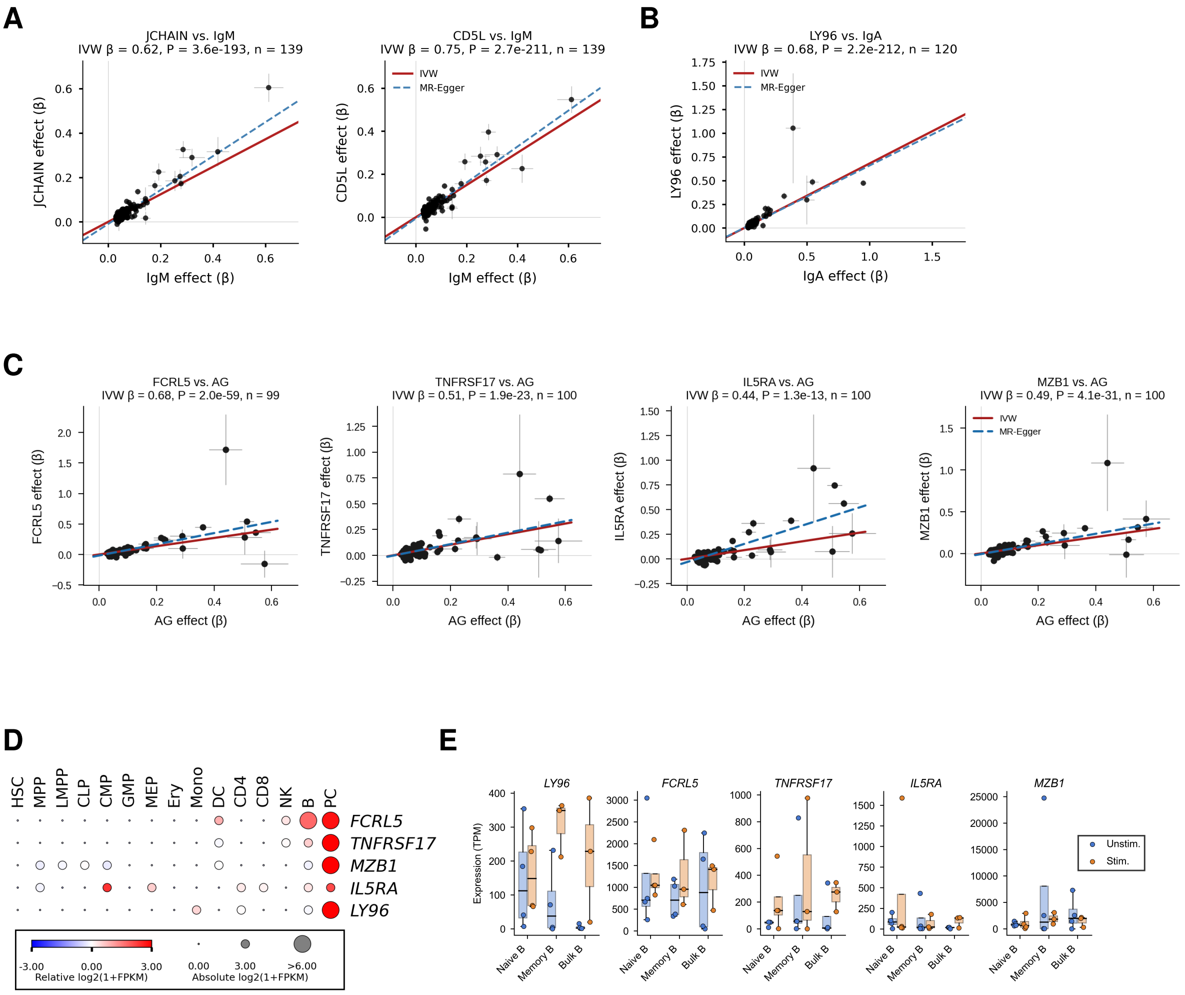

### Figure S5

Effects of Ig-associated variants in the BloodVariome dataset, comprising GWAS data for 1,005 immune cell traits in 11,983 individuals (Lopez de Lapuente Portilla *et al.*, companion manuscript). In short, we used high-resolution flow cytometry and pattern recognition algorithms to profile 92 T cell, 16 B cell, 7 NK cell, 6 monocyte, and 4 dendritic cell (DC) populations. For each population, we quantified and GWAS:ed three kinds of traits: (1) cell frequencies (absolute counts and relative frequencies within parent and grandparent populations; n = 286 traits); (2) surface protein expression (median fluorescence intensity for antibody targets; n = 557); and (3) cell morphology (forward scatter, FSC, as a proxy for size; side scatter, SSC, for internal structural complexity; n = 163).

**(A)**-**(L)** Colocalizing variants with significant effects on B-cell traits, consistent with B-cell-intrinsic cellular effects; **(M)**-**(O)** Variants primarily affecting T helper cells; **(P)**-**(Q)** Variants primarily affecting regulatory T cellds; **(R)**-**(V)** Variants primarily affecting dendritic cells. The title of each subsection describes the BloodVariome variant.

Circular icons represent cell populations. The labels underneath denote associated traits. The length of the bars indicate -$\log_{10} P$-values. Color indicates effect size ($\beta$). A single star indicates genome-wide significance; two stars represent study-wide significance (*i.e.*, further Bonferroni correction for the effective number of independent traits in BloodVariome, estimated at 146 based on phenotype correlations; see Lopez de Lapuente Portilla *et al.*, companion manuscript). Suggestive associations (*i.e.*, those with P-value within two logarithms of genome-wide significance) are also indicated without stars. The color of the cell icon indicates the effect size of the most significant trait associated with that cell type.

Also shown are lead variants and associated traits in the Ig GWAS.

The plots below show candidate gene expression in for hematopoietic cell types (mRNA-sequencing data from PMID 30726743 and 31792411). Color indicates log2-transformed, median-centered relative expression; dot size absolute expression. No heatmap is shown for variants with unclear candidate gene assignments.

Abbreviations: central memory (CM), effector memory (EM), natural killer cell (NK), conventional dendritic cell (cDC), plasmacytoid dendritic cell (pDC), peripheral blood monocytic cells (PBMC), effect allele frequency (EAF), association number this study (loc# XX), association number in the BloodVariome study (cs_XX).

#### A. chr13:108308032[TGCTG→T; EAF 2.099%] (rs200748895, cs_53)

Ig lead variant: rs374039502-A (loc# 362).
Ig associations: IgG↑, IgM↑, IgA*IgG*IgM↑, IgA*IgG↑
Candidate gene: *TNFSF13B* (nearest gene, cis-pQTL)
Candidate mechanism: Low-frequency variant in the 3'-UTR of *TNFSF13B*, which encodes B-cell activating factor (BAFF). This cytokine stimulates B-cell development and is produced by various immune and non-immune cells, mainly myeloid cells. The rs374039502-A variant upregulates total Ig levels. Consistent with this, BloodVariome revealed an association with increased total B cell frequency. Plasma proteomics data from the UK Biobank showed increased plasma levels of TNFSF13B ($\beta$ = 0.79, P = 3.1e-246) and several B-cell and plasma cell markers, including TNFRSF17/BCMA and FCRL5 (**Table S29**).

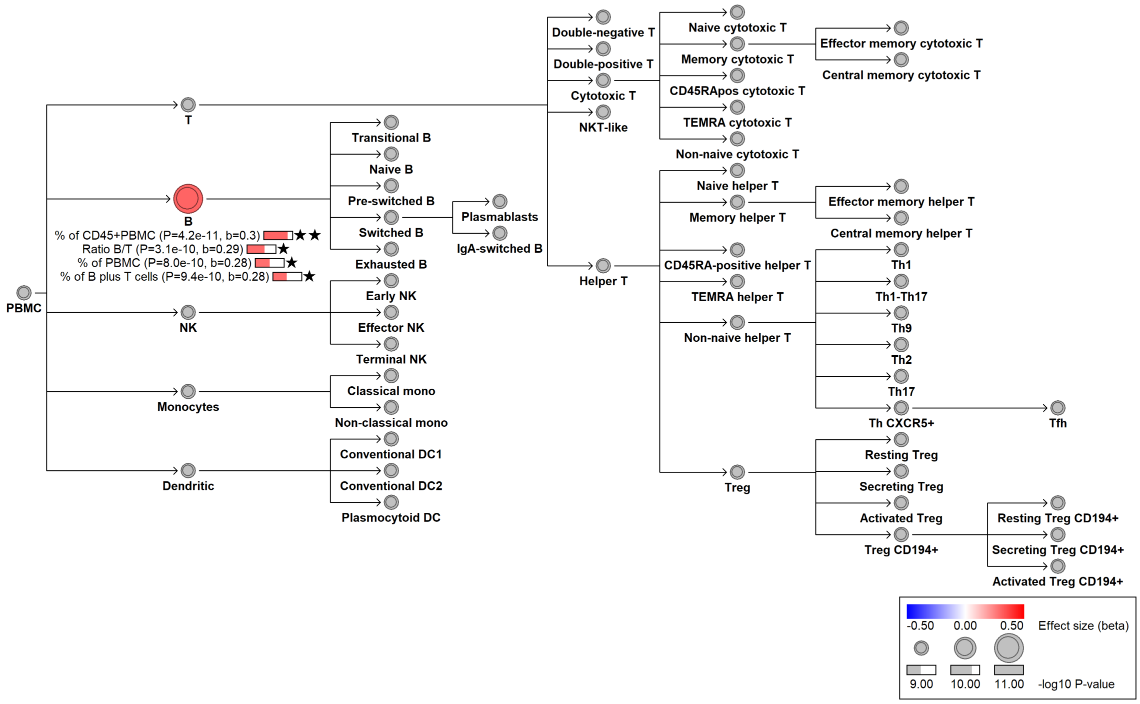


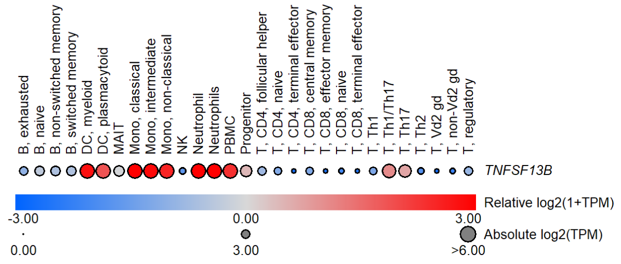


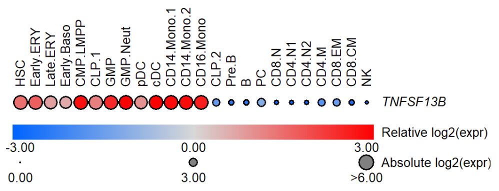

#### B. chr19:17549491[A→G; EAF 46.672%] (rs11666267, cs_129)

Ig lead variant: Same (loc# 463)
Ig associations: IgM↓
Candidate gene: *NIBAN3* (p.Ser572Gly)
Candidate mechanism: Missense variant in *NIBAN3* associated with reduced frequencies of total, transitional, and naïve B cells, as well as lower IgM. NIBAN3 has been identified as a membrane protein on chronic lymphocytic leukemia cells, but its role in human B cell biology remains unclear. In mice, Niban3 deletion enhances B cell proliferation and IgM/IgG3 production by reducing apoptosis, whereas overexpression promotes apoptosis. Thus rs11666267-G likely encodes a gain-of-function variant.

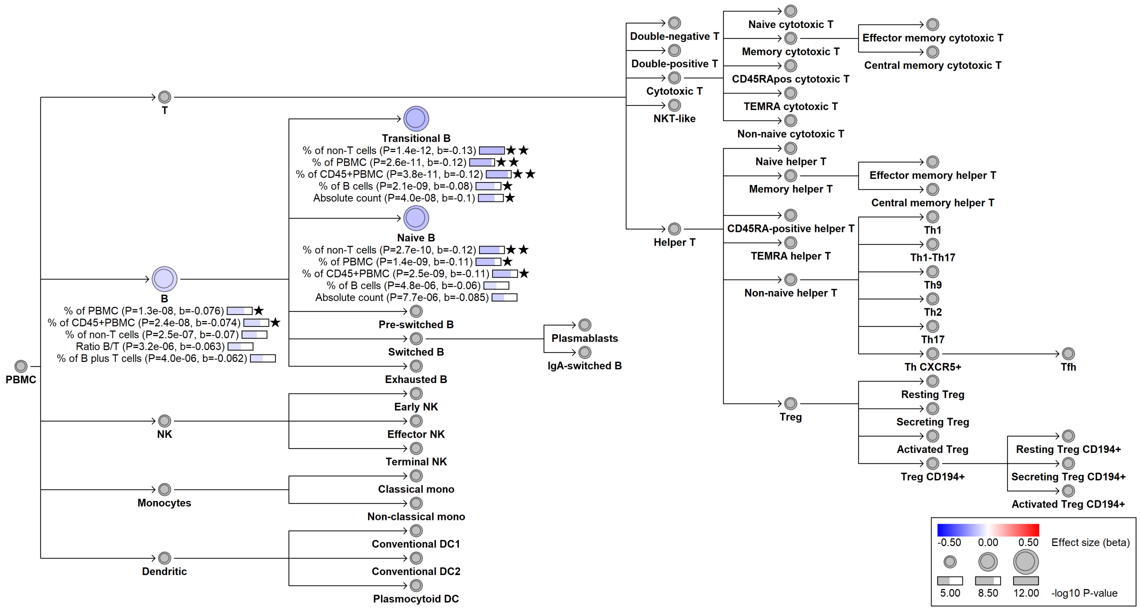


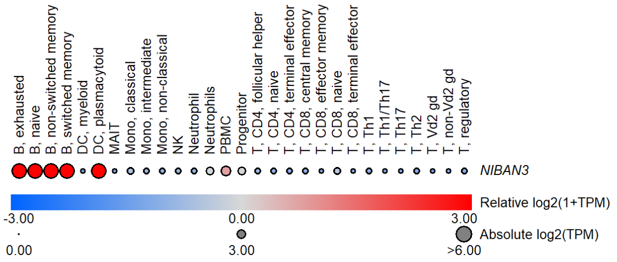


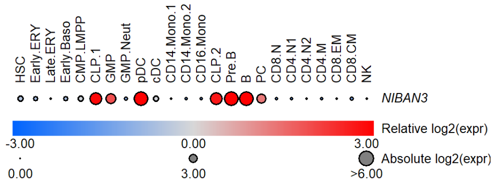

#### C. chr15:40110818[G→GC; EAF 46.946%] (rs36081508, cs_69)

Ig lead variant: chr15:40105735[T→G; 47.9%] (rs539846, loc# 391)
Ig associations: IgA↑, IgA/IgG↑, IgA/IgM↑, IgM/(IgA*IgG)↓
Candidate gene: *BMF*
Candidate mechanism: Increases both IgA and IgA-switched (IgA$^+$) B cell frequency, and downregulates *BMF* in switched memory B cells (P = 3.1e-9, $\beta$ = -0.34 in ImmunExUT), thus increasing the frequency of these cells as BMF is normally pro-apoptotic. This leads to an increase in IgA+ switched B cells and IgA.

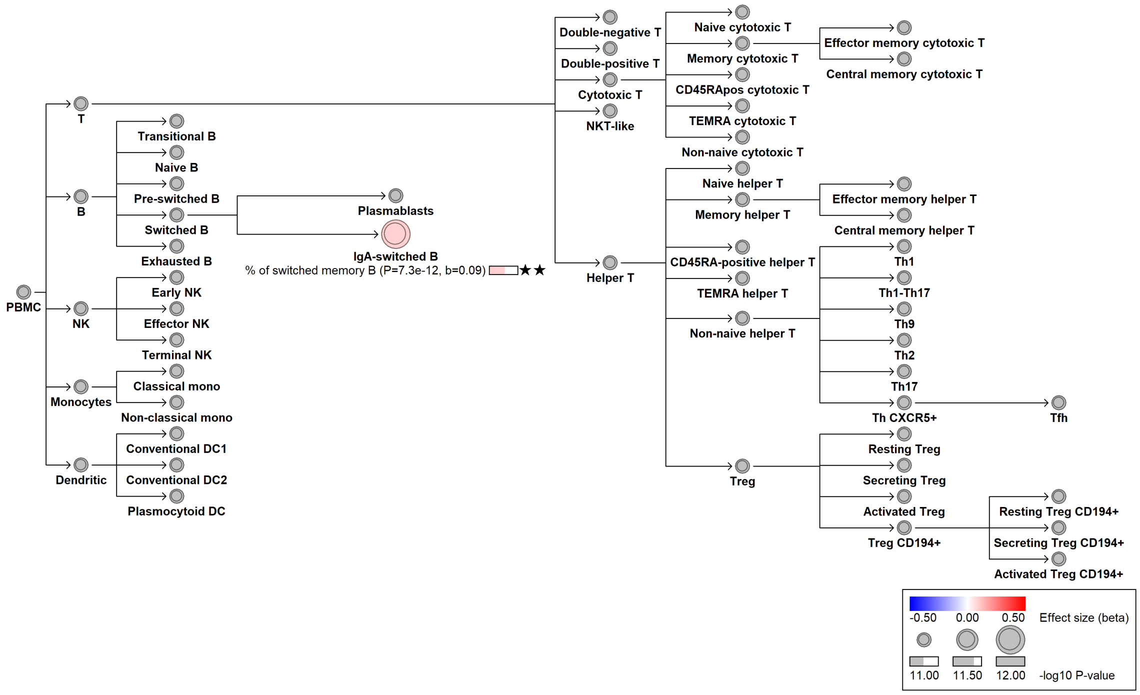


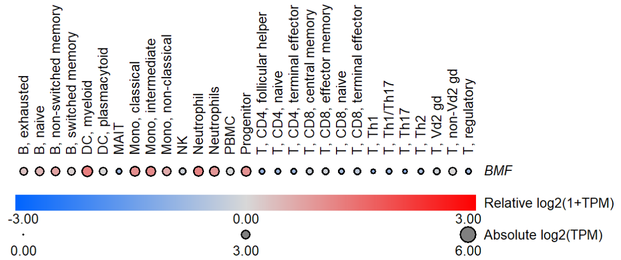


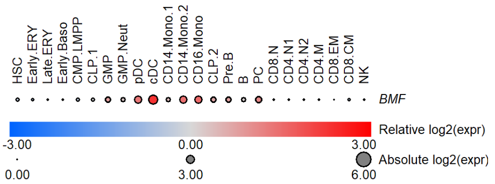

#### D. chr17:16940415[G→T; EAF 0.876%] (rs72553883, cs_214)

Ig lead variant: Same (loc# 424)
Ig associations: IgA*IgG*IgM↓, IgA*IgG↓
Candidate gene: *TNFRSF13B* (p.Ala181Glu)
Diseases associations: Known loss-of-function variant in *TNFRSF13B*, which encodes the TACI receptor, essential for B-cell development beyond the class-switching stage. Consistent with stalled B cell development, BloodVariome shows increased pre-switched B cell frequencies. The variant reduces IgA and IgG levels combined, and is a risk variant for common variable immunodeficiency (CVID).

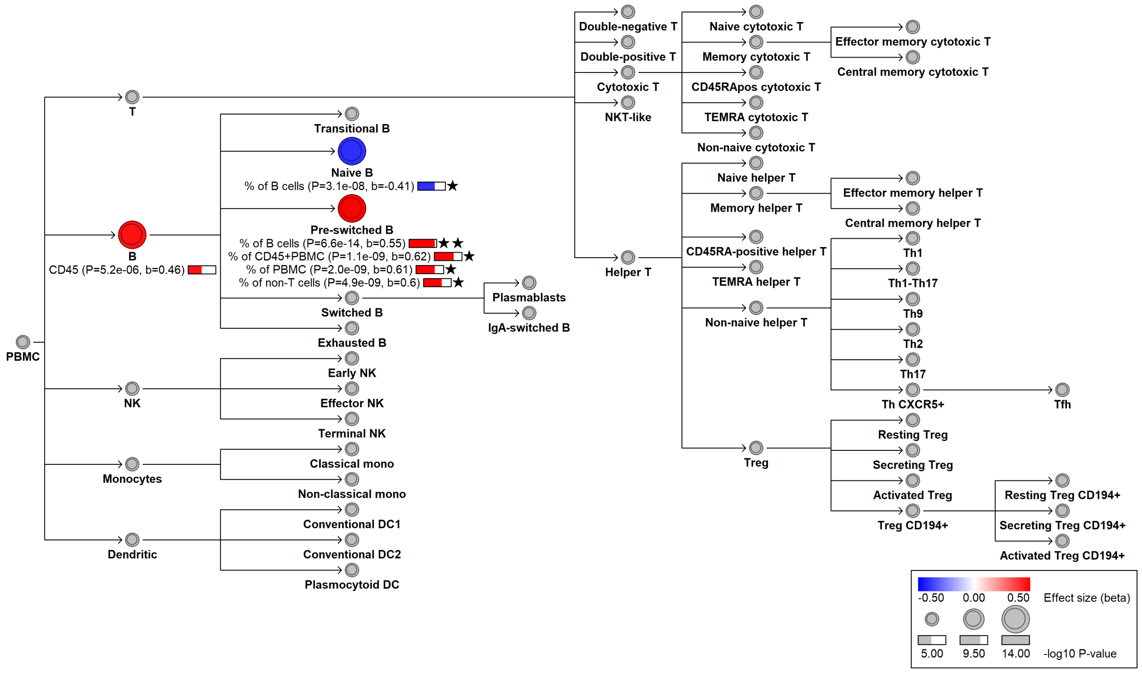


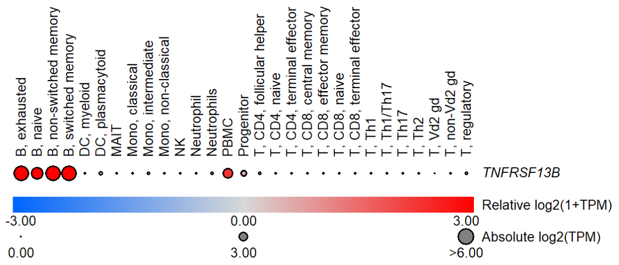


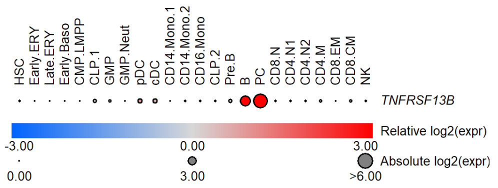

##### E. chr17:16948873[A→G; EAF 0.342%] (rs34557412, cs_117)

Ig lead variant: Same (loc# 426)
Ig associations: IgA↓, IgG↓, IgA*IgG*IgM↓, IgA*IgG↓, IgA/IgG↓, IgM/(IgA*IgG)↑
Candidate gene: *TNFRSF13B* (p.Cys104Arg)
Diseases associations: Known loss-of-function variant in *TNFRSF13B*, which encodes the TACI receptor, essential for B-cell development beyond the class-switching stage. Consistent with stalled B cell development, BloodVariome shows increased pre-switched B cell frequencies. The variant reduces IgA and IgG levels combined, and is a known risk variant for common variable immunodeficiency (CVID).

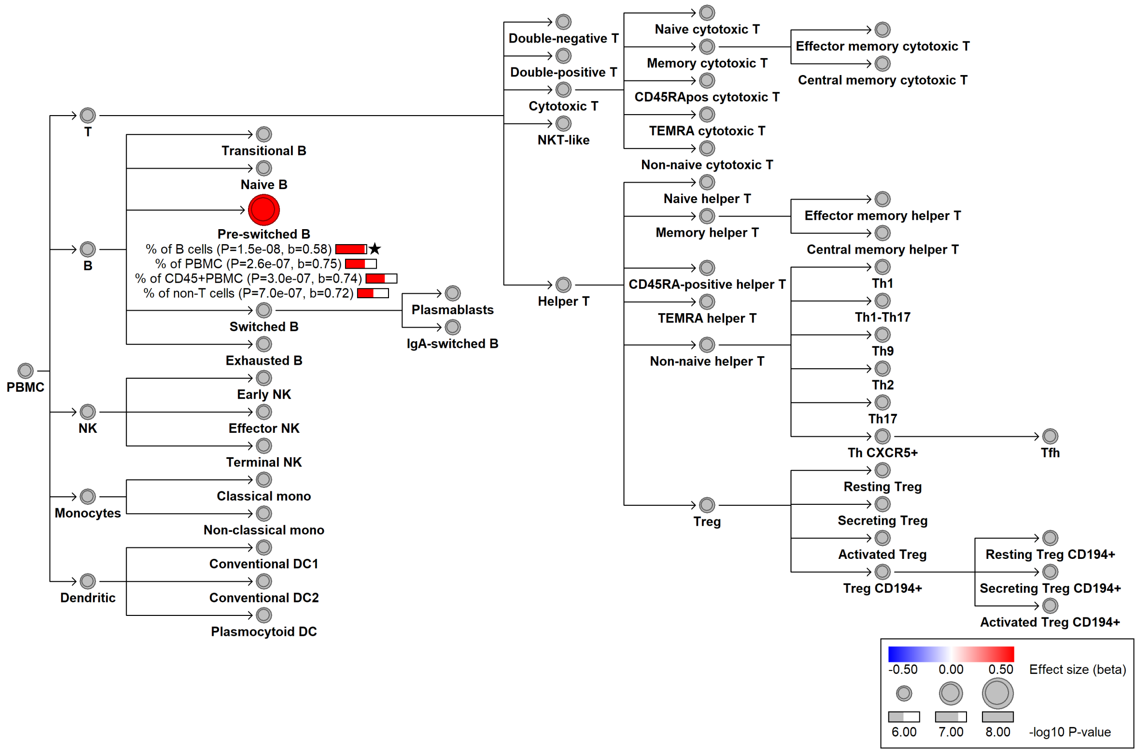


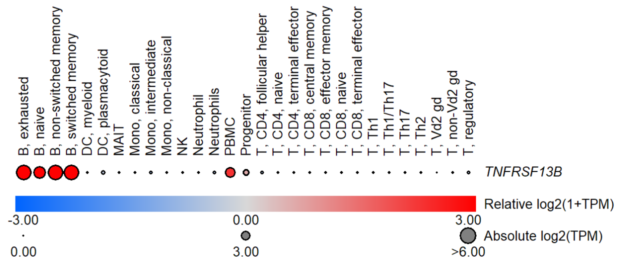


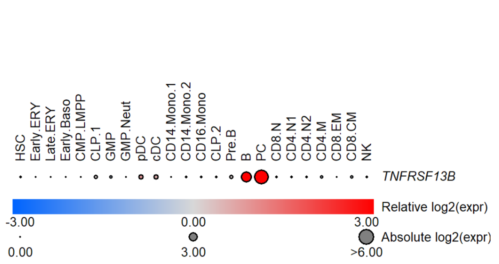

#### F. chr1:24965206[C→T; EAF 1.021%] (rs188468174, cs_116)

Ig lead variant: Same (loc# 9)
Ig associations: IgA↓, IgA*IgG*IgM↓, IgA*IgG↓, IgA/IgG↓, IgA/IgM↓, IgM/(IgA*IgG)↑
Candidate gene: *RUNX3*
Candidate mechanism: Rare *RUNX3* promoter variant with a strong negative effect on IgA. Shifts long-vs-short transcript isoform proportions (PMID 28628107). BloodVariome reveals an association with increased plasmablast frequency, suggesting a compensatory effect due to reduced plasma cell function and/or impaired plasmablast homing.

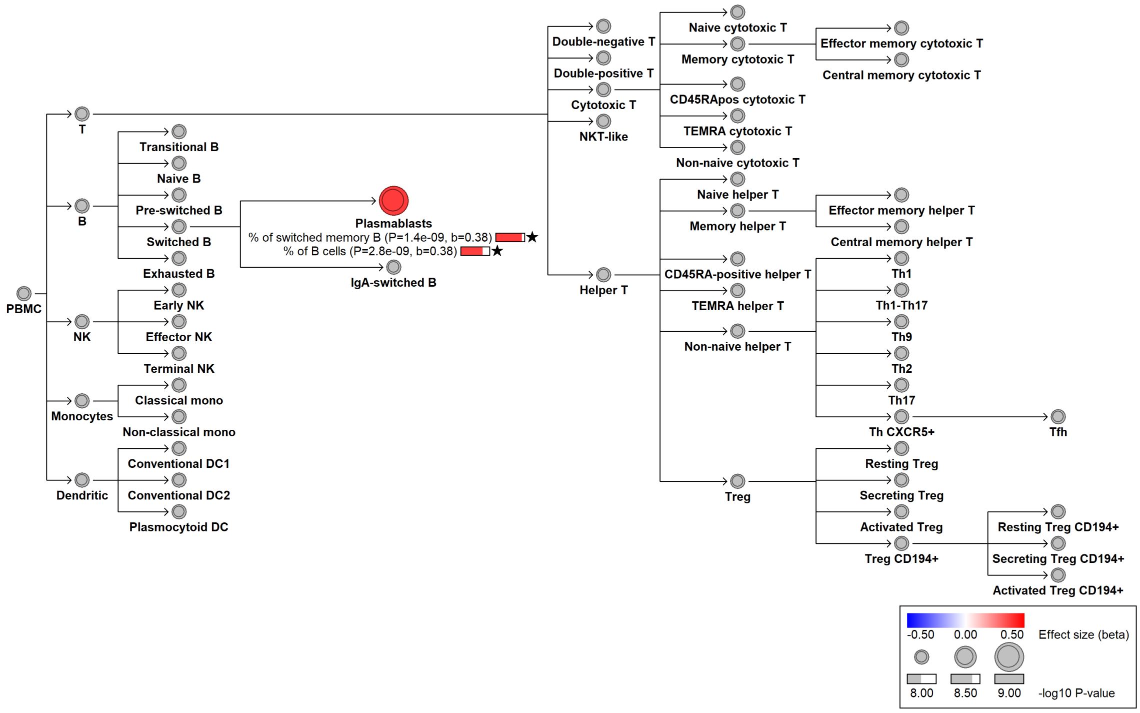


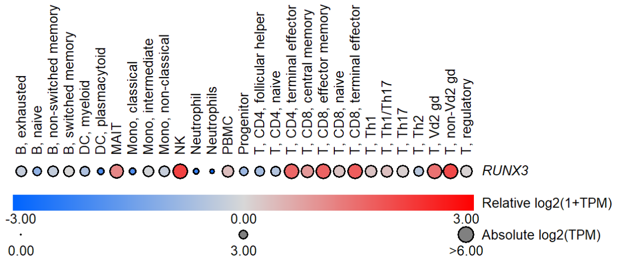


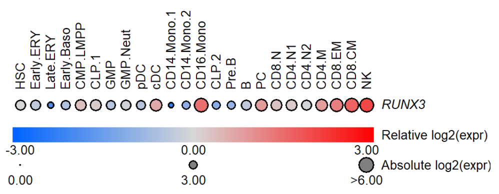

#### G. chr17:63937688[G→T; EAF 36.758%] (rs4968674, cs_25)

<u>Ig lead variant</u>: chr17:63942988[C→T; EAF 41.93%] (rs2058194, loc# 444)
<u>Ig associations</u>: IgA↓, IgG↓, IgA/IgG↑
<u>Candidate gene</u>: *CD79B*
<u>Candidate mechanism</u>: IgD, together with *CD79A* and *CD79B*, form the B-cell receptor (BCR) complex, critical for antigen recognition and B-cell activation. The variant decreases IgA and IgG, and shows a coincidental association with reduced *CD79B* expression in naive and unswitched B-cells in the ImmunExUT compendium (P = 6.4e-6 and 9.6e-5, respectively; PMID 33930287), consistent with reduced BCR function.

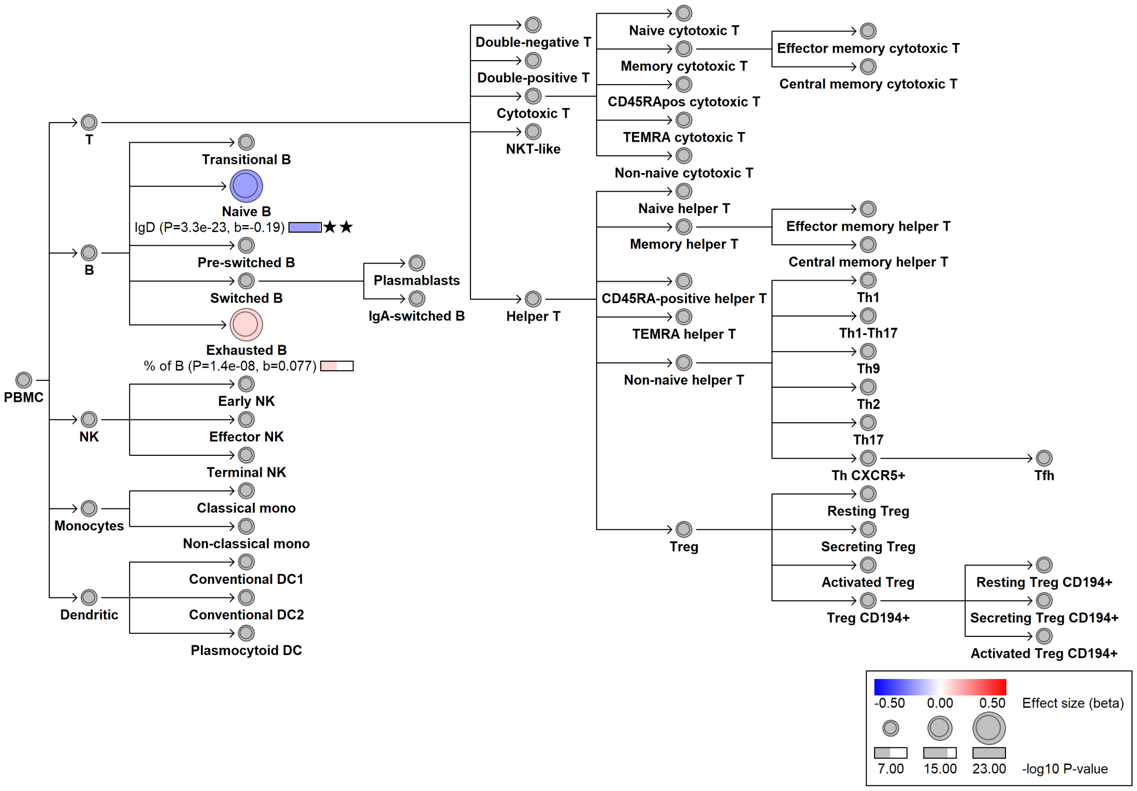


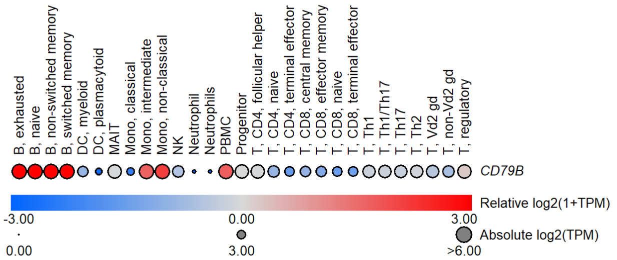


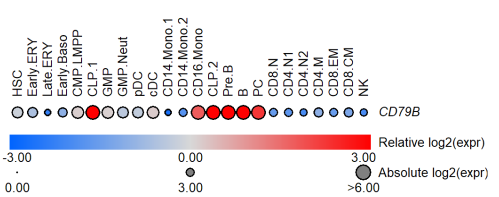

##### H. chr15:40624039[A→G; EAF 40.19%] (rs17747633, cs_115)

Ig lead variant: chr15:40563011[T→A; EAF 43.7%] (rs10152546, loc# 392)
Ig associations: IgM↓, IgG/IgM↑
Candidate gene: *CCDC32* (p.Lys2Ile and eQTL)
Candidate mechanism: Downregulates *CCDC32* in naive B cells (P = 9.2e-8, $\beta$ = -0.53 in ImmunExUT). Additionally, rs10152546 encodes a missense variant (AlphaMissense 0.37). *CCDC32* is involved in clathrin-mediated endocytosis. BloodVariome shows altered IgD (B-cell receptor) expression on transitional and naive B cells; and altered frequencies of total, IgA-switched, and exhausted B cells.

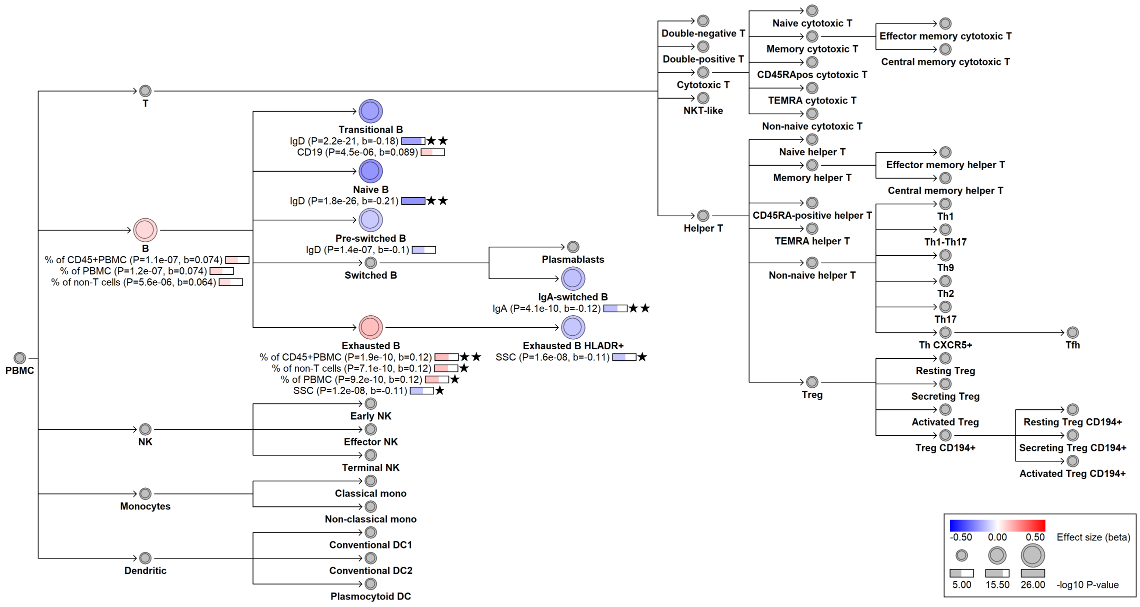


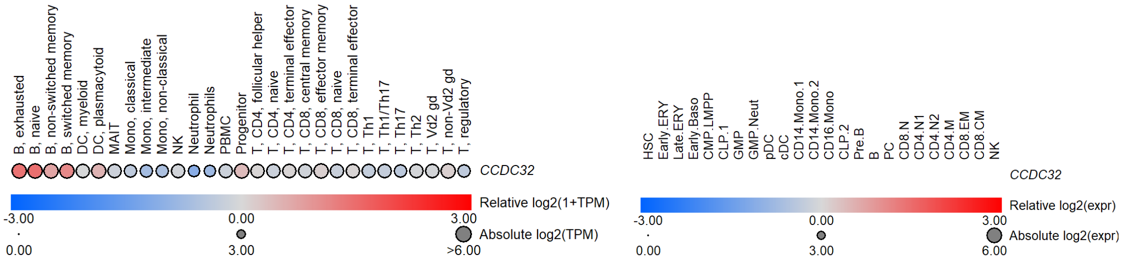

#### I. chr7:2256917[A→G; EAF 0.517%] (rs144787122, cs_62)

Ig lead variant: Same (loc# 260)
Ig associations: IgG↓
Candidate gene: *SNX8* (p.Ile414Thr)
Candidate mechanism: Rare missense variant in *SNX8* with strong negative effects on IgG levels and IgD expression on naive and transitional B cells. IgD is a key component of the B-cell receptor complex. *SNX8* encodes a sorting nexin involved in endosomal sorting and protein trafficking. Many sorting nexins, including SNX8, contain a BAR domain required for homodimerization and membrane curvature sensing. The Ile414Thr variant maps to this domain and predicted deleterious (AlphaMissense score 0.77). *SNX8* is preferentially expressed in B cells, but its functional role in humoral immunity remains unclear.

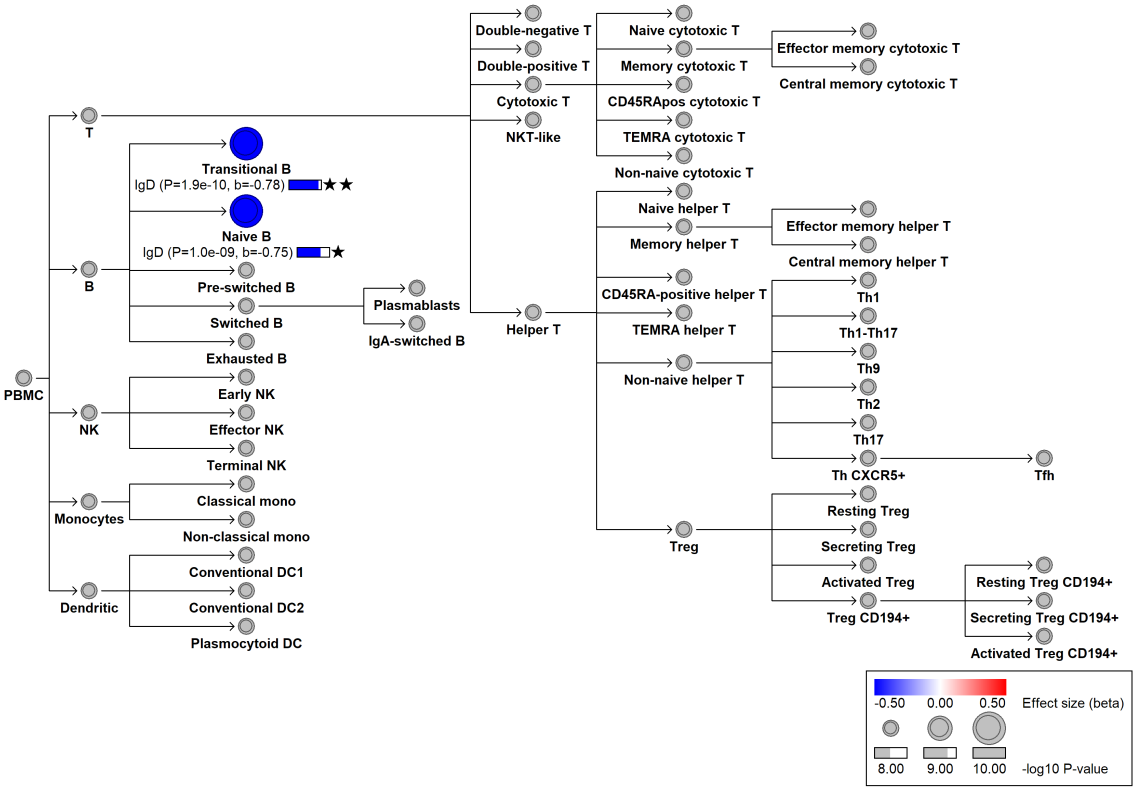


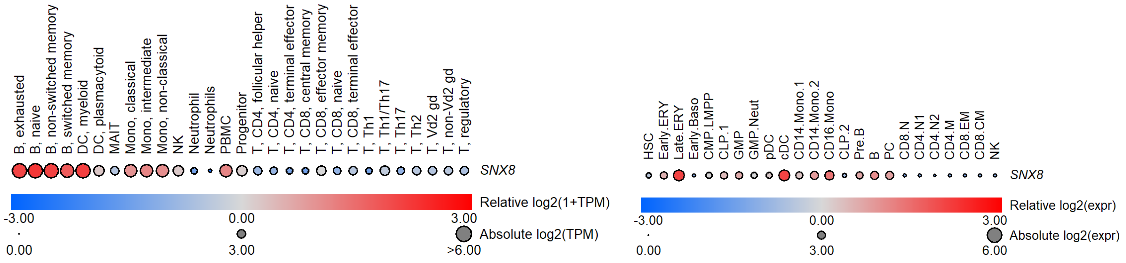

#### J. chr2:60356235[A→G; EAF 3.196%] (rs76529516, cs_114)

Ig lead variant: Same (loc# 69)
Ig associations: IgM↓, IgA/IgG↓, IgG/IgM↑
Candidate gene: *BCL11A* (nearest gene)
Candidate mechanism: *BCL11A* is essential for lymphoid development and deletion leads to apoptosis of early B cells in animal models (PMID 23230003). BloodVariome revealed decreased CD19 expression on exhausted B cells and a borderline association with higher transitional B cell frequencies, suggesting delayed B-cell development.

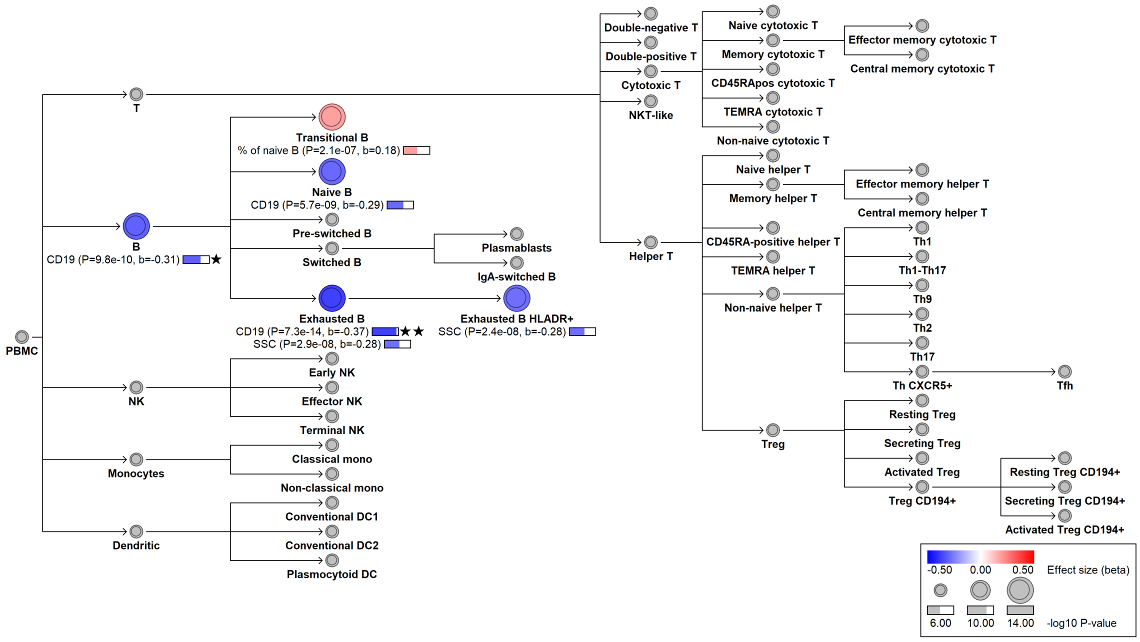


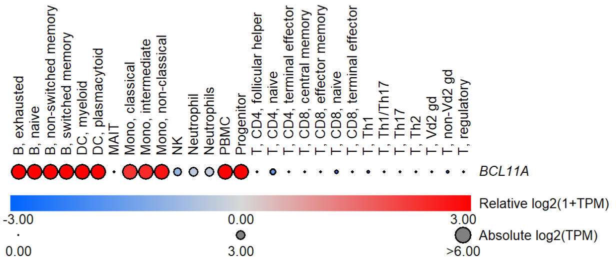


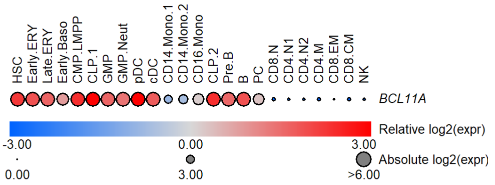

#### K. chr1:45331833[C→G; EAF 24.413%] (rs3219489, cs_171)

Ig lead variant: Same (loc# 13)
Ig associations: IgA↓, IgM↓, IgA/IgM↑, IgG/IgM↓, IgM/(IgA*IgG)↑
Candidate gene: *TESK2* (eQTL in B cells), *MUTYH* (p.Gln335His)
Candidate mechanism: Decreases CD19 surface expression on pre-switched B cells.

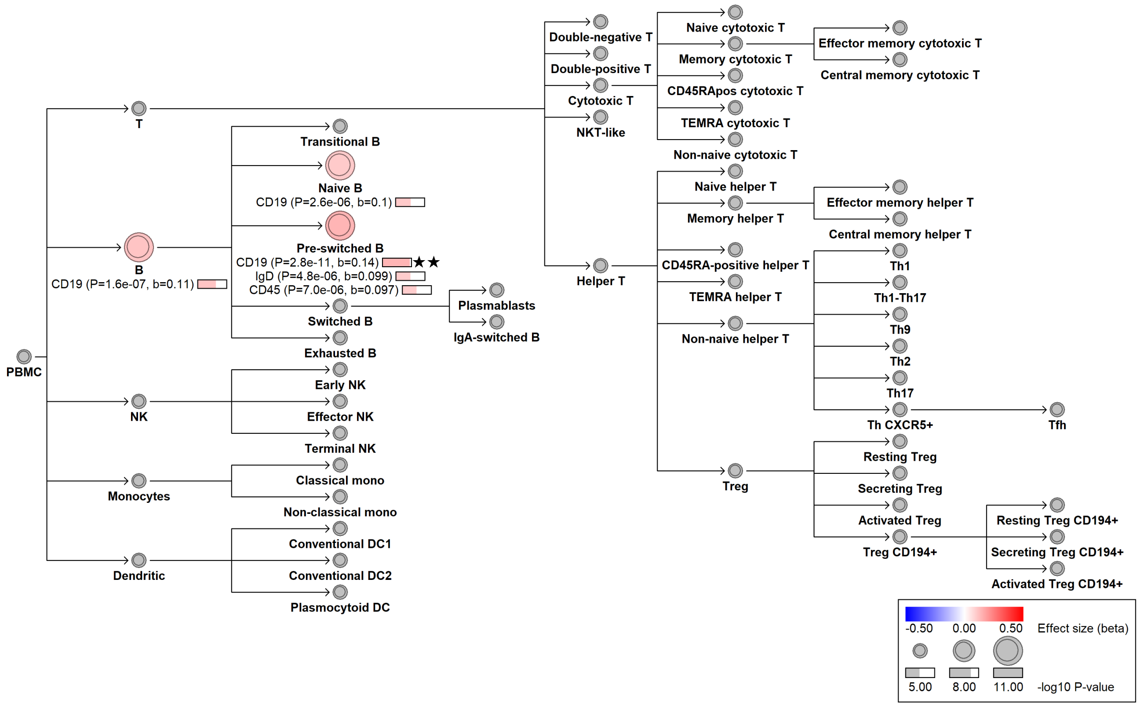


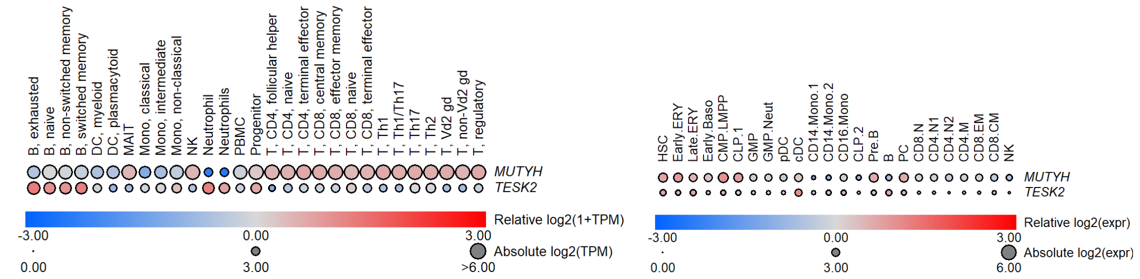

#### L. chr12:111446804[C→T; EAF 46.218%] (rs3184504, cs_194)

Ig lead variant: Same (loc# 355)
Ig associations: IgA↑, IgG↑, IgA*IgG*IgM↑, IgA*IgG↑, IgA/IgG↑, IgA/IgM↑, IgM/(IgA*IgG)↓
Candidate gene: *SH2B3* (p.Trp262Arg)
Candidate mechanism: Well-known, highly pleiotropic variant associated with a wide range of diseases and quantitative traits. Encodes a regulator of cell signaling, downstream of various cytokine and hormone receptors.

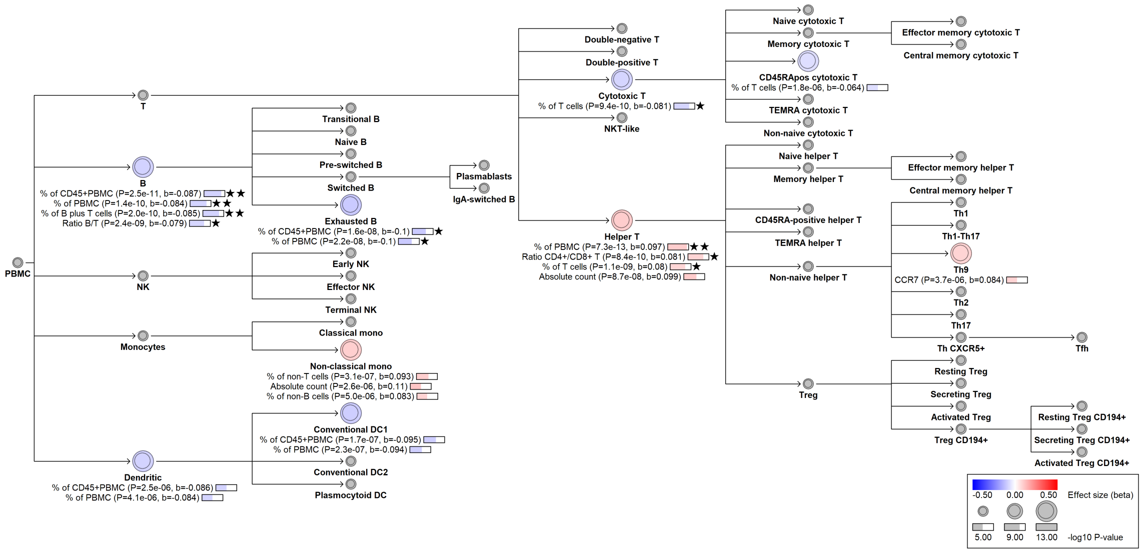


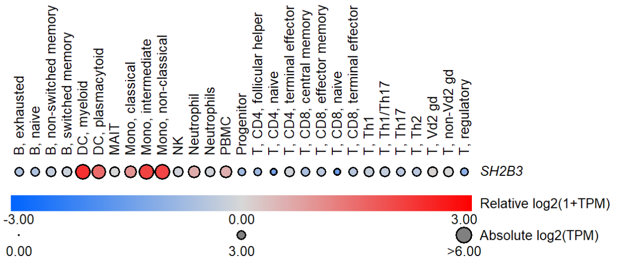


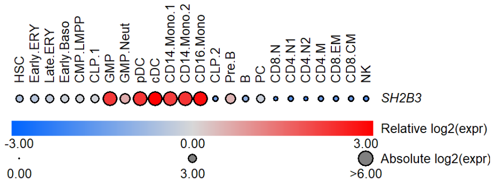

#### M. chr1:200908722[!→G; EAF 27.404%] (rs41299637, cs_141)

Ig lead variant: chr1:200911850[T→C; 30.1%] (rs296520, loc# 61)
Ig associations: IgA↓, IgA/IgG↓, IgA/IgM↓, IgM/(IgA*IgG)↑
Candidate gene: *INAVA* (p.Cys453Arg)
Candidate mechanism: Protective against Inflammatory Bowel Disease, Celiac disease, Crohn's disease, Ulcerative colitis, Ankylosing spondylitis, Multiple sclerosis, and Psoriasis. Decreases IgA. *INAVA* is expressed in gut mucosa and maintains intestinal barrier integrity (PMID 30355448). Consistent with the decreased risk of inflammatory bowel disease and decreased IgA, we find decreased frequencies of Th9 cells, which contribute to inflammatory diseases, including inflammatory bowel disease (PMID 26885239, 29881387).

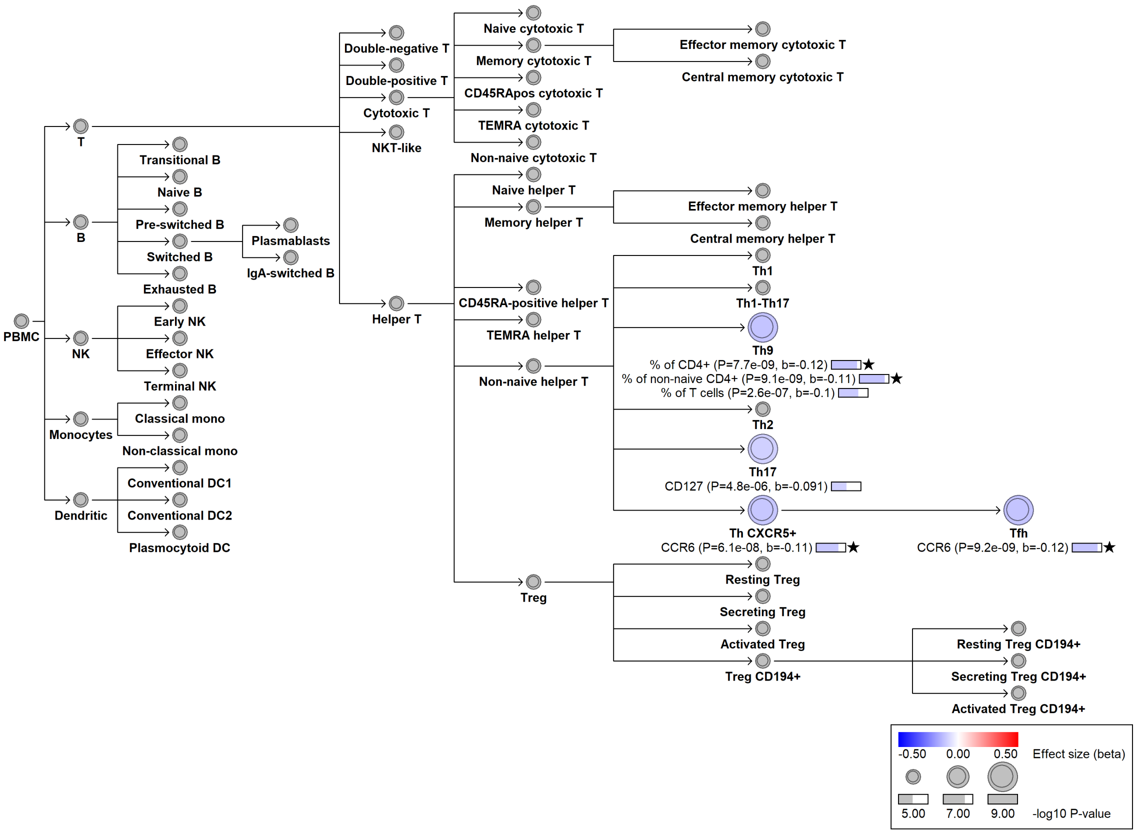


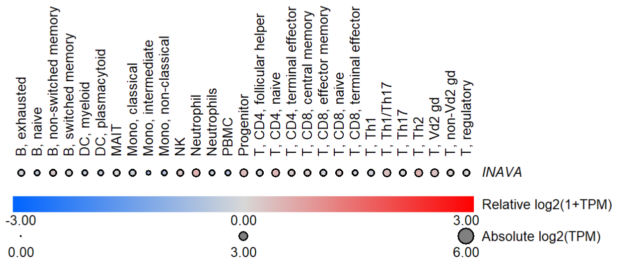


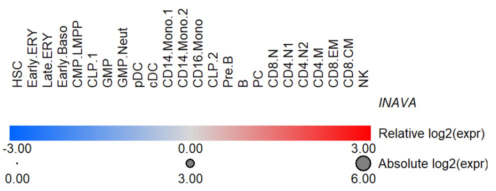

#### N. chr10:6052734[C→T; EAF 7.707%] (rs61839660, cs_101)

Ig lead variant: Same (loc# 305)
Ig associations: IgA↑, IgA*IgG↑, IgA/IgG↑, IgA/IgM↑, IgM/(IgA*IgG)↓
Candidate gene: *IL2RA* (eQTL)
Diseases associations: Risk variant for multiple autoimmune and allergic conditions. Increases plasma IgA levels and surface expression of IL2RA (CD25) on helper T cells and regulatory T cells. Additionally, we noted reduced frequencies of resting regulatory T cells.

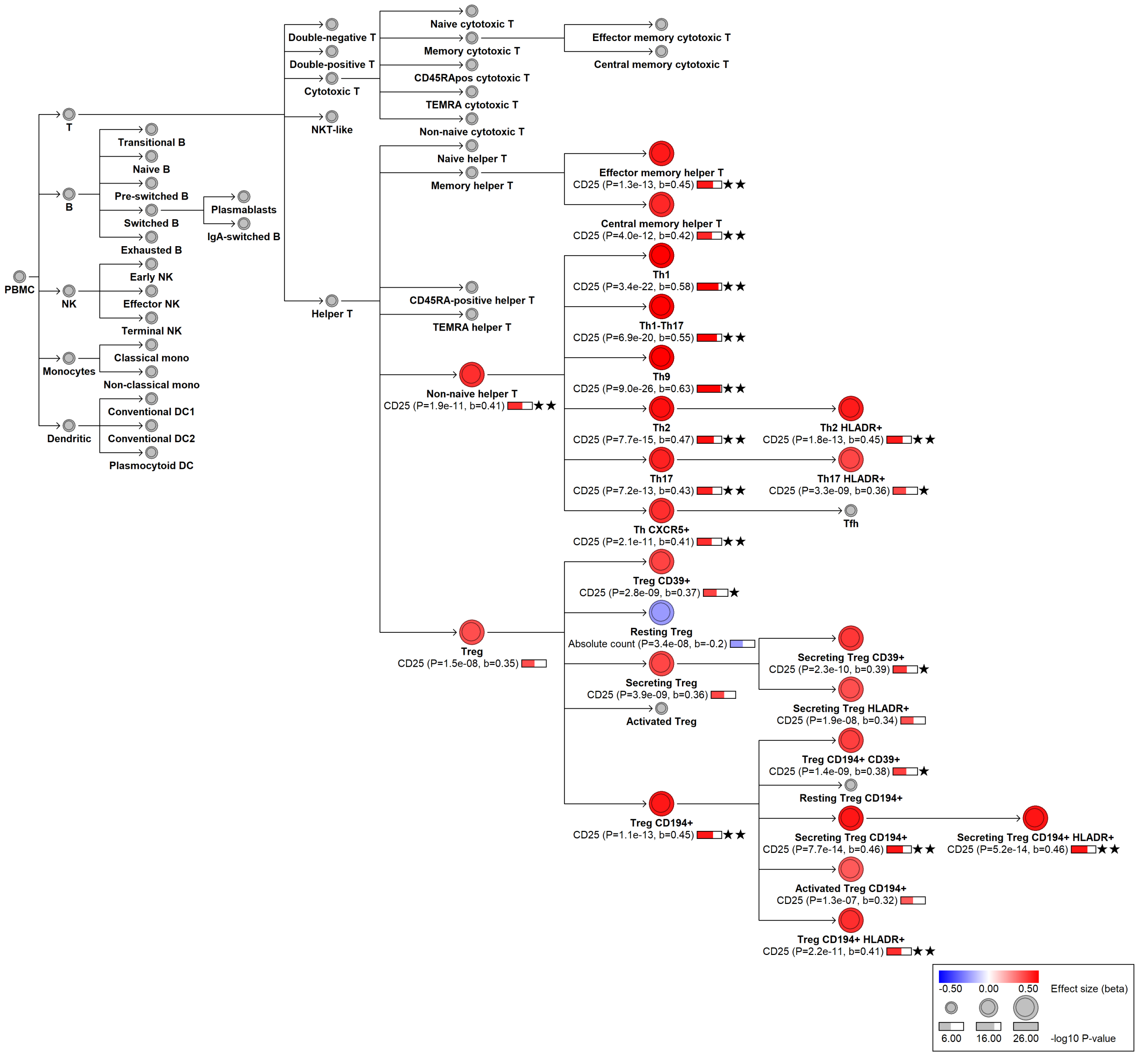


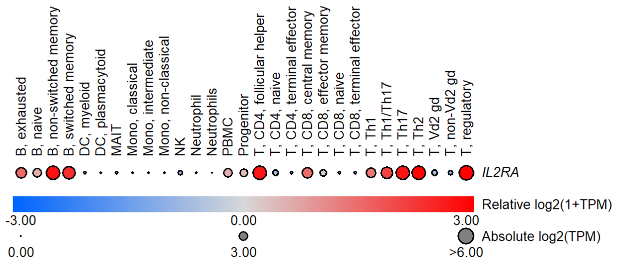


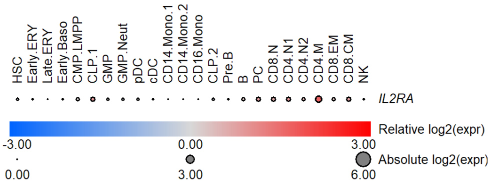

#### O. chr22:30196498[!→C; EAF 46.49%] (chr22:30196498, cs_170)

Ig lead variant: chr22:30181782[G→A; 34.5%] (rs714027, loc# 495)
Ig associations: IgA↓, IgG↓, IgA*IgG*IgM↓, IgA*IgG↓, IgA/IgG↓, IgM/(IgA*IgG)↑
Candidate gene: *ASCC2*, *LIF*, *MTMR3*, *OSM*
Diseases associations: This variant is associated with lower IgA and IgG levels. In Blood-Variome, we see increased expression of CCR7 on helper T cell subsets. The candidate gene assignment at this locus is ambiguous and none of the candidate genes have known connections to CCR7. The variant is also associated with autoimmune diseases.

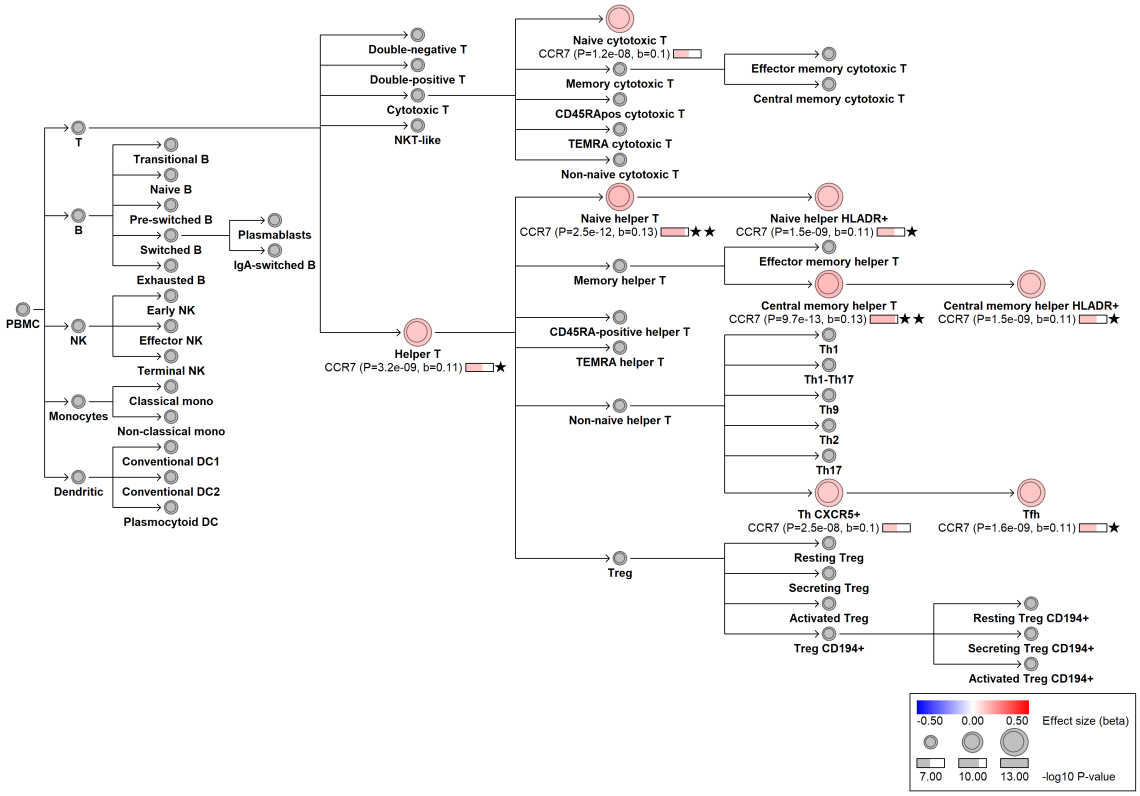

#### P. chr9:114930602[C→T; EAF 21.292%] (rs3181356, cs_218)

Ig lead variant: Same (loc# 299)
Ig associations: IgA↑, IgA*IgG*IgM↑, IgA*IgG↑, IgA/IgG↑, IgA/IgM↑
Candidate gene: *TNFSF8* (eQTL)
Candidate mechanism: Increases IgA and the frequency of IgA-switched B cells. It also affects Treg frequencies and CD25 (IL2RA) expression on Treg subsets. *TNFSF8* encodes CD30 ligand, a cytokine crucial in immune regulation that binds to its receptor CD30 (TNFRSF8), promoting T-cell responses. The variant downregulates *TNFSF8* in B cells, monocytes, neutrophils, and whole blood.

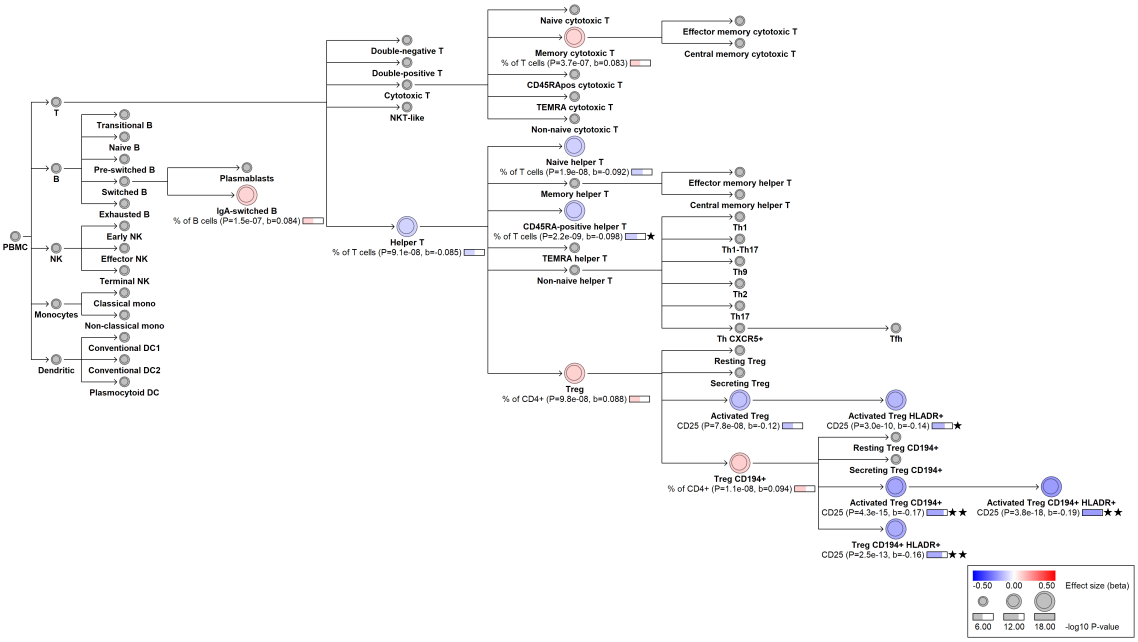


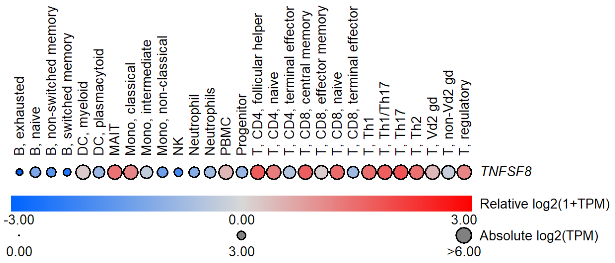


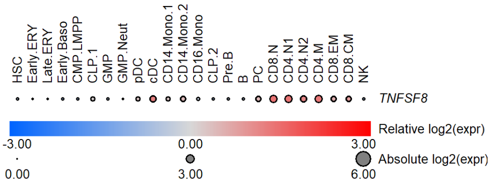

#### Q. chr1:113834946[G→A; EAF 11.711%] (rs2476601, cs_40)

Ig lead variant: Same (loc# 19)
Ig associations: IgM↓, IgA*IgG*IgM↓, IgA/IgM↑, IgG/IgM↑, IgM/(IgA*IgG)↓
Candidate gene: *PTPN22* (p.Trp620Arg)
Candidate mechanism: Known risk variant for multiple autoimmune diseases. We find that the variant reduces IgM. Using BloodVariome, we identify a colocalized association with increased regulatory T cell frequencies and a suggestive association with CD19 expression on transitional and naive B cells.

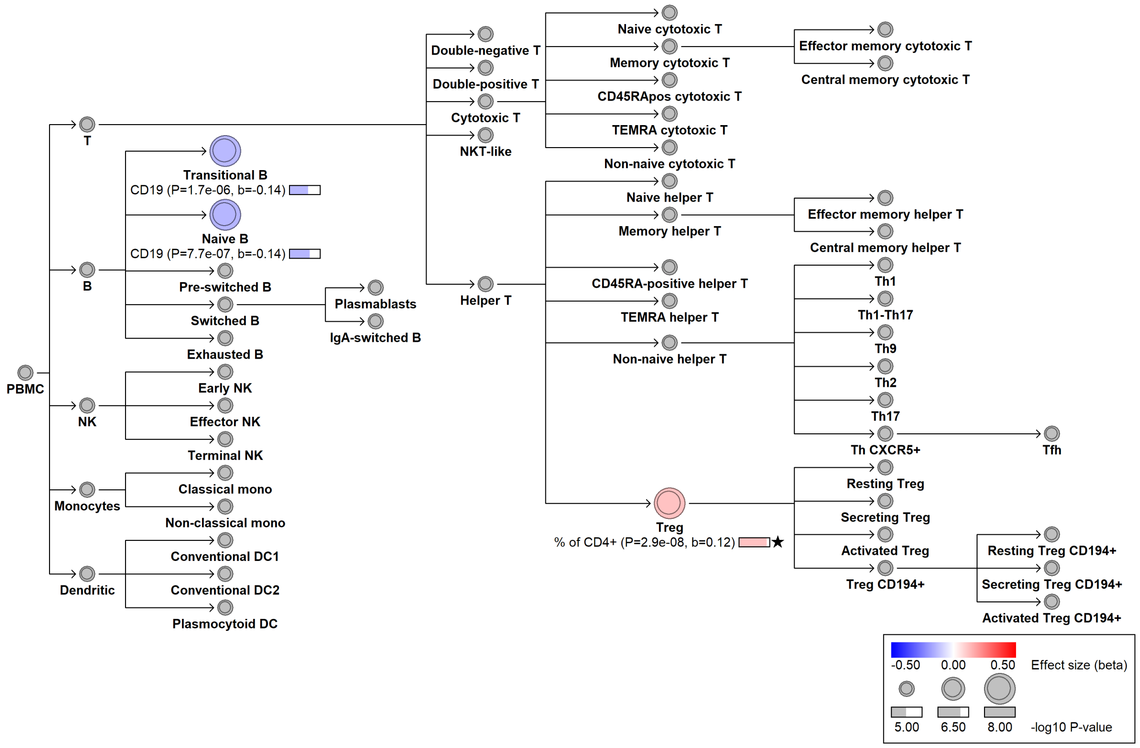


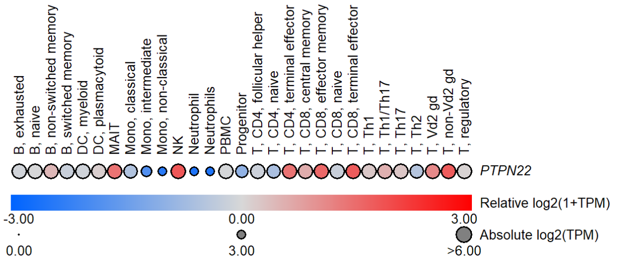


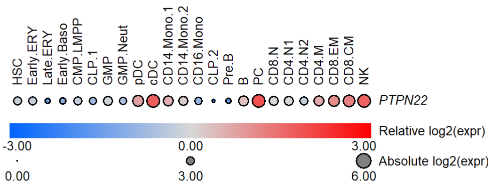

#### R. chr13:28029870[T→C; EAF 1.592%] (rs76428106, cs_3)

Ig lead variant: Same (loc# 357)
Ig associations: IgA↑, IgG↑, IgA*IgG*IgM↑, IgA*IgG↑, IgM/(IgA*IgG)↓
Candidate gene: *FLT3*
Candidate mechanism: The strongest known risk variant for autoimmune thyroiditis; also a risk variant for rheumatoid arthritis and acute myeloid leukemia. Intronic splice variant that causes a premature stop codon in about 30% of transcripts, yet has a net gain-of-function effect (PMID 32581359). Further supporting the gain-of-function model, we find increased frequencies of dendritic cells, particularly conventional type 2, along with increased monocyte frequencies and increased plasma IgA and IgG levels.

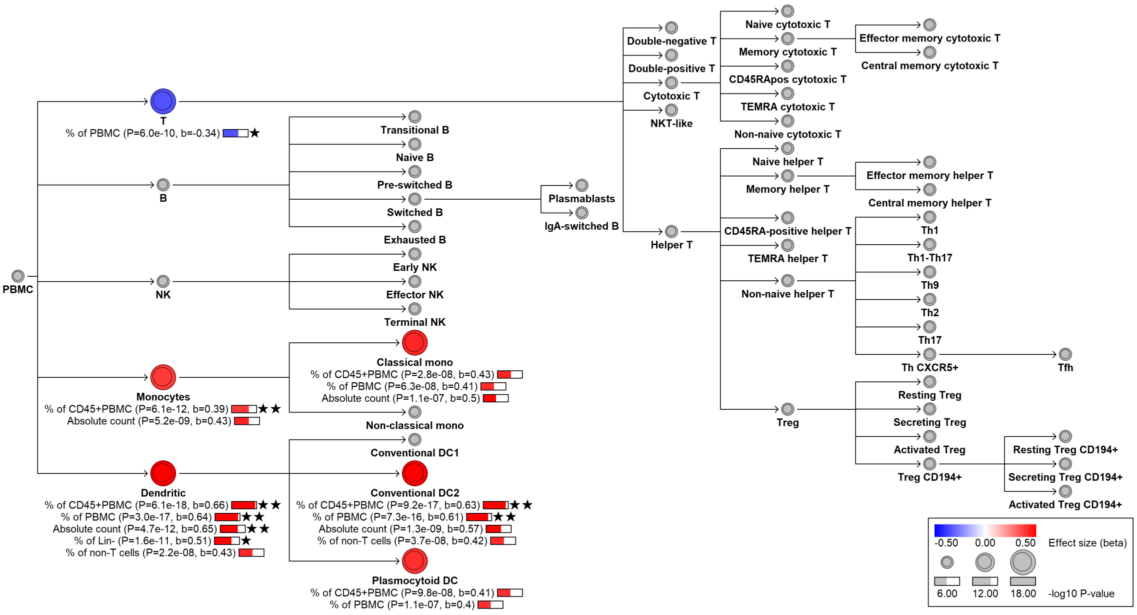


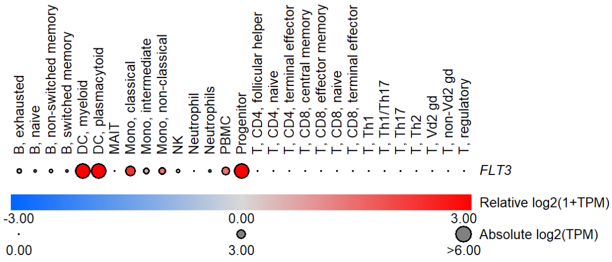


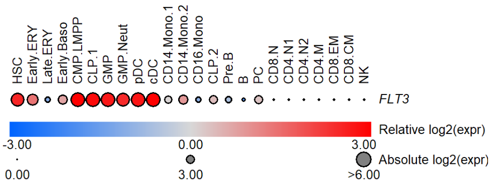

#### S. chr7:50268931[C→T; EAF 29.735%] (rs876038, cs_210)

Ig lead variant: chr7:50268114[T→C; EAF 27.04%] (rs876036, loc# 265)
Ig associations: IgA↓, IgG↓, IgA*IgG*IgM↓, IgA*IgG↓, IgA/IgM↓, IgM/(IgA*IgG)↑
Candidate gene: *IKZF1* (eQTL)
Candidate mechanism: *IKZF1* encodes a transcription factor expressed in several immune cell types. It is classically associated with B cell development, but was recently connected also to human dendritic cell development (PMID 29588478). This variant reduces IgA and IgG, and shifts the dendritic cell composition from plasmacytoid to conventional type 2.

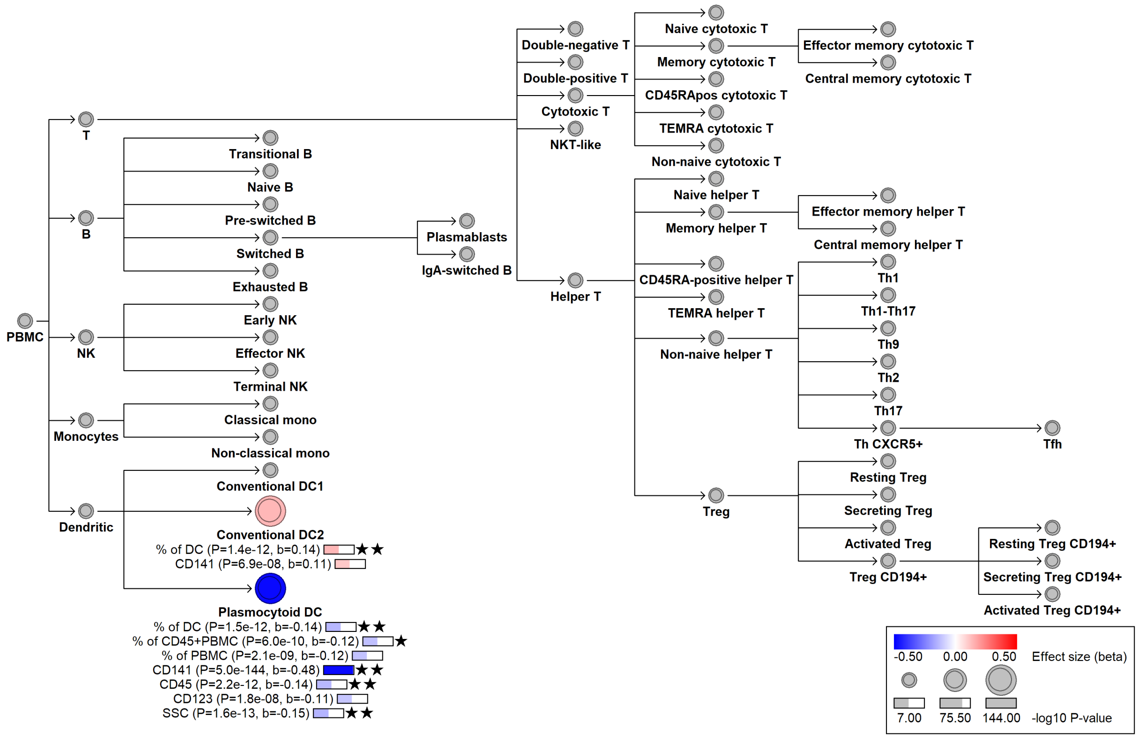


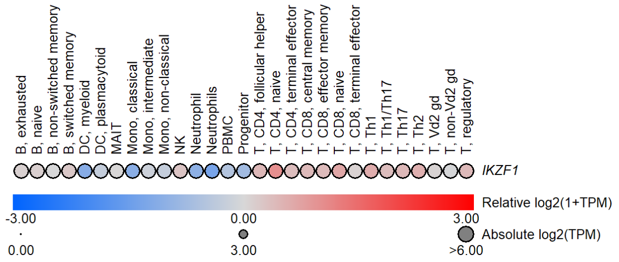


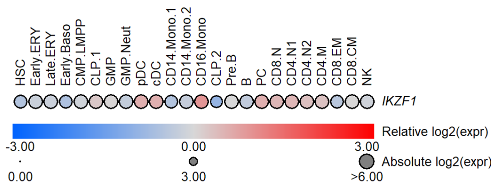

#### T. chr6:90271938[C→T; EAF 31.239%] (rs45553631, cs_191)

Ig lead variant: chr6:90296508[G→A; EAF 33.55%] (rs1504215, loc#249)
Ig associations: IgA/IgM↑, IgM/(IgA*IgG)↑
Candidate gene: *BACH2* (eQTL in memory Treg)
Diseases associations: Increases IgA relative to IgM. Reduces conventional dendritic cells type 2. *BACH2* is a transcription factor expressed in several blood and immune cell types. Associates with Allergy, Ankylosing spondylitis, Asthma, Chronic obstructive pulmonary disease, Crohn’s disease, Eczema, Inflammatory bowel disease, Nasal polyps, Primary sclerosing cholangitis, Psoriasis, and Ulcerative colitis.

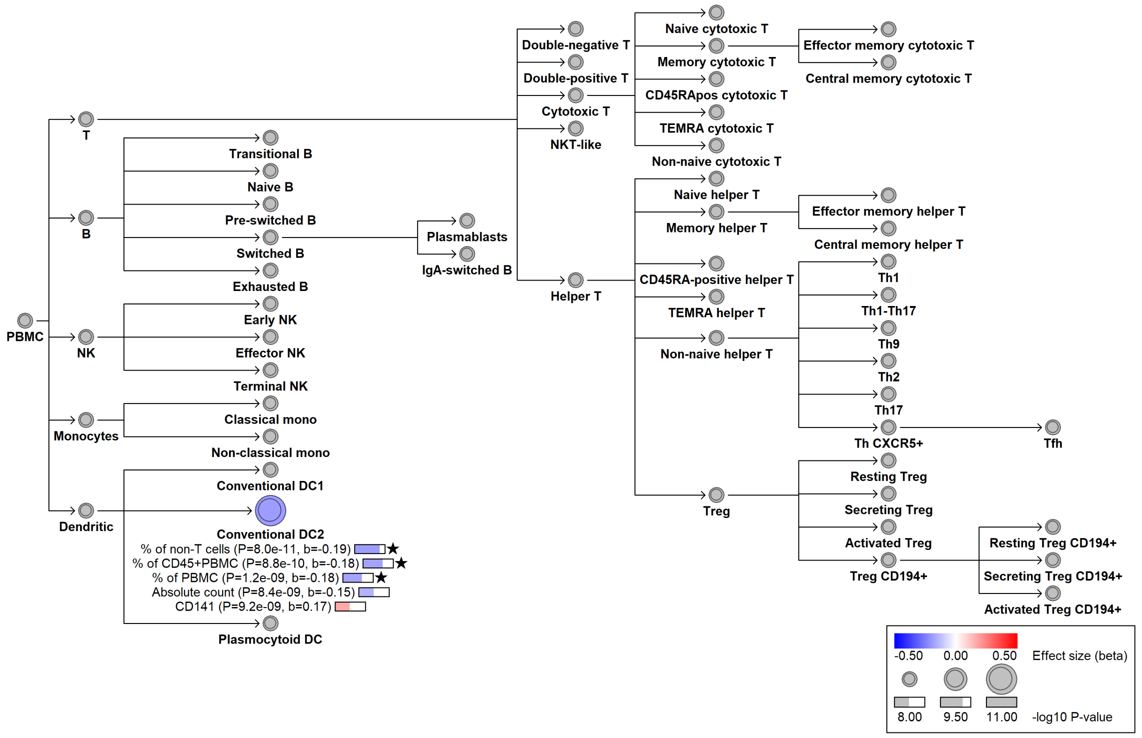


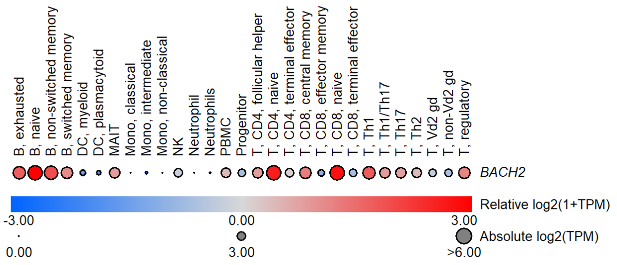


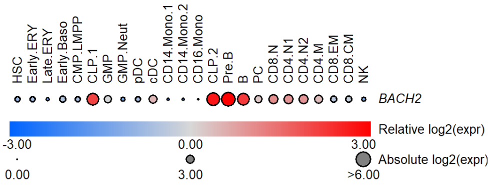

#### U. chr16:88978825[C→T; EAF 15.329%] (rs62045817, cs_144)

Ig lead variant: Same (loc# 415)
Ig associations: IgM↓, IgA*IgG*IgM↓
Candidate gene: *CBFA2T3* in BloodVariome; *SPG7* in Ig GWAS.
Candidate mechanism: Unclear candidate gene and mechanism. Decreases IgM and total Ig levels, along with side scatter in plamacytoid dendritic cells.

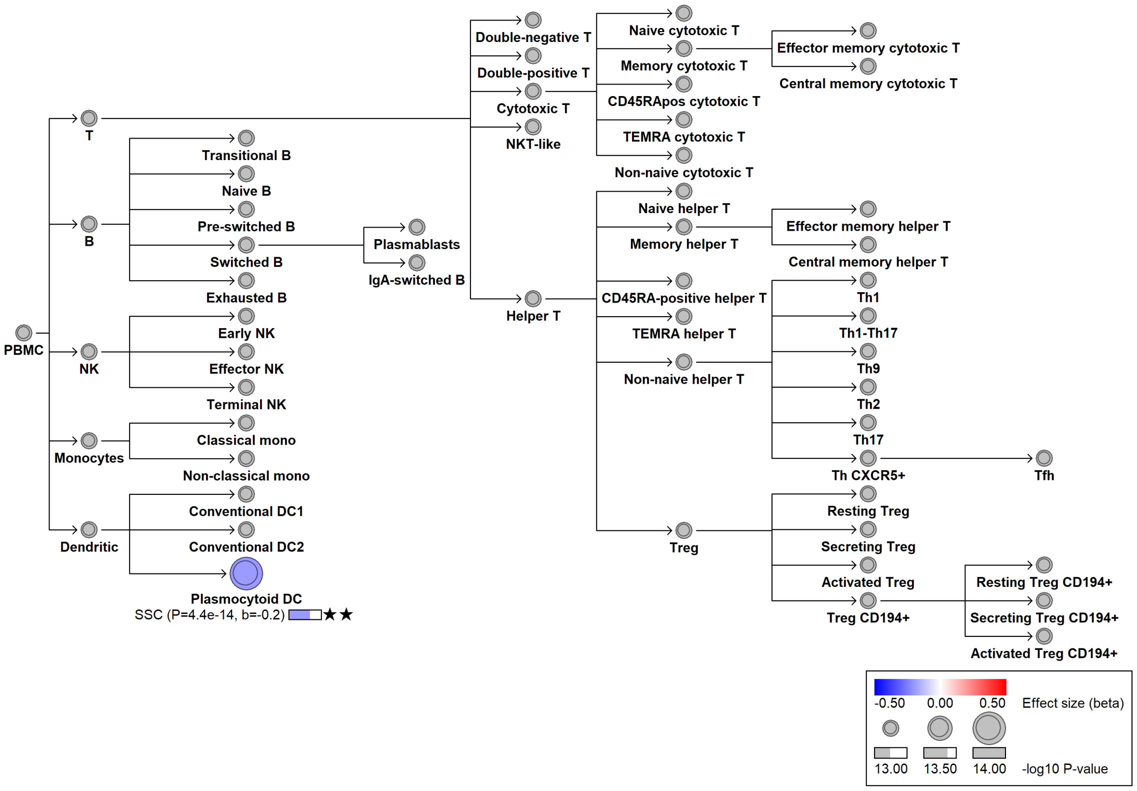

#### V. chr3:47240813[A→!; EAF 39.95%] (rs2276853, cs_98)

Ig lead variant: Same (loc# 106)
Ig associations: IgG↑
Candidate gene: *NBEAL2* in BloodVariome; *KIF*, *SCAP9* in Ig GWAS.
Diseases associations: Upregulates the thrombomodulin receptor (CD141) expression on conventional dendritic cells type 1. Unclear candidate gene.

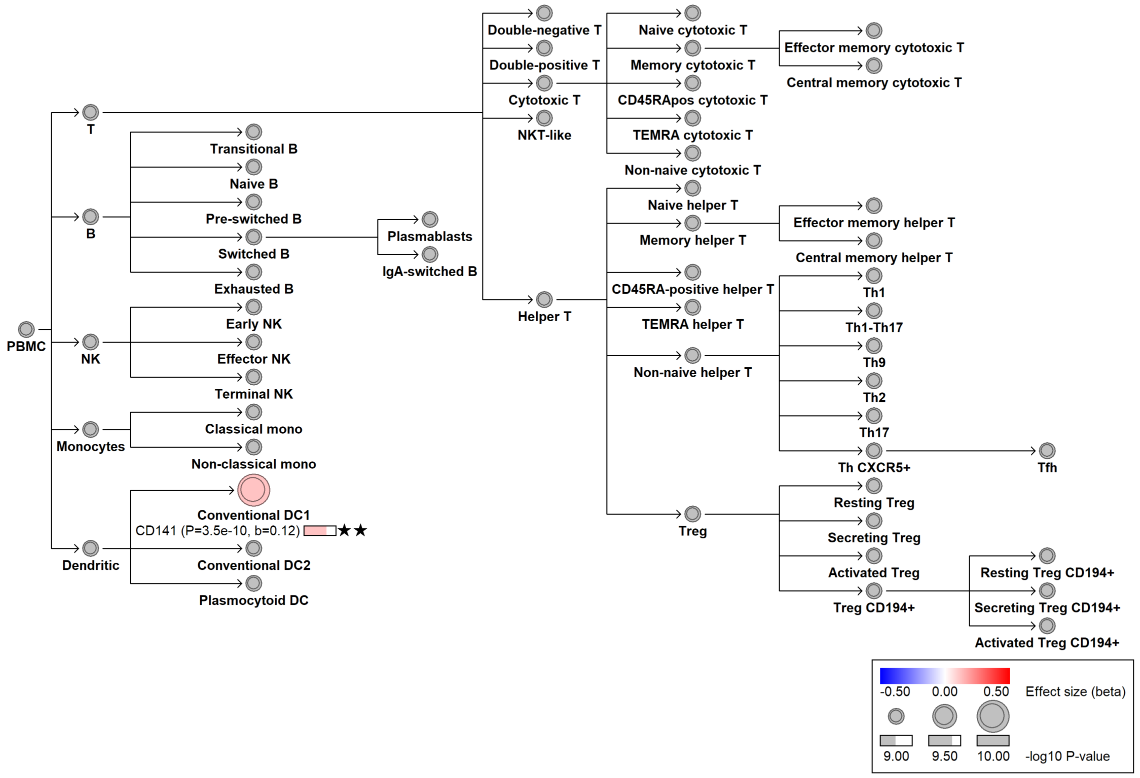

### Figure S6

Expression of *BMF* by rs539846 genotype across immune cell subsets. The IgA-increasing G allele downregulates *BMF* in switched memory B cells. y-axes indicate normalized gene expression; x-axes indicate genotype with sample sizes shown below each box. All data are from ImmuNexUT.

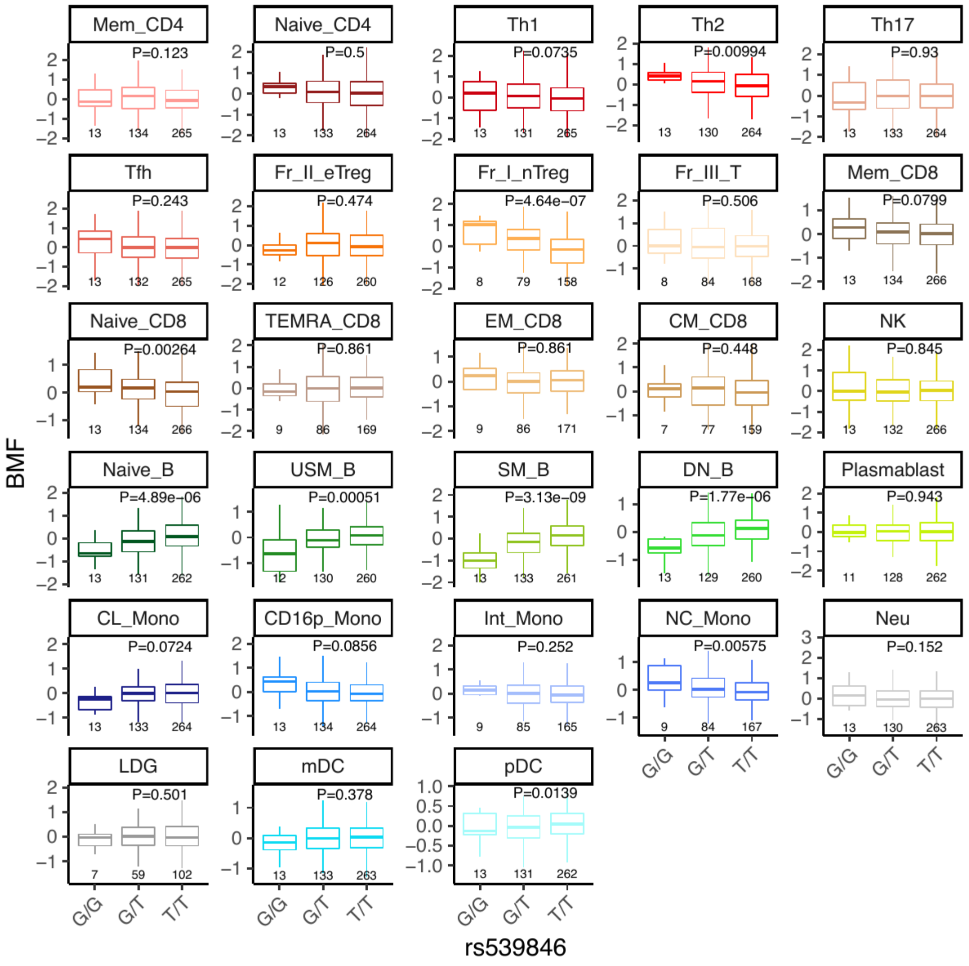

### Figure S7

**(A)** Expression of *ZNF608* by rs12517864 genotype across immune cell subsets. The IgM-decreasing A allele upregulates *ZNF608* in B cell subsets. **(B)** Expression of *MCTP2* by rs6497185 genotype across immune cell subsets. The IgM-decreasing C allele upregulates *MCTP2* in B cell subsets, particularly unswitched memory B cells. **(C)** Expression of *VDR* by rs886441 genotype across immune cell subsets. The M/AG-decreasing G allele downregulates *VDR* in plasmablasts. y-axes indicate normalized gene expression; x-axes indicate genotype with sample sizes shown below each box. All data are from ImmuNexUT. **(D)** ChIP-seq tracks at the *VDR* locus showing IRF4 and BLIMP1 occupancy in plasmablasts derived from memory B cells, and H3K27Ac marking active chromatin. The vertical line indicates the position of rs886441. Peripheral blood memory B cells were differentiated to plasmablasts *ex vivo* via CD40L/anti-BCR/IL-2/IL-21 stimulation (Day 0–3) followed by signal withdrawal into IL-2/IL-21 conditions (Day 3–6). Data from GEO: GSE142493 (PMID: 32843533).

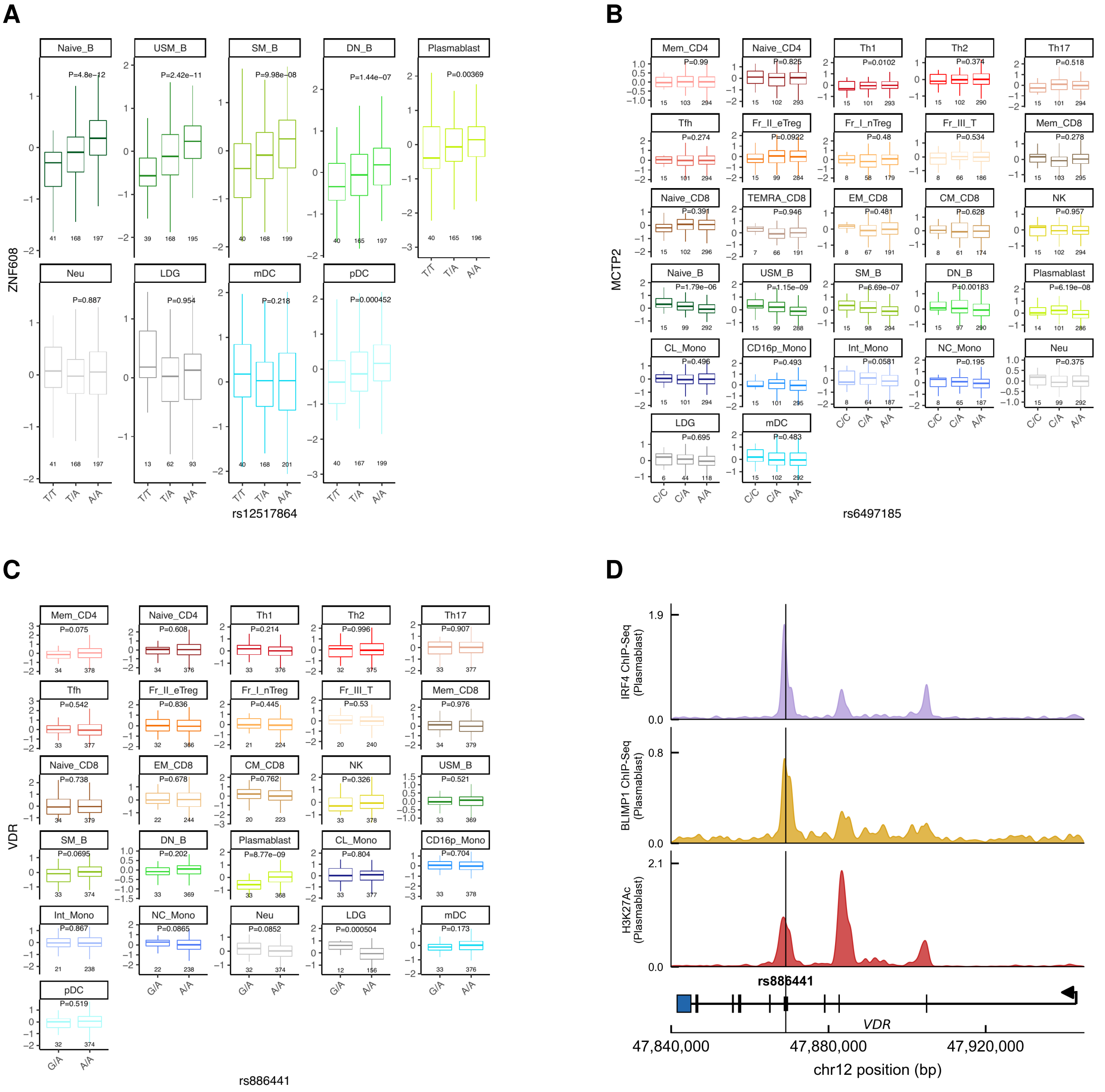

### Figure S8

**(A)** Expression of *ARHGAP15* by rs140397066 genotype in bulk B cells. The IgA-decreasing G allele reduces *ARHGAP15* expression. Data from deCODE. **(B)** Single-cell eQTL showing reduced *ARHGAP15* expression in B cell clusters for rs140397066-G carriers. Data from deCODE. **(C)** Chromatin accessibility at the *ARHGAP15* locus in memory B cells and plasmablasts, with positions of rs79716587 and rs140397066 indicated. Data from Calderon *et al.* **(D)** Allelic series at *ARHGAP15* showing effects of rs79716587-A (common) and rs140397066-G (rare) on IgA levels.

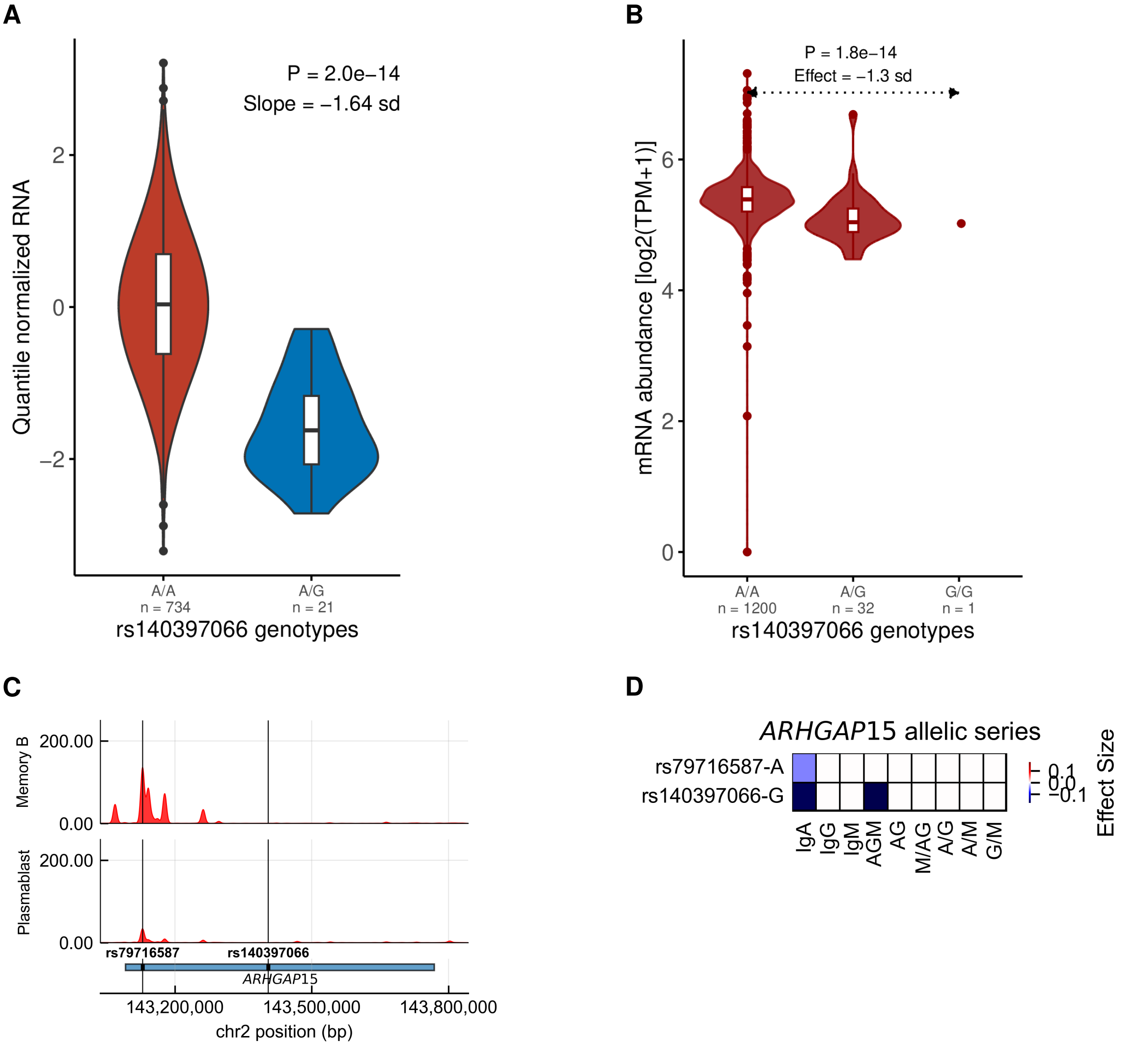

### Figure S9

Selected examples of candidate molecular mechanisms among the 24 independent signals at the polymorphic Fc receptor locus on chromosome 1. **(A)** Expression of *FCGR2B* by rs143596860 genotype across immune cell subsets. The AGM-increasing A allele decreases *FCGR2B* expression, most strongly in mDCs, consistent with de-repression of antigen-presenting cells. This signal also colocalizes with an eQTL for *FCRLB* in mDCs, though the association with *FCGR2B* expression is substantially stronger. **(B)** Expression of *FCRLB* by rs12139524 genotype across immune cell subsets. *FCRLB* remains poorly characterized, with no established role in immunoglobulin regulation. Here, the AG-decreasing C allele downregulates *FCRLB* in memory B cell subsets, particularly unswitched memory and double-negative B cells, providing evidence that reduced *FCRLB* expression in memory B cells leads to decreased class-switched immunoglobulin production. **(C)** Domain structure of FCGR2B (UniProt:P31994), showing the two extracellular immunoglobulin-like C2-type domains (IgC2-1, IgC2-2), transmembrane domain (TM), and immunoreceptor tyrosine-based inhibitory motif (ITIM). The rare missense variant rs149249317 (p.Pro47Ser) is classified as benign by computational predictors (CADD = 0.127, AlphaMissense = 0.085), likely because P47 falls just outside the annotated IgC2-type domain boundary. However, proline residues are systematically enriched at interdomain linker regions in multi-domain proteins, and P47 occupies precisely such a position, in the hinge immediately preceding the first IgC2-type domain of the FCGR2B ectodomain. **(D)** Allelic series showing effects of rs143596860-A, rs12139524-C, and rs149249317-T (p.Pro47Ser) on immunoglobulin traits. The strong increase in IgG associated with P47S is consistent with loss of FCGR2B inhibitory function. eQTL data from ImmuNexUT.

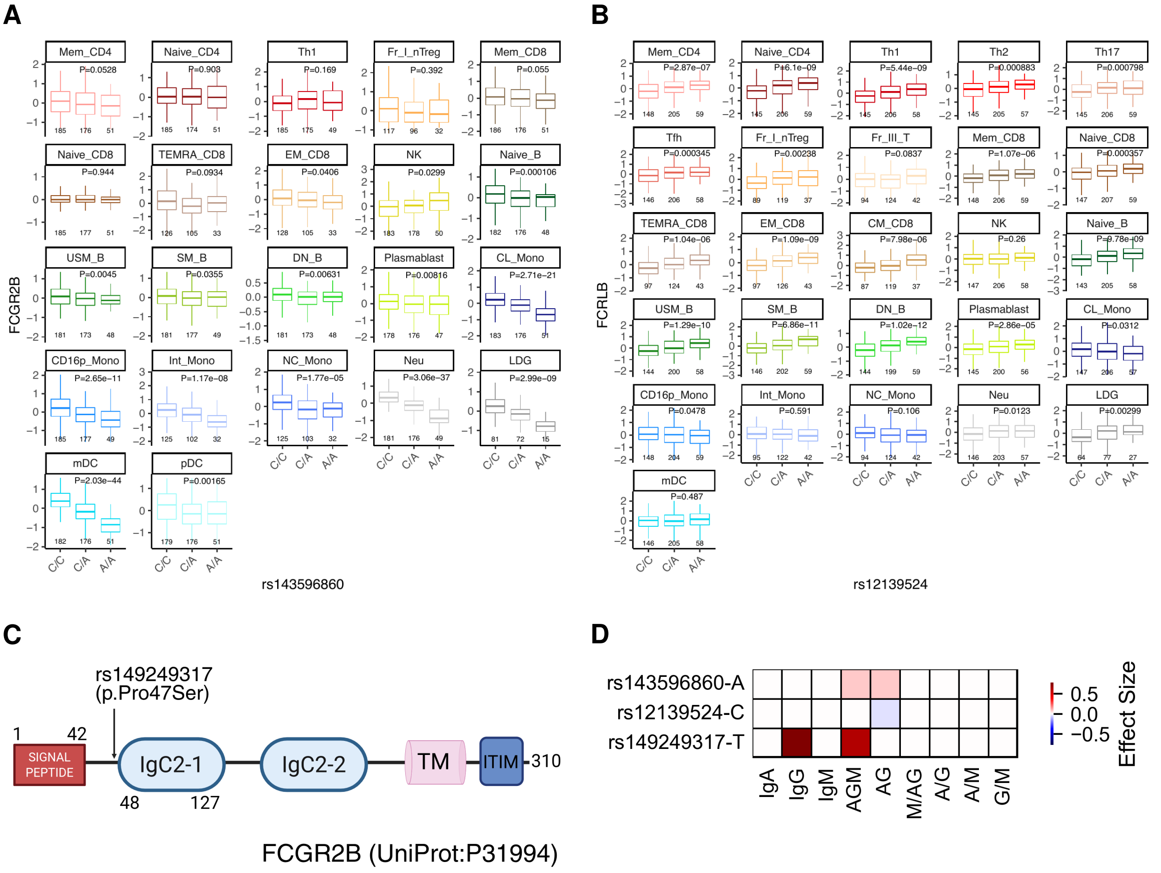

### Figure S10

**(A)** Expression of *TNFRSF13B* in whole blood by rs577044977 genotype across 17,695 Icelanders. $y$-axes indicate normalized gene expression; $x$-axes indicate genotype with sample sizes for each genotype. **(B,C)** Expression of *TNFSF13* by rs9907657 and rs3803800 genotype across immune cell subsets (data from ImmuNexUT; PMID 33930287).

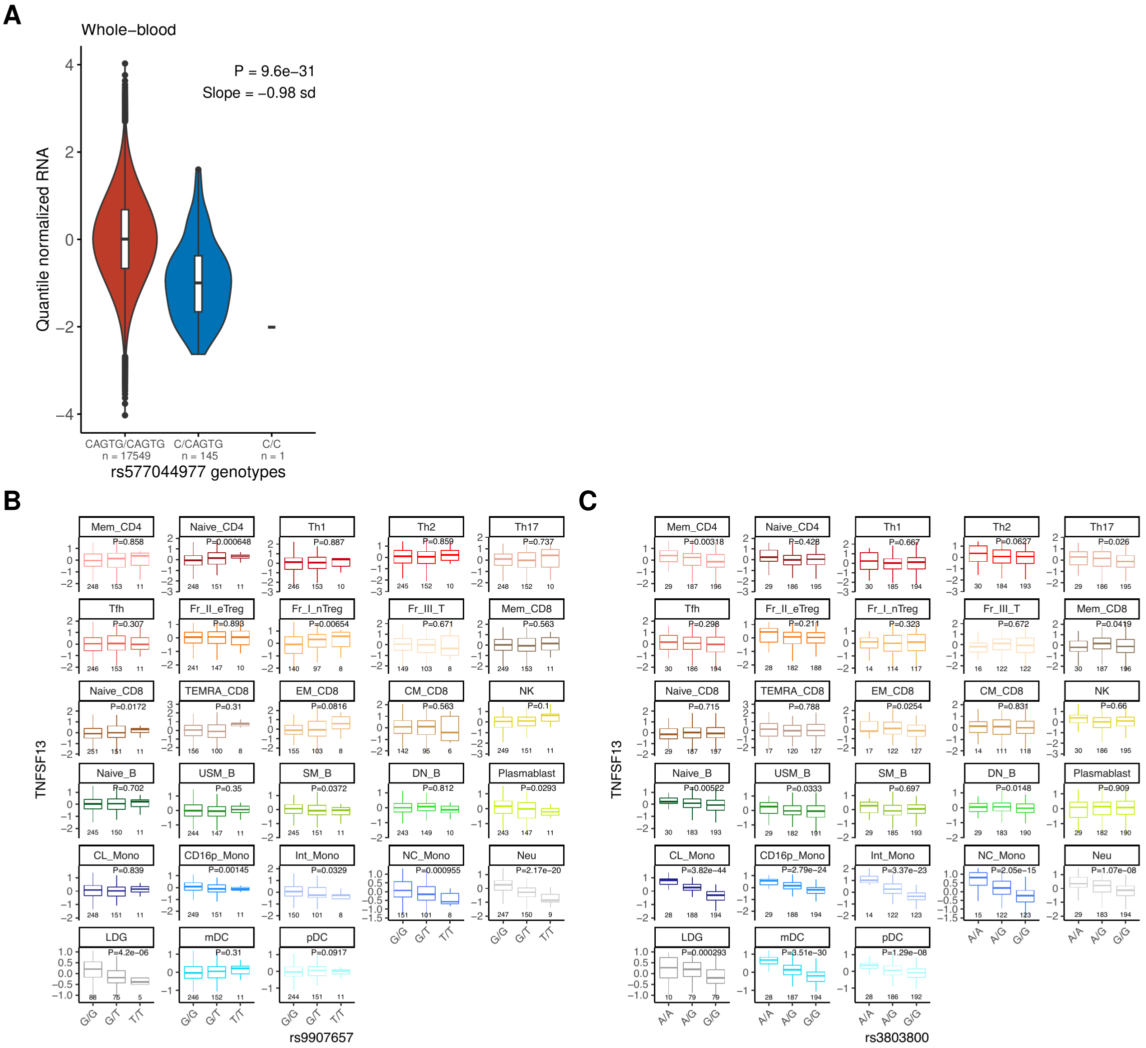

### Figure S11

siRNA dose-response for *EGR1* knockdown in MOLM13 cells. *EGR1* mRNA levels were measured by qPCR normalized to *ACTB* and expressed as percentage of the untreated control (0 nM). The 50 nM dose (red bar) was selected as the optimal concentration for subsequent experiments, achieving 46.5% knockdown. Error bars indicate 95% confidence intervals from technical triplicates.

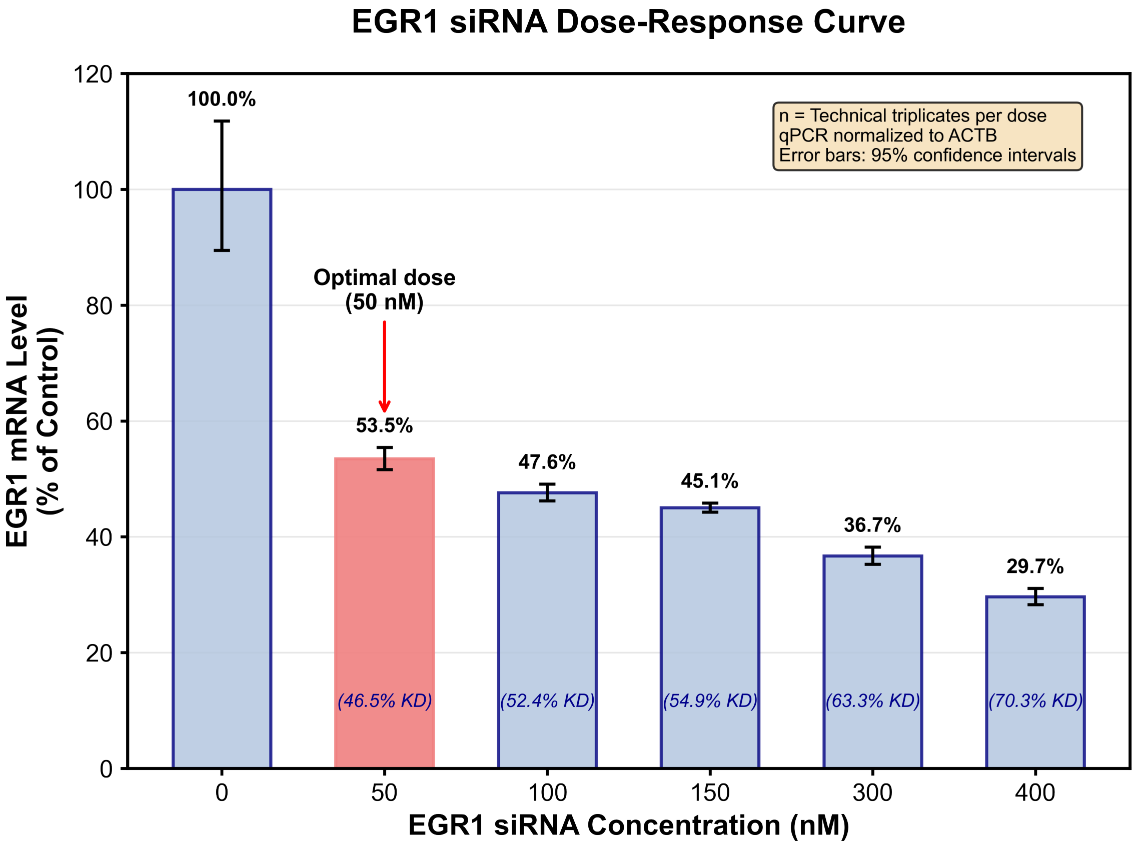
